# An AI System for Autonomous Algorithm Evolution in Drug Development

**DOI:** 10.64898/2026.08.16.745117

**Authors:** Zhimeng Zhou, Yang Nan, Minjie Mou, Yuntao Qian, Yixiu Liu, Zhengyu Zuo, Hao Yang, Weixian Xu, Bo Li, Wanghao Jiang, Yanlin Ren, Yang Liao, Yimeng Wang, Yinghong Li, Qingxia Yang, Zhiheng Xi, Tiantian Mi, Huaicheng Sun, Pengfei Liu, Feng Zhu

**Affiliations:** School of Pharmacy, Innovation Institute for Artificial Intelligence in Medicine of Zhejiang University, National Key Laboratory of Advanced Drug Delivery and Release Systems, Zhejiang University, Hangzhou 310058, China; Qing Yuan Research Institute, School of Computer Science, Shanghai Jiao Tong University, Shanghai 200240, China; Shanghai Innovation Institute, Shanghai 200237, China; College of Computer Science and Artificial Intelligence, Fudan University, Shanghai 200437, China; Department of Pharmacy, The Second Affiliated Hospital, Zhejiang University School of Medicine, Hangzhou 310009, China; School of Pharmacy, Hebei Medical University, Shijiazhuang 050017, China; Wisdom Lake Academy of Pharmacy, Xi’an Jiaotong-Liverpool University, Suzhou 215123, China

**Author notes:** Corresponding Authors: Prof. Feng Zhu, Prof. Pengfei Liu, Dr. Minjie Mou. These authors contributed equally to this work (Zhimeng Zhou, Yang Nan, Minjie Mou).

**Keywords:** Drug development, Artificial intelligence system, Target identification, Autonomous algorithm evolution, Large language models

## Abstract

Artificial intelligence (AI) is increasingly permeating the drug development pipeline. Numerous algorithms for accelerating this multi-stage and multi-task process have been constructed, which depends heavily on expert design and labor-intensive task-specific optimization. Given that AI-driven acceleration of drug development is recognized as a cumulative, often synergistic, effect across multiple stages, the autonomous evolution of existing algorithms across the entire pipeline is demanded to achieve a holistic advancement. Here, we present DrugEvolve, a multi-role large language model system for systematic and autonomous algorithm evolution in drug development. DrugEvolve realizes a closed-loop evolution process by incorporating *Researcher*, *Engineer*, and *Analyst* domains, and enables an iterative design, implementation, evaluation, and refinement of algorithm by leveraging scientific knowledge and accumulated evolutionary experience. Across eleven representative tasks spanning target identification, drug discovery, preclinical study, and clinical trial, DrugEvolve autonomously evolved the corresponding task-specific algorithms and achieved substantial performance enhancement on 120 benchmark test sets. Moreover, it showed robust generalizabilities across heterogeneous data modalities (ranging from biological sequence and graph to molecular topology and textual language), and realized gains in both predictive and generative tasks. Collectively, this AI system can serve not only as an algorithmic infrastructure for drug development, but also as a transferable paradigm for broader scientific domains.

## 1. Introduction

Most recently, artificial intelligence (AI) systems based on large language models (LLMs) have demonstrated remarkable capabilities in automating and accelerating scientific discovery^1–8^. Despite this progress, the broader scientific workflow is frequently bottlenecked by the inherently manual and iterative creation of computational models for subsequent experimentation^9,10^. This bottleneck stems from conventional algorithm design paradigms, which rely heavily on human intuition or expediency over exhaustive validation^11,12^. The resulting protracted development cycles severely limit large-scale algorithm discovery and the scope of viable research avenues^13^.

This bottleneck is particularly severe in drug development^14^. Drug development is a lengthy and costly process^15^ involving multiple complex stages (from target identification, to drug discovery, to preclinical, and to clinical research) and heterogeneous tasks, each requiring corresponding domain-specific computational tools for acceleration^16,17^. The inherent complexity of this field makes designing efficient algorithms both difficult and time-consuming, necessitating the integration of extensive interdisciplinary knowledge, including but not limited to cell biology, molecular biology, pharmacology, medicinal chemistry, and computer science^18,19^. Distilling effective algorithmic design principles from a tremendous and exponentially growing knowledge base is a long and challenging trial-and-error process, typically requiring tedious work often spanning many years^9,20^. The Nobel Prize-winning project AlphaFold serves as a prime example: despite DeepMind committing massive human and computational resources, its evolution from version 1.0 to 3.0 still took nearly six years (spanning from 2018 to 2024)^21–23^. This fully reflects the resource-intensive nature of algorithm development, the core bottleneck of which lies in the need for human experts to repeatedly execute the closed loop of “model design, training & tuning, evaluation, and development” (**Figure 1a**).

**Figure 1.**
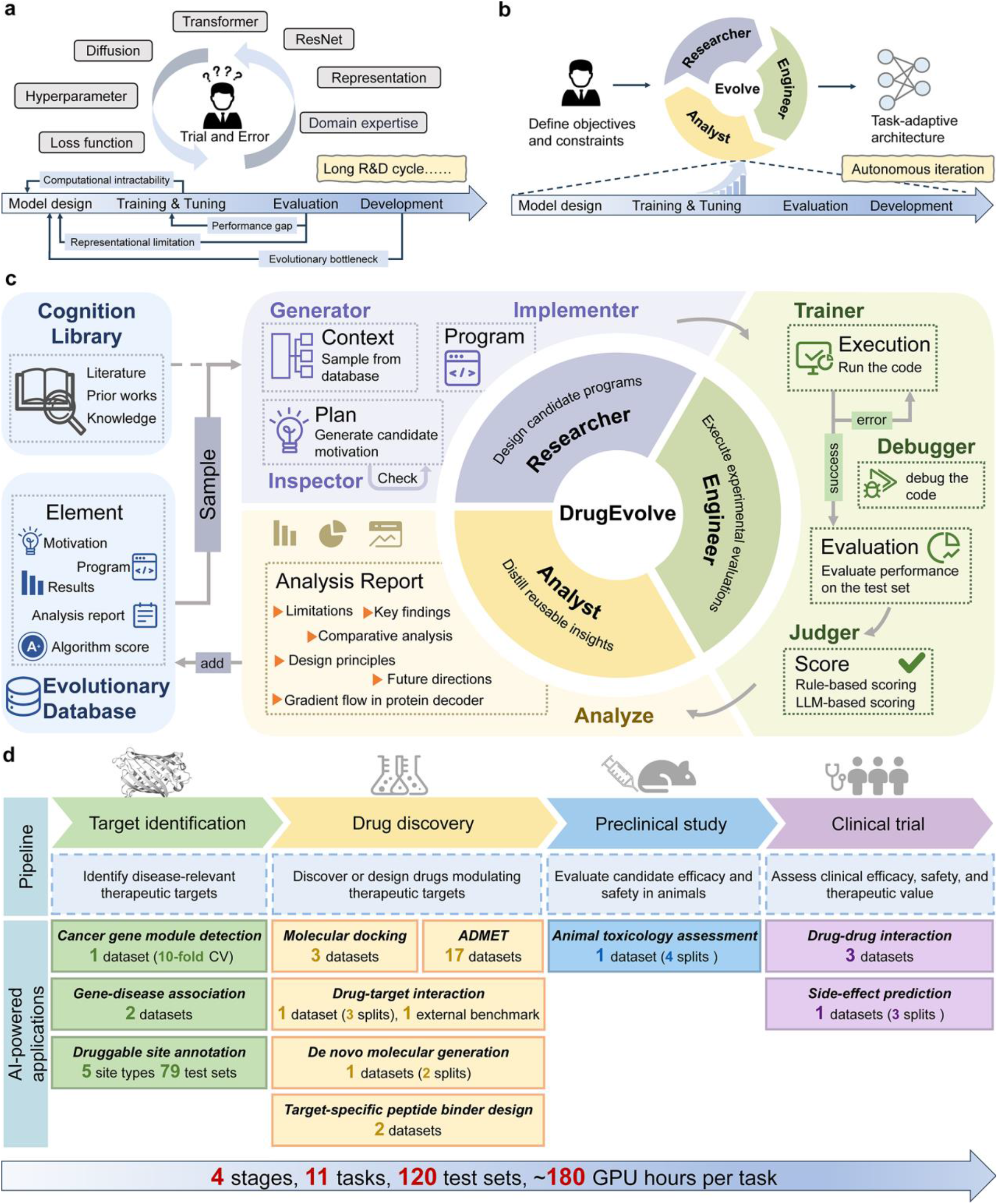
DrugEvolve enables autonomous algorithm evolution for drug development. **a.** Conventional AI algorithm development requires repeated human-led cycles of model design, implementation, training, evaluation, and refinement. **b.** Human scientists define the task objectives and constraints, while DrugEvolve autonomously executes the complete optimization loop, substantially accelerating algorithm development. **c.** Overall framework of DrugEvolve. DrugEvolve integrates multi-role LLM agents across the *Researcher*, *Engineer*, and *Analyst* domains to enable autonomous algorithm evolution. The framework combines scientific knowledge with accumulated evolutionary experience to iteratively discover and optimize task-specific algorithms. **d.** Overview of DrugEvolve applications across the drug development pipeline. DrugEvolve supports diverse tasks spanning four major stages of drug development, including target identification, drug discovery, preclinical study, and clinical trial. Numbers within each task box indicate the corresponding benchmark datasets and evaluation splits.

Although LLM-based coding agents such as Codex^24^ and AlphaCode^25^ have demonstrated the potential to automate algorithm design based on prompts, they often suffer from the limitation of “one-shot generation”^26^ and lack systematic iterative optimization capabilities. In contrast, the recently proposed ERA framework can assist scientists in designing computational tools, possesses the ability to build machine learning models from scratch, and supports continuous model iteration^27^. Similarly, AI systems such as AlphaEvolve^28^, CodeEvolve^29^, MLEvolve^30^, and AIBuildAI^31^ have successfully achieved autonomous algorithm evolution in specific “AI for AI” domains, including signal processing, computer vision, and natural language processing. The essence of autonomous algorithm evolution lies in sustained optimization and summarization capabilities, which are becoming central to automated machine learning algorithm discovery^30,32^.

Given that drug development represents a highly complex, multi-stage, and multi-task process, numerous advanced algorithms integrating extensive domain expertise and empirical knowledge have been successfully deployed across virtually all representative tasks^23,33–37^. Conducting autonomous evolution based on these powerful empirical algorithms, essentially “standing on the shoulders of giants”, represents a more efficient approach for convergence, and the demand for such application scenarios is substantial. Furthermore, the acceleration of drug development by AI is recognized as a cumulative, and often synergistic, effect across multiple stages^38–40^. In other words, merely optimizing one specific task is inadequate to achieve meaningful efficiency gains; rather, a holistic acceleration across the entire drug development pipeline is essential. However, none of the existing systems have attempted to achieve autonomous algorithm evolution across the entire drug development pipeline (spanning from target identification to clinical research^16^), let alone explore the extent to which AI systems can enhance the performance of these algorithms for drug development.

In this study, we introduce DrugEvolve, an AI system designed for autonomous algorithm evolution in drug development. As shown in **Figure 1b**, DrugEvolve leverages a specialized multi-role LLM framework (including *Researcher*, *Engineer*, and *Analyst*) alongside advanced sampling algorithms to execute systematic and sustained algorithmic evolution. Specifically, for distinct tasks across each drug development stage, the system autonomously executes a closed-loop cycle of problem parsing, algorithm design, model debugging, result analysis, reflective feedback, and iterative optimization. Starting from foundational algorithm designs established by human scientists, DrugEvolve can systematically evolve these architectures, thereby driving performance enhancements without manual intervention. To validate the intelligence and practicality of DrugEvolve, we selected 11 representative tasks and 120 benchmark test sets spanning the four stages of drug development: target identification, drug discovery, preclinical study, and clinical trial. Our experiments show that DrugEvolve successfully achieves autonomous algorithm evolution and remarkable performance improvements across nearly all tested tasks. This work represents a pioneering step towards fully AI-driven algorithmic evolution in drug development. Given the vast array of remaining tasks and specialized expert models in this field, the potential application scenarios for DrugEvolve are extensive. All in all, DrugEvolve serves not only as an algorithmic infrastructure for AI-driven drug development, but also as a transferable paradigm for autonomous algorithm evolution, accelerating the algorithmic innovation across broader scientific domains.

## 2. Results

### 2.1 The framework of DrugEvolve

DrugEvolve integrates multi-role LLM agents into a closed-loop framework for autonomous algorithm evolution in drug development (**Figure 1c**). Starting from a baseline algorithm for a specific task, along with task-specific datasets and predefined evaluation criteria and objectives provided by human scientists, the system automatically and iteratively performs algorithm design, implementation, evaluation, and optimization across three interconnected domains: *Researcher*, *Engineer*, and *Analyst*. The framework is motivated by the observation that effective algorithm development typically benefits from the integration of scientific knowledge, accumulated experience, and exploration of multiple candidate solutions. Accordingly, the three domains are designed to emulate these complementary aspects of human research.

Within the *Researcher* domain, candidate algorithms are proposed by integrating the scientific knowledge from *Cognition Library* with the prior evolutionary experience from *Evolutionary Database*. These candidates are then evaluated for conceptual novelty and scientific validity before being translated into executable implementations. The *Engineer* domain is responsible for algorithm implementation, debugging, model training, and performance evaluation. Candidate algorithms are assessed using task-specific quantitative metrics, with optional LLM-based qualitative evaluation providing complementary insights into the overall characteristics of selected evolved solutions. Following evaluation, the workflow advances to the *Analyst* domain, where evolutionary outcomes are systematically examined to identify factors associated with success or failure and to infer promising directions for subsequent iterations. The resulting insights are stored in the *Evolutionary Database* and reused in future iterations, thereby enabling continual accumulation of evolutionary knowledge. Coupled with various sampling strategies that balance exploration and exploitation during historical knowledge retrieval, these components empower DrugEvolve to achieve transparent, memory-driven, and continuously improving algorithm evolution.

### 2.2 Application overview of DrugEvolve in drug development

Previous studies have described drug development as a multistage process encompassing target identification, drug discovery, preclinical research, and clinical trial^16,41^. In this study, to address the algorithmic bottlenecks constraining progress across this continuum, we leveraged DrugEvolve on 11 representative tasks spanning the four main stages of drug development, establishing a robust computational paradigm designed to empower successive phases of the drug discovery pipeline (**Figure 1d**). These tasks align with the critical decision milestones in drug development, systematically covering the associated computational challenges. Specifically, at the *Target Identification* stage, cancer gene module detection extracts disease-relevant signals from multi-omics networks, gene-disease association prediction links candidate genes to specific diseases, and protein druggable site annotation identifies residue-level regions for therapeutic modulation of prioritized targets. In the mainstream paradigm of target-based *Drug Discovery*, once potential targets are identified, molecular docking and drug-target interaction prediction methods effectively accelerate the screening of bioactive molecules^42,43^. Additionally, de novo molecular generation and target-specific peptide binder design facilitate the exploration of novel chemical spaces. Subsequently, the prediction of ADMET (absorption, distribution, metabolism, excretion, and toxicity) properties supports the prioritization of candidates with favorable pharmacokinetic and safety profiles. Animal toxicology assessment further extends this process to *Preclinical Study* for evaluating drug safety, while drug-drug interaction and side-effect prediction identify potential risks in *Clinical Trial* prior to drugs’ clinical use.

In summary, the drug development pipeline comprises a multitude of intricate and successive stages. Since each stage involves heterogeneous data modalities (ranging from biological sequence and graph to molecular topology and textual language), methodological approaches, and prediction objectives, it necessitates highly specialized AI-driven computational approaches. However, the inherent limitations of existing approaches present a pressing need for autonomous algorithm evolution. Addressing this challenge, DrugEvolve successfully conducted autonomous algorithm evolution across nearly all evaluated tasks, underscoring its remarkable versatility. Despite substantial variations in task complexity, baseline architectures, optimization objectives, and computational costs, the system improved algorithmic performance across 11 tasks and 120 benchmark test sets. Notably, excluding the druggable site annotation task used for evolutionary trajectory analysis, the average computing cost for each task was approximately 180 GPU hours (**Supplementary Table S1**), with some requiring only a few GPU hours. This demonstrates our system’s broad applicability and practical executability, and indicates that the evolved models in this study have not yet reached their evolutionary limits, leaving room for future improvement.

### 2.3 Benchmarking autonomous algorithm evolution across diverse tasks

#### 2.3.1 Target Identification

##### DrugEvolve improves multi-omics modeling for cancer gene module detection

The advent of high-throughput and multi-omics sequencing technologies has established gene module detection as a cornerstone for the large-scale interpretation of biological data^44^. By elucidating the coordinated biological pathways underlying disease pathogenesis, this approach provides a systems-level foundation for downstream therapeutic target discovery and drug development^45^. To improve the integration of multi-omics information for cancer gene module detection, DrugEvolve adopted the CGMega^46^ model as the evolutionary starting point. The resulting CGMega-Evo preserves the original Transformer backbone but is augmented with a structural representation branch and a residual correction strategy, empowering it to capture higher-order network structural patterns beyond the reach of conventional multi-omics feature aggregation (**Supplementary Figure S1a**).

On the MCF-7 breast cancer multi-omics dataset^46^, CGMega-Evo improved cancer gene prediction under the fixed test evaluation protocol. Following the 10-fold cross-validation strategy, the same held-out test set was evaluated using ensemble predictions from models trained with different training-validation partitions. CGMega-Evo achieved superior overall performance compared to the baseline CGMega, with the MCC and F1 score increasing from 0.769 to 0.809 and from 0.812 to 0.844, respectively (**Supplementary Table S2**). Furthermore, CGMega-Evo demonstrated marginally higher performance across three overall evaluation metrics (AUROC, AUPRC, and ACC). These improvements indicate that CGMega-Evo provides more reliable cancer gene prediction under imbalanced classification settings. Beyond overall predictive performance, although a slight decrease in recall was observed, CGMega-Evo demonstrated substantially improved precision and specificity. This indicates a more accurate identification of cancer-associated genes and a more effective exclusion of background genes. Furthermore, across the ten training rounds, CGMega-Evo consistently outperformed CGMega on the majority of metrics, supporting the robustness of the evolved architecture under varying training-validation configurations (**Supplementary Figure S1b**). Together, these findings indicate the capacity of DrugEvolve to advance multi-omics modeling in complex biological systems and support the prioritization of new therapeutic targets.

##### DrugEvolve improves gene-disease association prediction

The prediction of gene-disease association (GDA) is a vital task in early-stage drug development, aiming to identify disease-associated genes and prioritize therapeutic targets^47^. To assess whether DrugEvolve can autonomously discover more effective architectures for GDA prediction, we applied it to the FusionGDA model^48^, yielding the evolved FusionGDA-Evo. Compared with FusionGDA, FusionGDA-Evo replaces the original self-attention and cross-attention fusion module with a physics-inspired Hamiltonian interaction module. As shown in **Supplementary Figure S2a**, it formulates gene and disease representations as interacting dynamic systems and performs feature integration via energy-driven message passing and dynamical state updates.

To assess the robustness across different data partitions, we evaluated both models using five independent random splits of the DisGeNET-EVAL benchmark^49^. FusionGDA-Evo demonstrated superior performance over FusionGDA across all splits. Specifically, it raised the mean test AUROC from 0.901 to 0.971 and AUPRC from 0.910 to 0.971, while achieving average improvements of 0.093, 0.096, and 0.096 in Accuracy, F1, and Fmax, respectively (**Supplementary Figure S2b**). The consistent improvements across independent partitions indicate that the evolved architecture yields stable performance gains, rather than overfitting to specific data splits. Furthermore, we evaluated FusionGDA-Evo on the external TDC dataset^50^ to assess its generalizability beyond the DisGeNET-EVAL benchmark. As illustrated in **Supplementary Figure S2c**, FusionGDA-Evo improved across all evaluation metrics, consistently outperforming FusionGDA by margins exceeding 0.070, thereby demonstrating robust generalization across different datasets.

##### DrugEvolve enhances broad annotation of protein druggable sites

Residue-level annotation of protein druggable sites is essential for understanding protein function and guiding downstream drug discovery^51,52^. To evaluate generalization capabilities across diverse site identification tasks, the recently published sequence-based framework, ALLSites, was selected as the evolutionary object for DrugEvolve, utilizing six datasets representing key modalities of non-covalent binding sites^53^, including DNA-protein (DPI), RNA-protein (RPI), protein-protein (PPI), peptide-protein (PepPI), small molecule-protein (SMPI), and carbohydrate-protein interactions (CarbPI). The highest-fitness model obtained after evolution was subsequently comprehensively evaluated on 79 independent test datasets spanning five major categories of druggable site annotation tasks, including non-covalent binding sites, covalent binding sites, catalytic sites, cryptic sites, and post-translational modification (PTM) sites (**Figure 2a**). Compared with ALLSites, the evolved architecture (ALLSites-Evo) substantially restructures the original framework by introducing a multi-scale gated CNN for feature extraction and replacing the original Transformer decoder with an attention-pooling module for global-local feature integration (**Figure 2b**).

**Figure 2.**
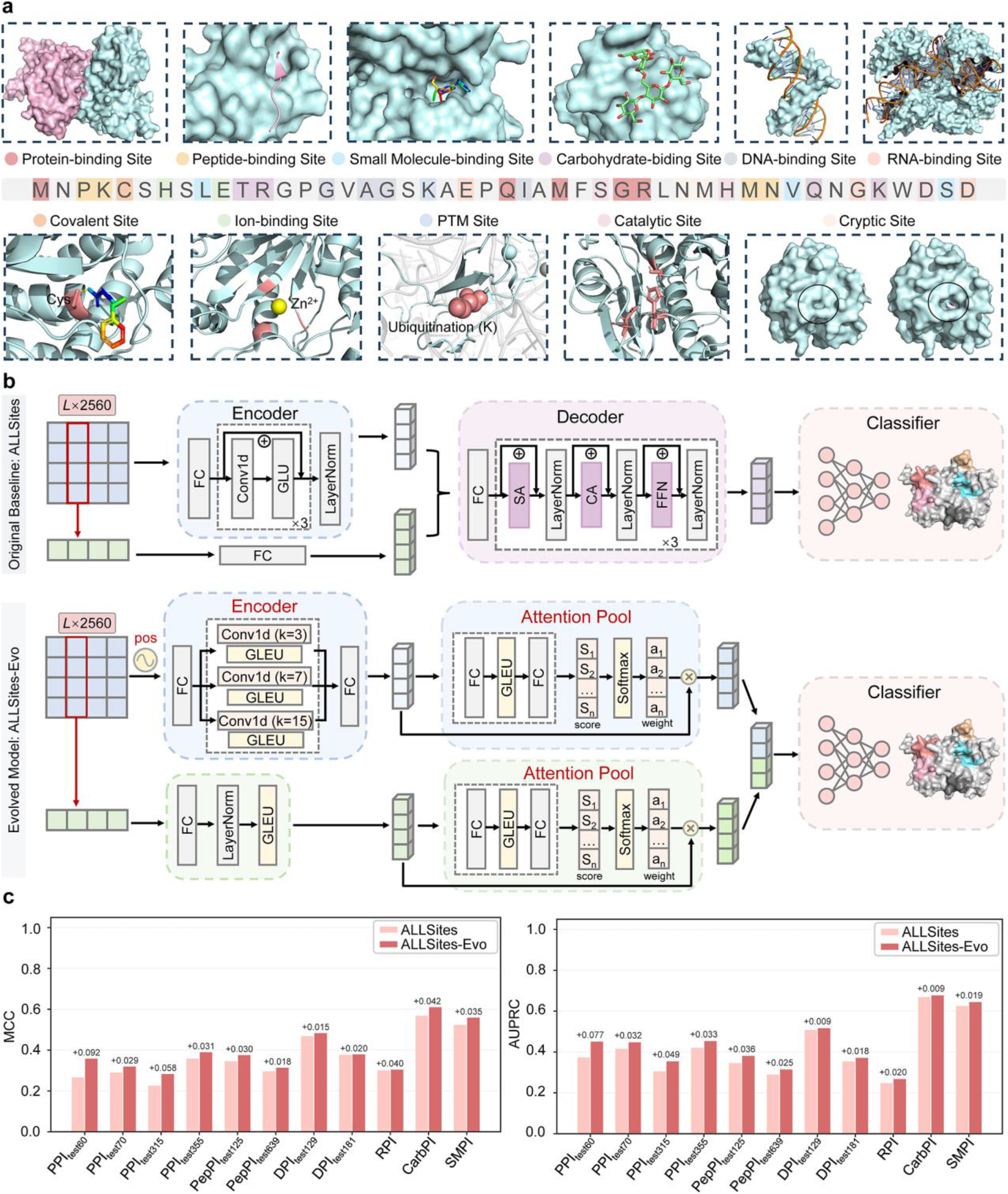
Autonomous evolution of ALLSites for protein druggable site annotation. **a.** Overview of the druggable residue types evaluated in this study, comprising six binding site types optimized during algorithm evolution (protein-, peptide-, small molecule-, carbohydrate-, DNA-, and RNA-binding sites) and five unseen site types for evaluating generalization (covalent-, ion-binding, post-translational modification, catalytic, and cryptic sites). **b.** Comparison of original ALLSites and evolved ALLSites-Evo. ALLSites-Evo introduces multi-scale feature extraction and parallel attention-based feature fusion. **c.** Performance comparison of ALLSites and ALLSites-Evo across 11 interaction-site benchmarks measured by MCC and AUPRC.

Across non-covalent binding-site prediction tasks, ALLSites-Evo consistently outperformed ALLSites across all 11 interaction-site benchmarks including the six datasets used for evolutionary optimization, while maintaining competitive performance against specialized methods (**Figure 2c, Supplementary Figure S3, and Supplementary Tables S3 and S4**).

Notably, these gains generalized to unseen non-covalent binding-site categories, yielding average MCC improvements of more than 0.075 across the 19 heterogeneous ligand and ion types in LigBind, as well as 0.074 across six representative classes in GPSites (including ATP, HEM, Zn²⁺, Ca²⁺, Mg²⁺, and Mn²⁺) (**Figure 3a and Supplementary Figure S4a,b**). Generalization examination further extended beyond non-covalent interactions to diverse site types not encountered during evolutionary optimization. Specifically, on the CovalentInDB dataset^54^, covalent binding-site prediction improved from an AUPRC of 0.713 to 0.823, covering nine types of covalent amino acid residues (**Figure 3b**). Notably, the model achieved consistent performance gains on cysteines, which represent the most common residue type (**Figure 3c**). This trend also extended to catalytic site prediction, where AUPRC increased from 0.649 to 0.719 on CataloDB, and the model consistently outperformed the specialized framework Squidly across CataloDB, Uni3175, and six additional independent test sets^55^ (**Figure 3d and Supplementary Figure S4c**). Improvements were also observed in cryptic site prediction^56^, with the AUROC increasing from 0.695 to 0.726 relative to the baseline, alongside concurrent enhancements in AUPRC and MCC (**Figure 3e**). The model’s robustness was further validated on PTM sites, with ALLSites-Evo achieving a mean AUROC improvement of 0.051 across eight benchmark sets spanning six PTM categories in PTMAtlas^57^ (**Supplementary Figure S5a**). Furthermore, it demonstrated exceptional stability under homology-controlled evaluations (at 90%, 80%, and 70% sequence identity thresholds), with performance variations confined to under 0.004 (**Supplementary Figure S5b**). These results indicat that ALLSites-Evo captures the transferable principles of residue-level functionality, rather than task-specific patterns.

**Figure 3.**
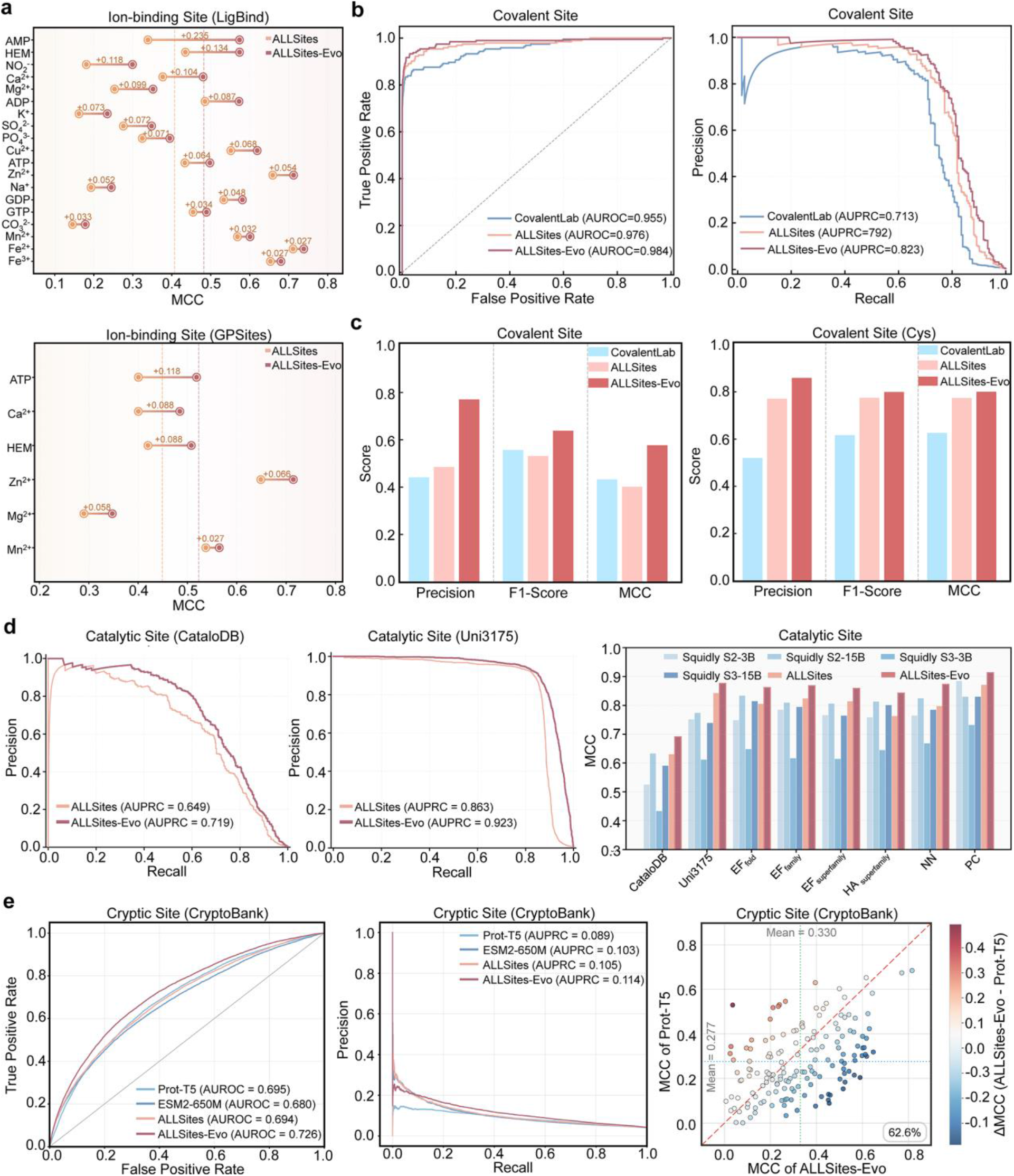
Generalization of ALLSites-Evo to unseen druggable site annotation tasks. **a.** Performance comparison of ALLSites and ALLSites-Evo on ion-binding site prediction across the LigBind and GPSites benchmarks, evaluated on multiple ion and endogenous ligand types using MCC. **b.** Performance comparison for covalent site prediction on the CovalentInDB benchmark, shown by AUROC and AUPRC. **c.** Performance comparison of CovalentLab, ALLSites, and ALLSites-Evo for covalent site annotation on the CovalentInDB and Cys benchmarks, evaluated by Precision, F1-score, and MCC. **d.** Performance comparison for catalytic site prediction on the CataloDB and Uni3175 datasets, together with reproduced Squidly variants under the original evaluation settings. The reproduced variants follow the Squidly framework and include Scheme 2 and Scheme 3 contrastive learning strategies with ESM2-3B and ESM2-15B backbones. Scheme 2 performs reaction-aware contrastive pair mining, whereas Scheme 3 introduces cross-class hard negative sampling. Scheme 1 was not reproduced because it was used as a comparative ablation setting in Squidly. **e.** Performance comparison of ALLSites and ALLSites-Evo for cryptic site annotation on the CryptoBank benchmark. Comparisons include fine-tuned ProtT5- and ESM2-650M-based models using AUROC, AUPRC, and protein-level MCC analyses.

Representative case studies further support these observations. Across enzymatic, covalent, and cryptic-site contexts, ALLSites-Evo recovered experimentally validated functional sites with high specificity. In catalytic site prediction, it effectively differentiated the catalytic lysozyme from its non-enzymatic homolog α-lactalbumin. Although they share highly similar structures, lysozyme and *α* -lactalbumin exhibit entirely different functions, and the model accurately identified two catalytic residues (GLU-35 and ASP-52) solely in the former (**Supplementary Figure S6a**). For covalent site prediction, it pinpointed the single experimentally validated reactive cysteine in an unseen protein (PDB: 5P9J), whereas CovalentLab failed to detect it (**Supplementary Figure S6b**). In cryptic site prediction, structural analysis of an apo-holo pair revealed that ALLSites-Evo recovered the vast majority of annotated cryptic-site residues, in contrast to competing methods that captured only partial regions (**Supplementary Figure S6c**). Together, these results demonstrate that ALLSites-Evo, developed by our DrugEvolve system, enables accurate and broadly transferable annotation of druggable sites on protein targets, establishing a robust foundation for protein characterization and AI-driven drug discovery.

#### 2.3.2 Drug Discovery

##### DrugEvolve drives more accurate molecular docking

Elucidating protein-ligand binding conformations is central to structure-based drug discovery, yet it remains challenging because molecular recognition relies on conformational flexibility and diverse physicochemical interactions^58,59^. Therefore, developing accurate docking methods remains one of the most challenging tasks in computer-aided drug discovery^60^. DrugEvolve evolved the well-established blind docking method, EquiBind^61^, to improve docking pose prediction. Compared to the original model, the evolved EquiBind-Evo focuses receptor keypoints within a ligand-conditioned binding region and further refines the Kabsch-aligned pose through local coordinate adjustments (**Figure 4a**).

**Figure 4.**
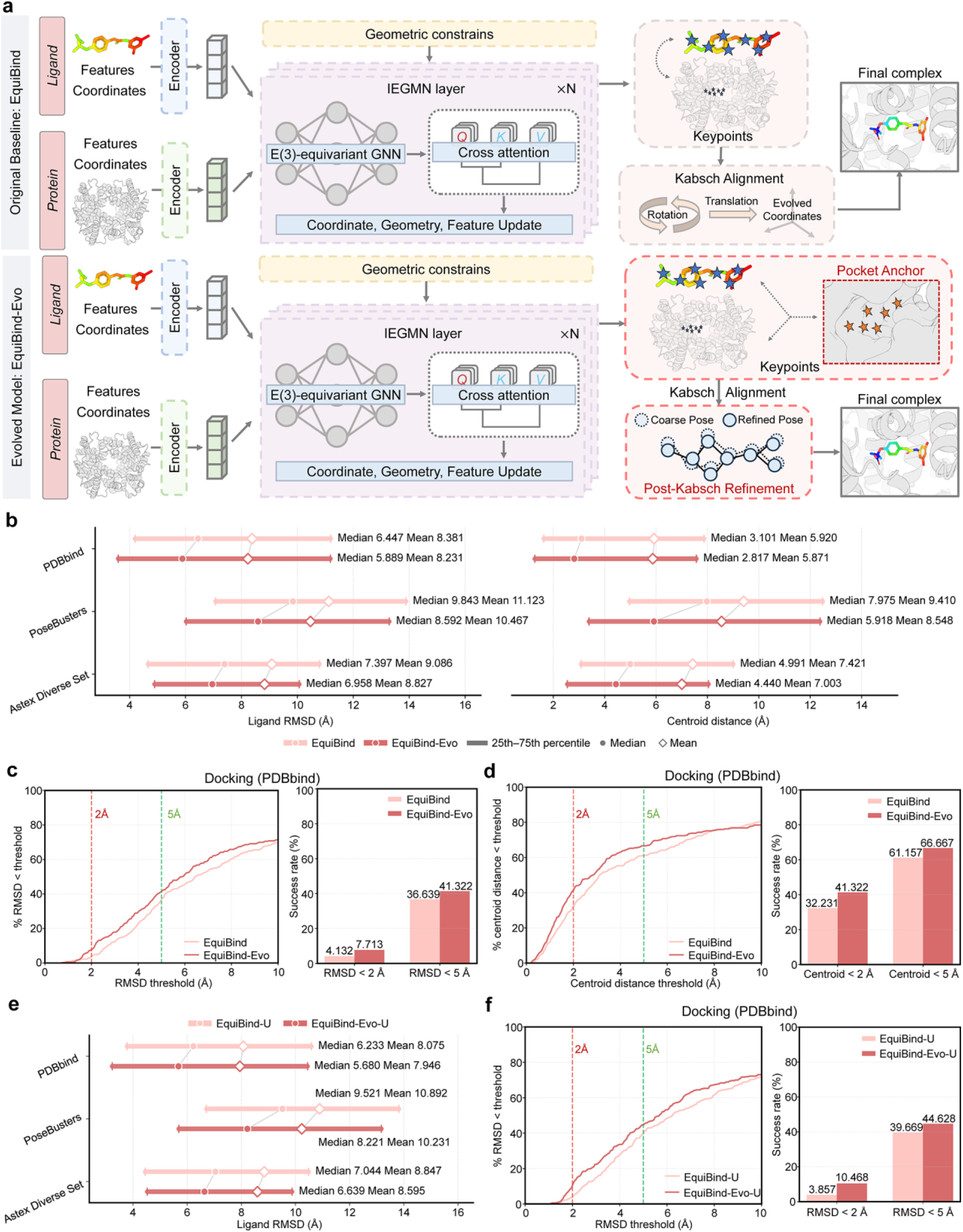
Autonomous evolution of EquiBind for molecular docking. **a.** Comparison of the original EquiBind and evolved EquiBind-Evo architectures. EquiBind-Evo improves blind docking by introducing ligand-conditioned pocket anchors and refining ligand poses through local coordinate adjustments. **b.** Performance comparison of EquiBind and EquiBind-Evo across the PDBbind, PoseBusters, and Astex Diverse Set benchmarks using ligand RMSD and centroid distance. Horizontal bars indicate the 25th-75th percentile range, central dots indicate medians, and diamonds indicate mean values. **c.** Success rate comparison of EquiBind and EquiBind-Evo under different ligand RMSD thresholds on the PDBbind test set. Curves show the percentage of predictions achieving RMSD values below each threshold, and bar plots summarize the success rates at the 2 Å and 5 Å thresholds. **d.** Success rate comparison of EquiBind and EquiBind-Evo under different centroid distance thresholds on the PDBbind test set. **e.** Ligand RMSD comparison between EquiBind-U and EquiBind-Evo-U across the PDBbind, PoseBusters, and Astex Diverse Set benchmarks. **f.** Success rate comparison of EquiBind-U and EquiBind-Evo-U under different ligand RMSD thresholds on the PDBbind test set.

We evaluated the evolved architecture both in its initial state (denoted EquiBind-Evo-U) and following fast conformer fitting (denoted EquiBind-Evo). On the temporally split PDBbind test set and two independent external benchmarks (PoseBusters and Astex Diverse Set), EquiBind-Evo consistently achieved lower ligand RMSD and centroid distance values than the original EquiBind in both mean and median statistics, indicating improved and robust prediction of ligand placement and binding-site localization (**Figure 4b**). We further analyzed the success rates across different RMSD and centroid distance thresholds on the PDBbind test set. Across the full threshold range from 0 to 10 Å, EquiBind-Evo consistently outperformed EquiBind for both metrics (**Figure 4c and Figure 4d**). Under the commonly used 5 Å threshold, EquiBind-Evo achieved higher success rates for both ligand placement and binding-site localization. Notably, under the more stringent 2 Å threshold, the proportion of ligand poses with RMSD below 2 Å increased from 4.1% to 7.7%, while the fraction with centroid distance below 2 Å increased from 32.2% to 41.3% (**Figure 4c and Figure 4d**). Similarly, the evaluation results on two additional external benchmarks demonstrated varying degrees of improvement. On the PoseBusters dataset, EquiBind-Evo exhibited improvements in both ligand placement and binding-site localization. In contrast, on the Astex Diverse Set, it demonstrated more moderate gains, primarily in centroid distance-based localization (**Supplementary Table S5**). Consistent improvements were also observed prior to conformer correction. Relative to EquiBind-U, EquiBind-Evo-U increased the proportion of sub-2-Å ligand poses from 3.9% to 10.5% on the PDBbind test set and nearly doubled the sub-5-Å success rate from 10.5% to 19.6% on PoseBusters (**Figure 4e and Figure 4f**). Regarding the Kabsch RMSD metric, performance remained broadly comparable, with modestly lower mean values observed across all three benchmarks (**Supplementary Table S5**).

These results indicate that the model evolved by DrugEvolve enhances the intrinsic geometric prediction quality prior to any refinement, rather than deriving performance gains from the conformer-fitting procedure.

Representative complexes selected exclusively from the independent test sets of the three benchmark datasets further illustrated the complementary geometric improvements learned by EquiBind-Evo. For the representative complex from the Astex Diverse Set (PDB ID: 1OPK; ligand: P16), EquiBind placed the ligand in an incorrect receptor region, whereas EquiBind-Evo successfully recovered the crystallographic binding pocket, thereby reducing the centroid distance from 14.045 Å to 1.755 Å (**Supplementary Figure S7a**). For another complex from PoseBusters (PDB ID: 7JXX; ligand: VP7), the Kabsch RMSD remained unchanged at 0.172 Å, while the ligand RMSD substantially decreased from 5.394 Å to 1.489 Å, indicating that EquiBind-Evo preserved the accurate local conformation of the ligand while substantially improving its global placement and orientation (**Supplementary Figure S7b**). In contrast, for the complex from the test set of PDBbind (PDB ID: 6K05; ligand: CQF), while both predictions correctly localized the ligand within the binding pocket, EquiBind-Evo reduced the Kabsch RMSD from 2.121 Å to 1.280 Å and the ligand RMSD from 5.537 Å to 1.313 Å (**Supplementary Figure S7c**). This demonstrates a marked improvement in recovering the internal ligand conformation. Collectively, these independent test cases highlight distinct sources of docking improvement achieved by the evolved model, spanning binding-pocket selection, rigid-body placement, and conformational refinement.

##### DrugEvolve enables generalizable prediction of drug-target interactions

The identification of drug-target interactions (DTI) serves as the foundation of modern target-based drug discovery^62,63^. However, the limited coverage of experimentally characterized drug-target interactions, together with the vast unexplored chemical and protein space encountered in practical applications, requires architectures capable of generalizing beyond the training distribution. To assess whether DrugEvolve can autonomously evolve architectures for this challenge, we applied it to DrugBAN^64^, yielding an evolved architecture termed DrugBAN-Evo. Compared with DrugBAN, DrugBAN-Evo differs primarily by introducing a hybrid bilinear interaction module that incorporates diagonal and low-rank interaction pathways, followed by Top-K sparse attention refinement (**Supplementary Figure S8a**).

On the BindingDB dataset, DrugBAN-Evo consistently outperformed DrugBAN across AUROC, AUPRC, F1, and MCC on the independent test set (**Supplementary Figure S8b**). These performance gains were particularly pronounced under cross-domain settings, demonstrating consistent improvements in both unseen-drug and unseen-protein test sets. Notably, the MCC for previously unseen protein targets increased from 0.472 to 0.555 (**Supplementary Figure S8b**). Under the more stringent Human-cold dataset, where neither test drugs nor test targets are seen during training, DrugBAN-Evo again achieved superior performance across all evaluation metrics, including a substantial MCC improvement of over 0.082 (**Supplementary Figure S8c**). The progressively larger gains observed under increasingly stringent evaluation settings suggest that DrugBAN-Evo learns transferable interaction representations rather than relying on memorized drug-target associations. Consequently, DrugEvolve can autonomously evolve DTI prediction architectures that generalize robustly beyond the training distribution.

##### DrugEvolve advances de novo molecular generation in unexplored chemical space

De novo molecular generation addresses a central challenge in drug discovery: efficiently navigating the vast chemical space to design novel, drug-like compounds with desired biological activities^65,66^. Within this paradigm, ligand-based molecular generation serves as a practical strategy, enabling generative models to learn from known active chemotypes and bias the design process toward target-relevant chemical space^67^. Among these methods, CharRNN has emerged as a highly effective and classical autoregressive model for de novo molecular generation ^68^. In this study, DrugEvolve further evolved the original architecture by redesigning its molecular representation and next-token prediction pipeline. Specifically, it introduces residual embedding propagation and combines a neural logit head, a low-rank adapter head, and a gated incorporation of a training-derived bigram prior to derive next-token logits (**Supplementary Figure S9a**).

Evaluation on the MOSES benchmark reveals that CharRNN-Evo not only better approximates the target molecular distribution but also substantially improves generalization to unseen scaffolds (**Supplementary Figure S9b**). While agreement with the reference distribution increased on both random test set (Test) and scaffold-based test set (TestSF), the scaffold split showed the largest margin of improvement, reducing FCD from 0.624 to 0.528. Although the Scaff metric decreased slightly on the random test set, it showed a doubling of similarity to the unseen-scaffold distribution on TestSF, demonstrating that the evolved model effectively captures scaffold patterns absent from the training data. Notably, novelty also increased markedly from 65.60% to 87.18%, indicating a broader exploration of chemical space beyond the training set, whereas internal diversity and uniqueness remained essentially unchanged. This demonstrates that the enhanced distribution learning and scaffold generalization were attained without a trade-off in molecular diversity or redundancy.

Scaffold-cluster analysis further elucidated the origin of this improvement. As shown in **Supplementary Figure S9c**, rather than uniformly increasing scaffold frequencies, CharRNN-Evo redistributed the generated molecules to better align with the scaffold composition of TestSF across most major scaffold families. Two representative examples illustrate the complementary mechanisms underlying this shift (**Supplementary Figure S9d**). In Case 1, CharRNN failed to generate molecules containing the displayed TestSF scaffold, which was successfully recovered by CharRNN-Evo, demonstrating the latter’s enhanced capability in expanding coverage to previously inaccessible scaffold families. In Case 2, although both models successfully generated new molecules containing the displayed scaffold, CharRNN-Evo produced substantially more representatives with this scaffold, yielding a scaffold distribution that more closely matched TestSF. Together, these examples indicate that the improved scaffold-split performance arises from both the recovery of previously uncovered scaffold families and a more faithful modeling of scaffold-family frequencies.

##### DrugEvolve enhances target-specific peptide binder design

Target-specific peptide binder design supports the development of selective therapeutic peptides against disease-relevant targets^37,69^. The newly proposed PepMLM formulates this task as target-conditioned masked language modeling (MLM), enabling de novo peptide generation from protein sequences without using any structural information^70^. However, the single-step reconstruction framework, instantiated via MLM, tends to bias generation toward recovering cognate binders, which may limit exploration of broader regions of peptide sequence space. Therefore, we targeted PepMLM for model optimization to evaluate whether DrugEvolve can recognize this limitation and achieve successful evolution. As a result, DrugEvolve discovered a substantially enhanced architecture, termed PepMLM-Evo, after only three evolutionary iterations. The evolved architecture replaces single-step peptide reconstruction with an iterative mask-based denoising framework under a cosine corruption schedule, combined with a receptor-peptide contrastive learning objective that induces interaction-aware representation alignment in a shared embedding space (**Figure 5a**). This enables progressive refinement of peptide sequences conditioned on protein context during generation.

**Figure 5.**
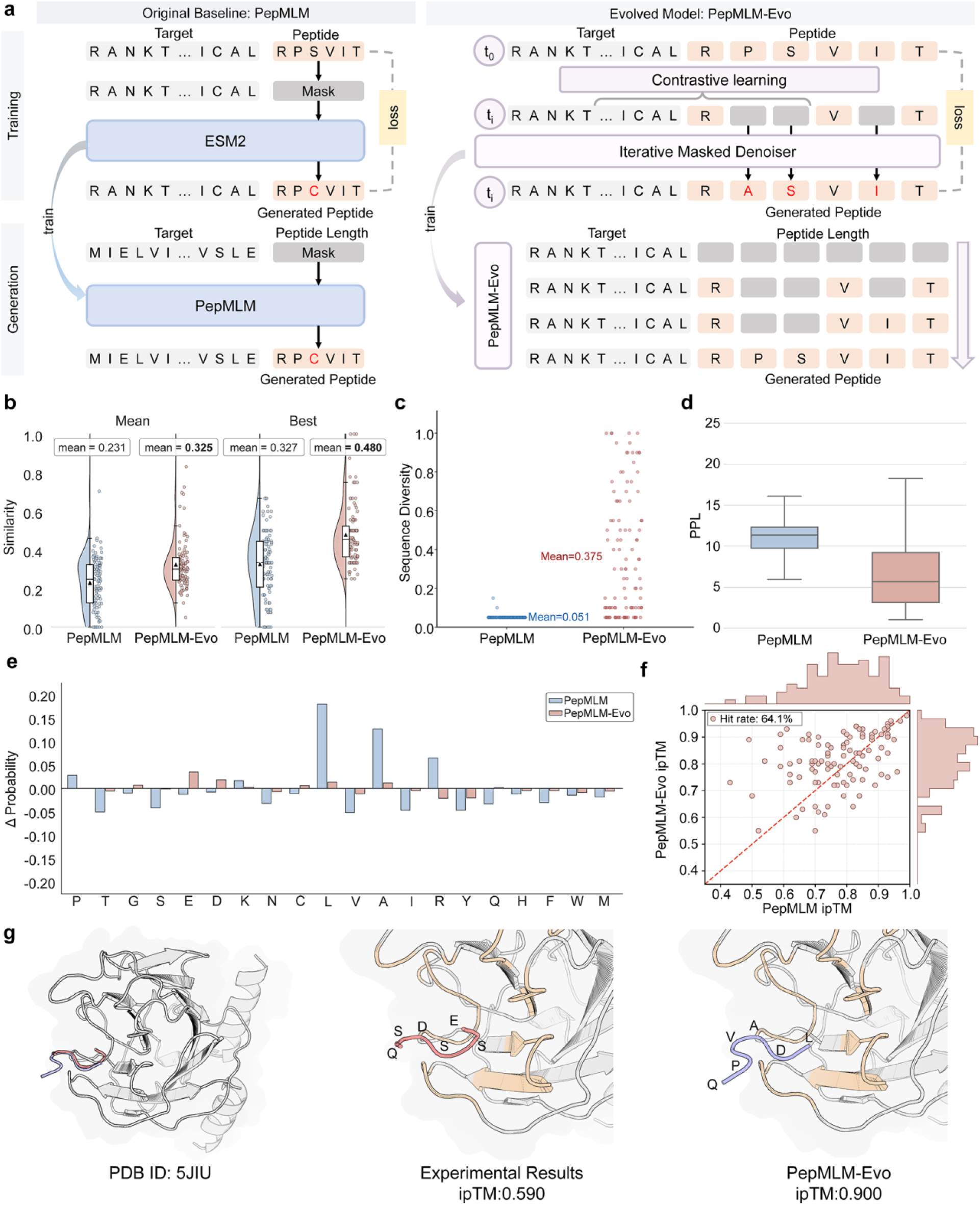
Autonomous evolution of PepMLM for target-specific peptide binder design. **a.** Comparison of the original PepMLM and evolved PepMLM-Evo architectures. PepMLM-Evo replaces single-step masked reconstruction with iterative mask refinement and introduces contrastive representation learning for protein-peptide pairs. **b.** Sequence similarity of peptides generated by PepMLM and PepMLM-Evo. The left two groups report the overall mean similarity, whereas the right two groups report the mean best similarity, calculated as the average of the highest similarity peptide for each target. Violin plots show the distribution of sequence similarity values, individual dots represent generated peptides, and black triangles indicate mean values. **c.** Sequence diversity of peptides generated by PepMLM and PepMLM-Evo. Each point represents the diversity of generated peptides for an individual target, and horizontal lines indicate the mean diversity across all targets. **d.** Peptide perplexity (PPL) comparison of peptides generated by PepMLM and PepMLM-Evo on the PepBench benchmark. Box plots summarize the PPL distribution across generated peptides. The central line indicates the median, the box represents the 25th-75th percentile range, and whiskers extend to values within 1.5 × the interquartile range. **e.** Differences in amino acid frequency distributions between designed peptides and the PepBench test set for PepMLM (blue) and PepMLM-Evo (red). Bars represent the difference between the amino acid frequency in generated peptides and that in the test set, calculated as generated frequency minus test-set frequency. **f.** In silico hit-rate evaluation of PepMLM and PepMLM-Evo based on ipTM scores predicted by AlphaFold3. Each point represents one target-peptide pair. Hits are defined as target-peptide pairs for which the peptide generated by PepMLM-Evo achieves an ipTM score equal to or higher than that generated by PepMLM. **g.** Structural comparison of an experimental peptide binder and a PepMLM-Evo-designed peptide in complex with the target protein (PDB ID: 5JIU). The target protein is shown in gray, the experimental peptide in red (ipTM = 0.590), and the PepMLM-Evo-designed peptide in purple (ipTM = 0.900). Contact residues within an 8 Å threshold are highlighted in yellow.

On the PepBench benchmark^71^, PepMLM-Evo consistently outperformed PepMLM across all major evaluation metrics. This improvement was reflected not only in increased similarity to native binders (mean: 0.231 to 0.325; best-case: 0.327 to 0.480, **Figure 5b**), but also in substantially enhanced sequence diversity (0.051 to 0.375, **Figure 5c**) and reduced perplexity (PPL) (**Figure 5d**). Consistent with these sequence-level improvements, amino acid composition analysis revealed that the peptides generated by PepMLM-Evo align more closely with experimentally validated binders, whereas those designed by PepMLM exhibited pronounced compositional biases (**Figure 5e**). Structural assessment using AlphaFold3^23^ indicated that these improvements extended beyond sequence-level metrics. Using the ipTM score as a proxy for binding confidence, PepMLM-Evo generated peptides with ipTM scores equal to or higher than those generated by PepMLM for 64.1% of targets (**Figure 5f**). Substantial performance improvements were also observed on the independent PepNN benchmark, further supporting the robustness and cross-dataset generalizability of the evolved architecture (**Supplementary Figure S10**). Notably, in a representative complex (PDB ID: 5JIU), AlphaFold3 predicted an ipTM score of 0.590 for the native peptide, whereas the peptide designed by PepMLM-Evo achieved a higher score of 0.900 (**Figure 5g**). Examination of the generated peptide sequences and predicted complex structures revealed that, despite substantial sequence divergence from the native binder, the designed peptide adopted a highly similar binding conformation and occupied the same interaction pocket. These findings suggest that the evolved PepMLM-Evo preserves key binding-recognition patterns while exploring broader regions of the peptide sequence space. Collectively, these results demonstrate that DrugEvolve efficiently evolves peptide-design architectures through rapid evolutionary search.

##### DrugEvolve improves generalizable prediction of ADMET properties

ADMET properties are central to the core objectives of lead optimization and efficacy evaluation, as well as chemical safety assessment^72,73^. They are widely recognized as key determinants of compound success during preclinical evaluation^74^. To systematically enhance predictive performance across diverse ADMET endpoints, DrugEvolve leverages the recently published FragNet^75^ model as the foundational architecture to discover more effective molecular representations. Building upon the original FragNet, FragNet-Evo substantially enhances hierarchical molecular representation learning through three key innovations: (1) fragment-guided atom feature refinement to strengthen cross-scale information exchange; (2) residual-gated normalization for stable feature propagation; and (3) multi-aggregation pooling to enrich graph-level representations. Altogether, these improvements yield highly expressive and robust molecular embeddings for downstream ADMET prediction (**Figure 6a**).

**Figure 6.**
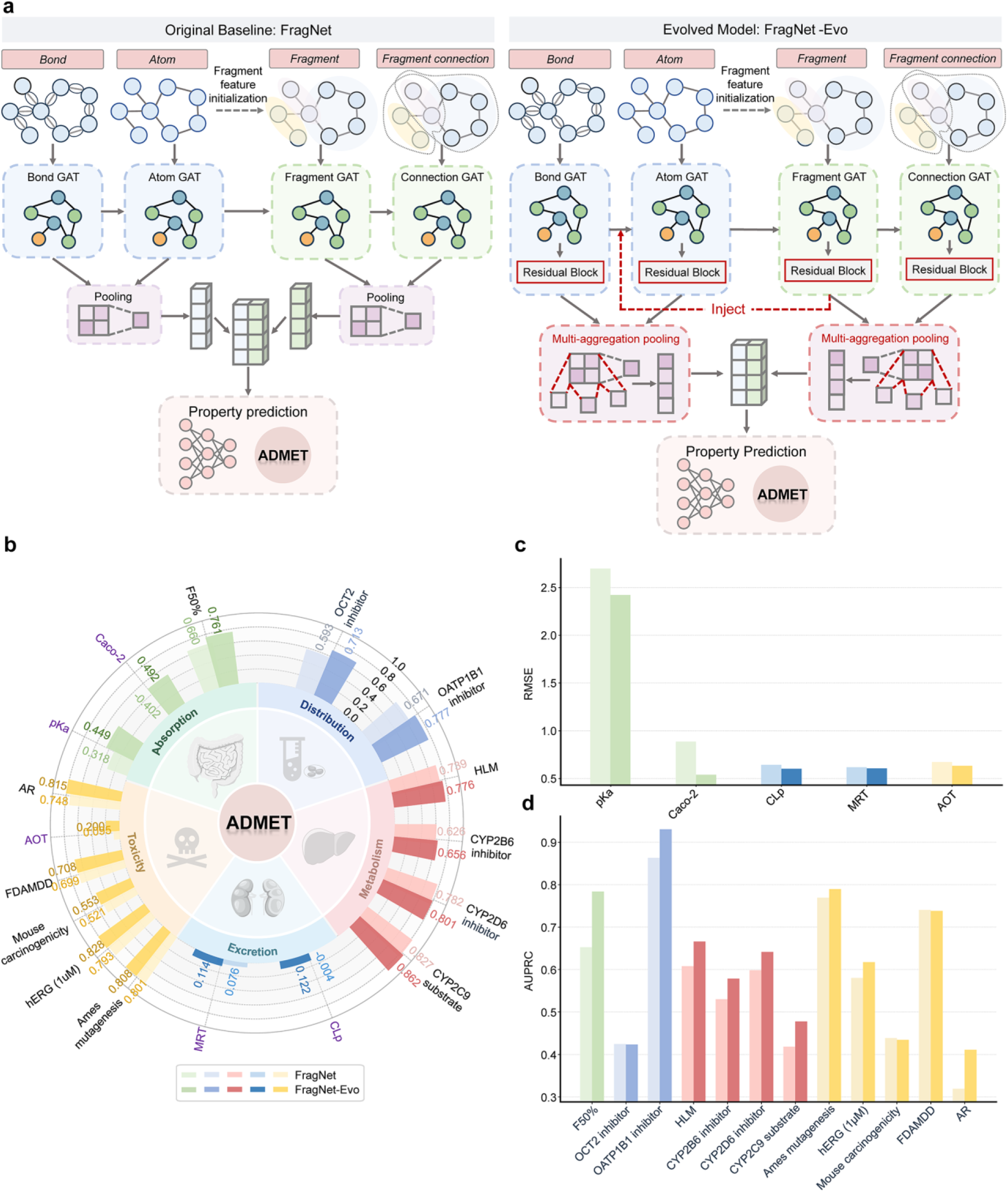
Autonomous evolution of FragNet for ADMET property prediction. **a.** Comparison of the original FragNet and evolved FragNet-Evo architectures. DrugEvolve evolved FragNet by incorporating cross-scale information propagation from fragment-level representations to atom-level features and introducing multi-aggregation pooling to enhance molecular representation learning. **b.** Performance of FragNet and FragNet-Evo across ADMET endpoints spanning absorption, distribution, metabolism, excretion, and toxicity. Each radial bar represents an individual ADMET endpoint grouped according to its corresponding property category. Classification tasks are evaluated by AUROC (black labels), and regression tasks by R^2^ (purple labels). **c.** Comparison of regression performance between FragNet and FragNet-Evo across ADMET endpoints measured by RMSE. **d.** Comparison of classification performance between FragNet and FragNet-Evo across ADMET endpoints measured by AUPRC.

We evaluated FragNet-Evo on 17 datasets curated from the admetSAR 3.0 database^74^, including 12 classification tasks and 5 regression tasks spanning the five major ADMET property categories. Under the stringent molecular similarity-based split, which substantially reduces structural overlap between the training and test sets, FragNet-Evo consistently outperformed FragNet across nearly all classification and regression tasks (**Figure 6b**). Specifically, in classification tasks, FragNet-Evo achieved an average increase of 0.050 in AUROC and 0.046 in AUPRC (**Figure 6b and Figure 6d**). For regression tasks, the average RMSE decreased by 0.143, while the average R^2^ improved by 0.259 (**Figure 6b and Figure 6c**). Notably, on the challenging Caco-2 permeability task, a widely used *in vitro* surrogate for intestinal drug absorption, the model demonstrated a remarkable improvement, with R^2^ jumping from −0.402 to 0.492. This highlights a substantial advancement in the quantitative prediction of oral absorption potential (**Figure 6**). These results highlight improved generalization across structurally dissimilar compounds and confirm that DrugEvolve autonomously evolves molecular graph architectures for reliable ADMET prediction, thereby accelerating the prioritization of candidates with favorable pharmacokinetic and safety profiles.

#### 2.3.3 Preclinical Study

##### DrugEvolve advances animal toxicology assessment

Animal studies are indispensable to preclinical research for evaluating drug efficacy and safety; however, their substantial costs, limited throughput, and ethical concerns have necessitated the development of alternative approaches^76^. By leveraging historical animal study data, AI models can predict animal toxicology for untested compounds, thereby facilitating the reduction of animal experimentation in alignment with the 3Rs (Replacement, Reduction, and Refinement) principles^77^. In this study, building upon data provided in a previous study^78^, we first established a baseline AnimalMLP model to predict 38 rat pathology measures. As illustrated in the **Supplementary Figure S11a**, DrugEvolve evolved the AnimalMLP by redesigning both the molecular representation and the response prediction strategy. Unlike the original AnimalMLP, which relies on a single descriptor-based representation, AnimalMLP-Evo incorporates multiple molecular views, including descriptor features, pretrained chemical language model embeddings, and molecular similarity retrieval. Response patterns retrieved from structurally similar compounds are further integrated as residual corrections to the neural predictions, achieving comprehensive modeling of compound-specific preclinical pathology profiles.

Using the dataset constructed via the workflow from the previous study^78^, we comprehensively evaluated AnimalMLP-Evo and AnimalMLP using compound-based 3-fold cross-validation across four settings: random (Random), low-similarity structure holdout (Structure), ATC therapeutic-class (ATC), and temporal (Time) splits. As shown in **Supplementary Figure S11b** and **Supplementary Figure S11c**, the median condition-level RMSE substantially decreased across all splits, accompanied by a marginal improvement in cosine similarity, indicating a closer recovery of the multivariate animal pathology profiles. Overall, the mean of the four RMSE medians across all splits decreased from 62.727 to 50.123, while the mean cosine similarity increased from 0.985 to 0.990. While the improvement in cosine similarity is marginal, the substantial reduction in RMSE highlights a marked enhancement in quantitative accuracy. This discrepancy arises because cosine similarity solely evaluates directional alignment (i.e., relative patterns), whereas RMSE is highly sensitive to absolute magnitudes, reflecting the model’s improved capability in precise quantitative prediction. The consistent gains across all four partitioning schemes indicate that AnimalMLP-Evo, evolved by DrugEvolve, achieves improved compound-level generalization.

#### 2.3.4 Clinical Trial

##### DrugEvolve enhances qualitative and quantitative predictions of drug-drug interactions

Drug-drug interaction (DDI) prediction plays a critical role in both pharmaceutical research and clinical applications, as it facilitates the early identification of potentially harmful interactions that may precipitate severe adverse drug reactions or necessitate drug withdrawal^79,80^. To facilitate accurate and high-throughput DDI predictions, we adopted the MeTDDI^81^ model as the evolutionary starting point for architecture evolution. While preserving the shared atom-motif backbone and co-attention module of the original MeTDDI, the evolved MeTDDI-Evo augments pairwise interaction modeling by incorporating a substructure-aware representation branch and a complementary feature fusion strategy. This generates more expressive molecular pair representations, which may enhance predictive performance in both DDI classification and quantitative PK fold change prediction (**Supplementary Figure S12a**).

To comprehensively evaluate the generalization ability of the evolved architecture, we followed the original MeTDDI protocol by employing two settings of increasing difficulty involving unseen drugs, where either one or both drugs in an interaction pair were excluded from the training set. As provided in **Supplementary Table S6**, classification performance remained broadly comparable to that of the original MeTDDI across both evaluation protocols, indicating that the evolutionary optimization preserves robust discrimination capability under stringent out-of-distribution conditions. In contrast, substantial improvements were observed in quantitative PK change prediction. Specifically, on the external regression benchmark, the evolved MeTDDI-Evo model reduced the RMSE from 0.991 to 0.859 while increasing the Pearson correlation coefficient from 0.709 to 0.765 (**Supplementary Figure S12b**). These results indicate that the primary benefit of this evolutionary optimization lies in enhancing quantitative prediction accuracy while maintaining classification performance, thereby improving the model’s generalization beyond the training distribution.

##### DrugEvolve advances generalizable drug side-effect prediction

Drug side-effect prediction is essential for drug development, enabling the early identification of adverse reactions to improve drug safety and reduce the risk of unexpected toxicity^82,83^. The SDPred^84^ model was originally developed to overcome the inability of conventional methods to quantitatively estimate side-effect incidence for novel drug candidates without annotated labels. Building upon this foundation, DrugEvolve adopted SDPred as the evolutionary starting point to engineer more robust predictive models. Deviating from SDPred, SDPred-Evo optimizes local feature interaction learning by substituting redundant block-specific encoders with shared residual representations and compact modeling, alongside the addition of task-specific prediction branches for classification and regression (**Figure 7a**).

**Figure 7.**
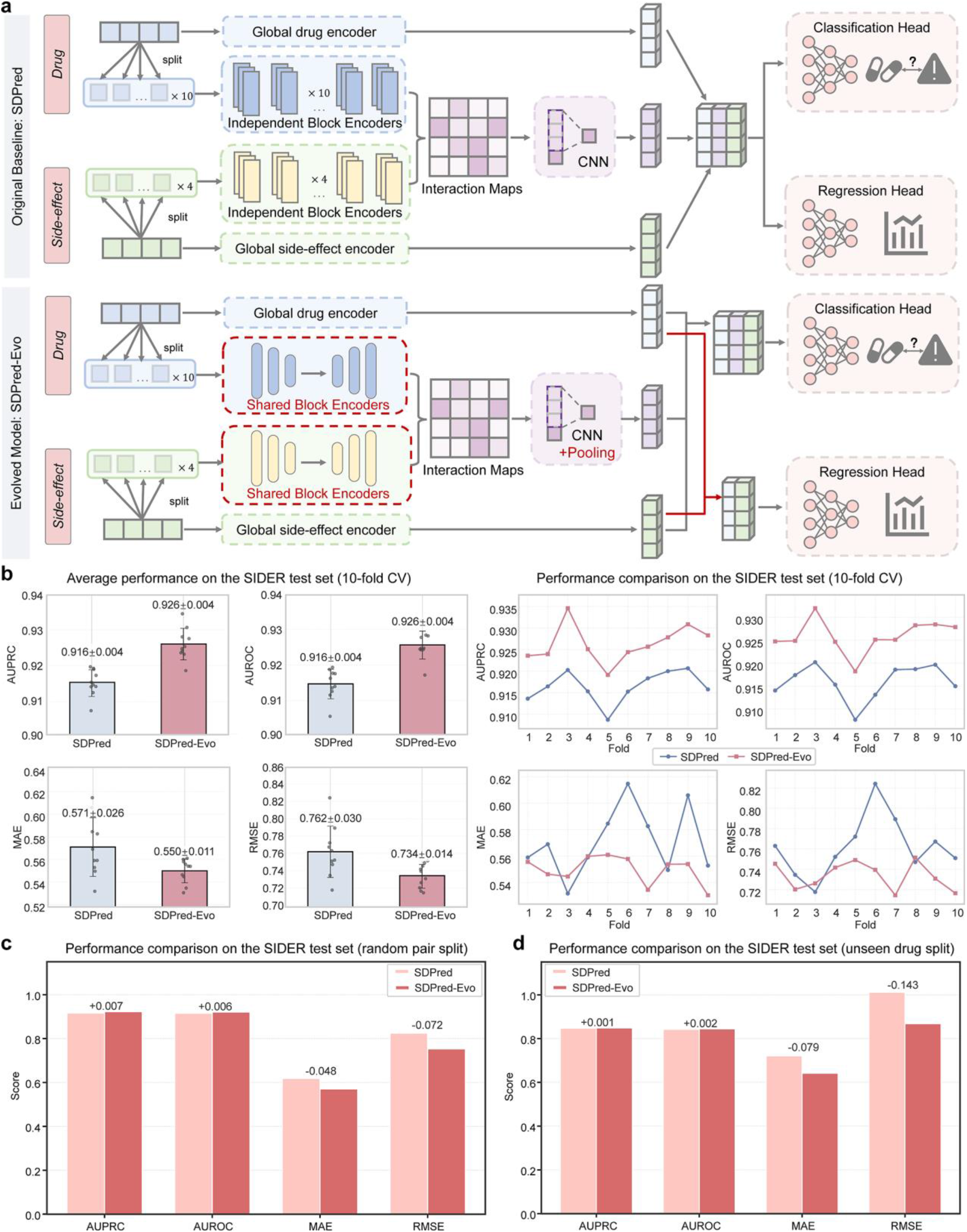
Autonomous evolution of SDPred for drug side-effect prediction. **a.** Comparison of the original SDPred and evolved SDPred-Evo architectures. SDPred-Evo replaces independent block-specific encoders with shared block encoders, introduces CNN-based feature pooling, and incorporates separate classification and regression heads for side-effect prediction. **b.** Ten-fold cross-validation performance comparison between SDPred and SDPred-Evo on the SIDER dataset. Fold-wise AUROC, AUPRC, MAE, and RMSE values are shown, with mean ± s.d. summaries on the left and fold-wise performance trajectories on the right. Individual points represent results from each fold, and error bars indicate the standard deviation across ten folds. **c.** Performance comparison of SDPred and SDPred-Evo on the SIDER test set under the random pair split setting, evaluated using AUROC, AUPRC, MAE, and RMSE. **d.** Performance comparison of SDPred and SDPred-Evo on the SIDER test set under the unseen-drug split setting, evaluated using AUROC, AUPRC, MAE, and RMSE.

Following the original 10-fold cross-validation protocol of SDPred on the SIDER dataset ^84^, SDPred-Evo consistently outperformed SDPred in nearly all test folds (**Figure 7b**). These performance gains were observed in both classification (AUROC and AUPRC) and regression (MAE and RMSE) metrics. Notably, the improvement was pronounced in quantitative side-effect prediction, where the mean RMSE across the ten folds decreased from 0.762 to 0.734 (**Figure 7b**). To further evaluate model generalization, we additionally constructed random splits of drug-side effect pairs. Under this setting, SDPred-Evo achieved classification performance comparable to SDPred, while substantially improving performance on the regression task, reducing the RMSE by 0.072 (**Figure 7c**). Similar trends were observed under the more challenging unseen drug split, where all test drugs were excluded from the training set. In this setting, SDPred-Evo further reduced the RMSE by 0.143 compared to the baseline (**Figure 7d**). The progressively larger gains observed under increasingly stringent evaluation settings suggest that SDPred-Evo captures robust transferable representations of drug-side effect relationships. This capability enables generalization to previously unseen compounds, facilitating more reliable quantitative predictions of side-effect incidence beyond the training distribution.

### 2.4 Evolutionary Dynamics and Mechanisms of DrugEvolve

To have an in-depth understanding of the underlying mechanism of DrugEvolve and guide future optimization, we chose protein druggable sites annotation for a long-term autonomous evolution experiment (exceeding 2,200 GPU hours). As a result, our DrugEvolve system generated a total of 331 architectures surpassing the baseline model ALLSites. Notably, as provided in **Figure 8a**, the cumulative number of superior architectures exhibited a near-linear growth trend, suggesting that the discovery of improved solutions was sustained throughout the evolutionary process rather than being exhausted during the early search stage. This reflects a scaling law for architecture discovery, where increased computational resources translate directly into continuous algorithmic breakthroughs. Such scaling law confirms DrugEvolve’s ability to sustain effective exploration over long-term optimization trajectories, thereby continuously expanding the repertoire of high-performing algorithms for drug development. Therefore, it is reasonable to expect that the marginal performance gains observed in certain tasks under limited computation can be overcome by scaling up computational resources.

**Figure 8.**
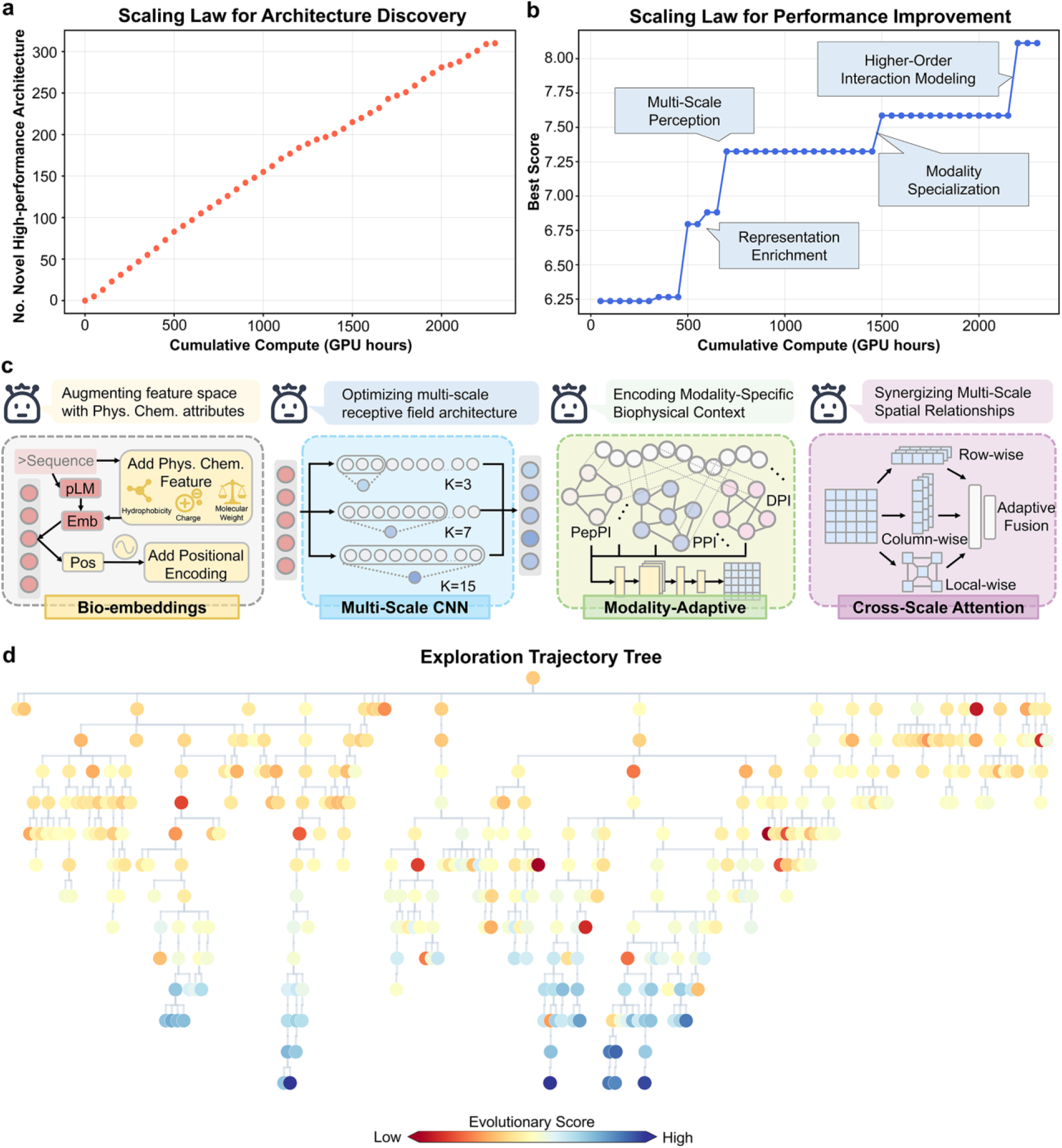
Scaling behavior and evolutionary dynamics of DrugEvolve. **a.** Scaling law between computational resources and architecture discovery. The results show the cumulative number of evolved architectures outperforming the baseline over time (GPU hours). **b.** Scaling law between computational resources and performance improvement. The results show the incumbent best evolutionary score over time (GPU hours). **c.** Representative architectural strategies discovered during evolution. The major design strategies identified during evolution include alternative representation learning, multi-scale feature extraction, task-specific architectural components, and attention-based feature integration. **d.** Evolutionary trajectory of algorithm discovery. Each node represents an evolved algorithm candidate connected through inheritance relationships, and node colors indicate evolutionary scores from lower (red) to higher (blue) values.

A complementary analysis of the incumbent best score further revealed sustained performance improvements throughout the evolutionary search (**Figure 8b**). Instead of prematurely converging to a single optimum, the algorithm improved through multiple successive phases. Notably, each stage of improvement was accompanied by distinct architectural changes, suggesting that DrugEvolve continuously explored alternative algorithmic solutions during the evolutionary process. In-depth examination of the evolved ALLSites-Evo architectures further showed that these design changes encompassed a broad spectrum of algorithmic innovations, including alternative representation learning strategies that potentially capture complementary biochemical characteristics of residues, multi-scale feature extraction mechanisms for integrating local residue patterns with broader sequence contexts, task-specific architectural components that facilitate task-adaptive feature learning, and attention-based modules for modeling long-range contextual dependencies across biological sequences (**Figure 8c**). The repeated emergence of these innovations across independent evolutionary trajectories further supports the capacity of DrugEvolve to discover diverse architectural solutions associated with improved predictive performance. Collectively, the concurrent increase in both the number of high-performing architectures and the incumbent best score indicates that DrugEvolve effectively balances exploration with the iterative refinement of promising solutions, enabling continual algorithmic evolution for drug development.

Furthermore, we leveraged the global evolutionary tree to visualize how DrugEvolve progressively refines algorithmic designs through iterative inheritance and modification (**Figure 8d**). Overall, as evolutionary depth increases, model performance generally improves, consistent with the progressive accumulation of beneficial architectural transformations through iterative inheritance and refinement. Although many evolutionary steps yielded performance improvements, a substantial fraction of descendants performed worse than their parents. Rather than being discarded, these underperforming descendants were retained within the evolutionary memory and analyzed as negative search experiences. This mechanism enables DrugEvolve to harness both successful and unsuccessful evolutionary outcomes to guide subsequent exploration, thereby reducing redundant searches in fruitless regions and preventing premature convergence. Additional mechanistic analyses of evolutionary process are provided in **Supplementary Figure S13**, with detailed discussion available in **Supplementary Discussion S1**.

## 3. Discussion

In this study, we present DrugEvolve, a multi-role LLM-driven framework designed for autonomous algorithm evolution in drug development. It establishes a closed-loop system by combining scientific knowledge, evolutionary memory, algorithm implementation and empirical evaluation. Starting from human-defined tasks and baseline models, DrugEvolve successfully evolved diverse computational models across 11 tasks on 120 benchmark test sets, covering four stages of drug development: target identification, drug discovery, preclinical research and clinical trial. The broad and consistent improvements observed across tasks demonstrate the feasibility and generality of autonomous algorithm evolution, establishing a reference paradigm that substantially expedites the creation of computational tools for the drug development pipeline.

Crucially, the DrugEvolve system is not confined to the specific baselines used in this study but accommodates various established algorithms as evolutionary starting points, thereby enabling complementary optimization trajectories. Beyond the immediate scope of this research, the established paradigm demonstrates broad applicability, allowing for systematic exploration of algorithmic design spaces across the entire drug development pipeline. Furthermore, its modular architecture offers a transferable framework for autonomous evolution in other data-intensive disciplines, extending its utility to fields such as synthetic biology, materials informatics and protein engineering. A critical practical consideration is that the duration of the evolutionary process is inherently tied to the computational overhead of the baseline model, as resource-intensive training and evaluation extend each evolutionary cycle, thereby prolonging the overall optimization trajectory. Consequently, future iterations of DrugEvolve will integrate autonomous engineering capabilities to accelerate model training and inference, substantially bolstering the efficiency of long-horizon evolutionary search. More broadly, this study establishes autonomous algorithm evolution as a scalable and transferable AI software infrastructure, providing a successful proof-of-concept and application paradigm for accelerating computational innovation across drug development and broader scientific discovery.

## 4. Methods

### 4.1 Construction of DrugEvolve

DrugEvolve is built upon four tightly integrated components: a multi-role LLM architecture for autonomous algorithm evolution, a *Cognition Library* for external knowledge augmentation, an *Evolutionary Database* for memory management, and optional sampling strategies for historical knowledge retrieval. Together, these components establish a closed-loop framework that integrates reasoning, external knowledge, evolutionary memory, and adaptive search to support continual algorithm optimization.

#### 4.1.1 Multi-Role Large Language Model Architecture for Algorithm Evolution

The main workflow of DrugEvolve for algorithm evolution is organized around three LLM-based functional domains: the *Researcher*, the *Engineer*, and the *Analyst*. Within each functional domain, multiple agents with distinct roles are powered by diverse LLMs^85^. The specific configurations for these agents are detailed in the **Supplementary Method S1**. At the beginning of the evolutionary loop, human scientists provide a detailed task description, a baseline model that serves as the starting point for evolution, as well as the relevant datasets and predefined evaluation metrics. The DrugEvolve system subsequently initiates the optimization process.

The *Researcher* domain begins with the *Generator*, which first retrieves relevant knowledge from the *Cognition Library* and historical evolutionary records from the *Evolutionary Database*. Knowledge retrieved from the scientific literature is integrated with accumulated experience to inform subsequent algorithm design. Guided by the task objectives specified by human scientists, the *Generator* formulates a rigorous operational plan and establishes the underlying design rationale. The *Inspector* subsequently evaluates whether the proposed algorithm is conceptually redundant with existing solutions. Conceptually redundant proposals are returned to the *Generator* for refinement until a distinct design is obtained. Once a proposal passes inspection, the *Implementer* translates the approved design into high-quality executable code.

The workflow then transitions to the *Engineer* domain, where the generated implementation undergoes execution, debugging, model training, and performance evaluation. The *Trainer* executes the proposed algorithm on the specified datasets using standardized training and validation procedures. If runtime errors or execution failures occur, the *Debugger* attempts to identify and resolve the underlying issues to ensure stable execution. Upon the completion of model training, the *Judger* assesses candidate algorithms primarily using predefined task-specific metrics, with an optional LLM-as-a-judge module providing complementary qualitative insights. Detailed scoring protocols for all tasks examined in this study are provided in the **Supplementary Method S2.**

Following evaluation, the *Analyst* distils the experimental outcomes into structured evolutionary knowledge that informs subsequent iterations. This stage evaluates performance against design objectives and baselines, analyzes the biological or chemical rationale of the evolved design, and identifies both systemic limitations and future optimization pathways. The resulting analytical report serves as a structured knowledge resource for subsequent evolutionary cycles. It also provides a transparent record for human scientists to assess system correctness and rationale. As DrugEvolve is driven by LLMs, differences in prompting schemes may influence agent behaviour and evolutionary outcomes. Analyses of these effects and representative prompt designs and examples are provided in **Supplementary Discussion S1** and **Supplementary Method S3**, respectively.

#### 4.1.2 Cognition Library for External Knowledge-Augmented Algorithm Evolution

To support algorithm evolution across diverse drug development tasks requiring multidisciplinary expertise, we constructed a dedicated *Cognition Library*^18^. Rather than relying exclusively on the internal knowledge of LLMs, DrugEvolve further incorporates evidence from the scientific literature, domain expertise, and prior experimental or computational studies to guide algorithm design^86^. Specifically, curated publications and findings from prior experimental and computational studies were organized into structured entries. These entries summarize key design rationales, methodological strengths and limitations, and relevant biological and pharmaceutical principles. Serving as a complementary source of prior knowledge, the *Cognition Library* expands the scientific context available to the LLM and facilitates more informed, interpretable, and mechanistically grounded algorithm evolution.

#### 4.1.3 Evolutionary Database for Memory Management

To preserve and reuse knowledge accumulated across iterative optimization cycles, DrugEvolve incorporates a dedicated *Evolutionary Database.* It systematically manages algorithm designs, performance evaluations, and analytical reports from each cycle. The limited context window of LLMs makes it difficult to retain complete histories over long evolutionary trajectories, making persistent storage necessary. The database is implemented as a local JSON-based experiment store and supports retrieval by experiment identity, execution order and contextual similarity. Its flexible design also supports different retrieval and sampling strategies. By maintaining an explicit record of algorithm designs, decisions, outcomes, and reports, the database enables evolutionary knowledge to accumulate throughout the optimization process. This persistent memory reduces information loss across iterations, limits redundant exploration of previously evaluated directions and supports more efficient algorithm evolution.

#### 4.1.4 Sampling Strategy for Heterogeneous Evolutionary Search

To support the retrieval of historical evolutionary knowledge across tasks with different search characteristics, DrugEvolve provides two complementary candidate-sampling strategies. The sampling strategy used for each task is predefined before evolution and remains fixed throughout the corresponding evolutionary run. The UCB1 algorithm, a variant of the upper confidence bound (UCB) framework, is used to construct a candidate sampler that balances the retrieval of high-utility candidates with the exploration of less frequently accessed ones^87^. This strategy is especially well-suited for scenarios where candidates can be ranked by scalar utility, enabling the search to refine promising evolutionary lineages while preserving opportunities to explore under-explored alternatives^88^. Another island-based evolutionary sampler is tailored for highly heterogeneous and rugged search landscapes^89^, such as the protein druggable site prediction task characterized by diverse site types in this study. It organizes candidates into island-labeled subpopulations and rotates the preferred island across successive sampling calls. Rather than relying on periodic migration, information is shared across islands through a global archive and by falling back to general candidates when the preferred island is depleted. Additionally, if behavioral feature descriptors are available, the sampler incorporates a shared Multi-dimensional Archive of Phenotypic Elites (MAP-Elites)-style archive^90^. This mechanism retains the highest-utility candidate in each feature space region, assessed via algorithmic complexity, design diversity, and other task-specific characteristics. Overall, the UCB-based strategy emphasizes balancing of exploration and exploitation under uncertainty, whereas the island-based strategy promotes sustained population diversity and broad exploration, thereby reducing the risk of premature convergence in poorly characterized search spaces. Detailed algorithmic descriptions and sampling strategies are provided in the **Supplementary Method S4**.

### 4.2 Application of DrugEvolve to diverse drug development tasks

In this study, DrugEvolve was applied to 11 representative tasks spanning four major stages of drug development. Detailed descriptions of all datasets and data splits used across these tasks are provided in the **Supplementary Method S5**, while the definitions and calculation procedures of the corresponding evaluation metrics for tasks are detailed in **Supplementary Method S6**. During evolutionary optimization, only the training and validation sets were utilized to guide algorithm evolution, while the test sets were strictly reserved for final performance evaluation.

Accordingly, hyperparameters were automatically adjusted throughout the optimization process based on their performance on these two sets. The task-specific optimization objectives, evaluation metrics, and evolutionary goals for all 11 tasks are described below.

#### 4.2.1 Target Identification

##### Cancer gene module detection

This task aims to identify clusters of functionally associated genes underlying cancer pathogenesis. Given a multi-omics integration graph where genes serve as nodes and PPIs define edges, the node features incorporate genomic, epigenomic, and high-throughput chromosome conformation capture (Hi-C)-derived 3D genome architecture. Consequently, the task is formulated as a gene-level classification problem to predict whether each gene is cancer-associated. The CGMega^46^ model addresses this task using a graph Transformer-based architecture. Specifically, stacked TransformerConv layers propagate information across the PPI network via attention-guided neighborhood aggregation, followed by graph pooling and fully connected layers for cancer gene classification. Initiating from the CGMega architecture, DrugEvolve evolved the model structure while preserving the original data preprocessing protocols. During the evolutionary process, performance was assessed on the MCF-7 breast cancer multi-omics dataset, targeting AUROC and AUPRC as the primary optimization objectives, alongside precision and MCC as secondary evaluation criteria. The evolved model, CGMega-Evo, retains the original graph Transformer backbone while introducing a complementary structural representation branch and a residual correction strategy. Within this branch, gene features are first projected into latent representations and encoded with learnable characteristic frequencies to capture frequency-dependent structural patterns. These representations are then utilized to derive neighborhood moments by contrasting observed PPI neighborhood patterns with null expectations. Next, structural fingerprints are generated and interact with the features to produce correction signals. Finally, these signals are fused with the original CGMega predictions, enabling the model to capture higher-order network contexts that extend beyond conventional multi-omics feature aggregation. Collectively, this design preserves the predictive capability of the original framework while enhancing the identification of cancer-associated gene modules through improved gene-network representation.

##### Gene-disease association prediction

This task aims to determine the association between a specific disease and a gene, thereby accelerating the discovery of novel therapeutic targets. Gene function is primarily executed by its encoded proteins, and protein sequence features are highly correlated with disease pathogenesis^91^. Building on this rationale, the FusionGDA^48^ model utilizes protein sequences and disease descriptions as inputs to predict gene-disease associations.

Specifically, a protein encoder (ESM-2^92^) and a text encoder (PubMedBERT^93^) are employed to encode these two entities, and their representations are subsequently integrated by a fusion module for association prediction. The model is first pretrained with a contrastive objective to learn discriminative gene-disease representations and then fine-tuned for GDA prediction. Using the AUROC metric on a randomly split validation set of DisGeNET-EVAL benchmark as the optimization objective, DrugEvolve evolved the FusionGDA architecture by preserving the pretrained encoders while allowing the representation learning and fusion modules to be optimized. Specifically, the evolved FusionGDA-Evo replaces the original linear projection with a pre-normalized two-layer multi-layer perceptron (MLP) projector, enabling entity-specific representations to be transformed prior to fusion. Furthermore, it replaces the attention-based fusion module with a Hamiltonian interaction framework, where protein and disease representations are assigned learnable interaction coordinates and charges to construct energy-based interactions within a shared latent space. These interactions facilitate iterative information propagation between the two entities via Hamiltonian message passing. Subsequently, an energy-aware pooling strategy aggregates both interaction-weighted and global features for downstream prediction. These optimizations enable more structured cross-entity information exchange while preserving essential entity-specific representations.

##### Annotation of protein druggable sites

Prediction of protein druggable sites is formulated as a binary classification task, where each residue in the protein sequence is assigned a binary label indicating whether it serves as a functional or binding site. We adopted the ALLSites^53^ as the baseline model, a Transformer-based framework that uses ESM-2^92^ to extract residue-level representations, followed by a gated convolutional encoder and Transformer decoder for site prediction. Notably, this model is capable of accurately predicting protein binding sites across diverse drug modalities. For evolutionary optimization, we selected a representative benchmark from each of the six non-covalent binding site types (DPI, RPI, PPI, PepPI, SMPI and CarbPI sites), while the remaining datasets were reserved for assessing generalization capabilities. Model evolution is guided by task-specific weighted objective functions. Specifically, distinct subsets of evaluation metrics are aggregated using pre-defined weights to construct a tailored fitness function for each benchmark. To ensure compatibility with the original prediction task, the input and output interfaces are kept unchanged, whereas all architectural components operating on the ESM-2 residue embeddings are open to structural evolution. As a result, DrugEvolve substantially restructures the baseline architecture by replacing the single-scale gated convolutional encoder with a multi-scale feature extractor, and substituting sequential Transformer decoding with parallel attention pooling and feature fusion. Protein and residue representations are independently extracted via attention-based pooling before classification. This architecture enhances multi-scale sequence modeling and representation efficiency, thereby improving generalization across diverse heterogeneous druggable residue prediction tasks.

#### 4.2.2 Drug Discovery

##### Molecular Docking

Molecular docking is a fundamental task in computer-aided drug discovery, which aims to predict the binding position, orientation, and conformation of a ligand given its initial structure and the three-dimensional structure of its target protein. We adopted EquiBind as the baseline, which updates molecular representations and coordinates by joint graph propagation through equivariant layers, followed by attention-based keypoint construction and Kabsch alignment to dock the ligand. While DrugEvolve operates on the raw, uncorrected coordinate outputs of EquiBind (denoted as EquiBind-U), the EquiBind baseline underwent chemical correction via the standard fast conformer-fitting procedure during final evaluation. The PDBbind benchmark was employed for optimization, utilizing ligand RMSD, centroid distance, and Kabsch RMSD to quantify docking quality. Furthermore, all components of the framework were accessible to evolutionary modification. DrugEvolve produced EquiBind-Evo-U and the refined EquiBind-Evo, which retains the equivariant interaction backbone while optimizing two downstream stages. Specifically, a ligand-conditioned pocket anchor focuses keypoint selection on coherent binding regions, and post-Kabsch refinement applies local corrections after global alignment. These algorithm changes improve pocket localization and pose precision while maintaining geometric consistency.

##### Drug-target interaction prediction

This task is formulated as a binary classification task that takes a drug-target pair as input to predict the interaction probability. We employed DrugBAN as our baseline, which leverages a graph convolutional network (GCN) for SMILES-based molecular encoding and a 1D convolutional neural network (1D CNN) for protein sequences. Furthermore, a bilinear attention network (BAN) is integrated for interaction modeling, followed by a fully connected classifier for final prediction^64^. During the architectural evolution process, a random split of the BindingDB dataset was utilized as the development benchmark, with AUROC, AUPRC, and MCC metrics serving as optimization objectives. The BAN module is designated as the primary target for architectural evolution, enabling DrugEvolve to explore alternative formulations of interaction modeling. The evolved architecture replaces the original BAN with a hybrid bilinear interaction module that integrates diagonal and low-rank interaction pathways, thereby enhancing the expressiveness of cross-modal modeling. In addition, a Top-K sparsification mechanism is introduced to refine attention representations, generating sparse interaction maps that highlight informative patterns while suppressing redundant signals^94^. These modifications refine the quality of interaction modeling and are proven to improve predictive performance, demonstrating robust generalization on out-of-distribution datasets.

##### De novo molecular generation

In this task, the model learns the distribution over drug-like molecules from SMILES strings and generates novel, chemically plausible structures with broad structural diversity. We adopted the CharRNN^68^ model as our baseline framework, which maps SMILES tokens to learned embeddings, captures sequential dependencies via a three-layer long short-term memory (LSTM) network, and predicts subsequent tokens based on the output probability distribution. The MOSES^95^ dataset was adopted as the optimization benchmark, utilizing FCD, Scaff, IntDiv2, and Novelty metrics to evaluate distributional fidelity, structural similarity, chemical diversity, and molecular novelty, respectively. Guided by this composite objective, DrugEvolve autonomously optimized the entire CharRNN pipeline to yield an evolved model, termed CharRNN-Evo. This evolved architecture replaces the original vocabulary-sized embedding with a more compact token representation and introduces a residual pathway to preserve embedding information across the recurrent network. For next-token prediction, the LSTM representation is processed by a standard neural output head and a low-rank adapter, which compresses and reconstructs the hidden state before vocabulary projection. The resulting logits are fused with a training-derived bigram prior via a context-dependent gate that adaptively regulates its influence. Together, the new architecture preserves embedding-level information within the recurrent representation and integrates learned sequence features with gated local SMILES statistics, thereby facilitating more realistic and diverse molecular generation.

##### Target-specific peptide binder design

In this task, the model takes a target protein sequence as input and generates potential peptide binders. We employed PepMLM as our baseline, framing peptide design as a target-conditioned masked language modeling task^70^. Specifically, fully masked peptide sequences are appended to the C-terminus of the target sequence and reconstructed using a fine-tuned ESM-2^92^ model. The PepBench dataset was adopted as the optimization benchmark, with the similarity between generated and reference peptides, alongside the internal sequence diversity of generated peptides, defined as the optimization objectives. This dual-objective formulation encourages both target-binding relevance and comprehensive exploration of the peptide sequence space. Given the modular and well-defined design of PepMLM, all components of the framework, including the model architecture, training objective, and generation strategy, are accessible to evolutionary innovation. Starting from PepMLM and guided by the aforementioned optimization objectives, DrugEvolve yields an evolved model termed PepMLM-Evo. This evolved model replaces single-step masked reconstruction with an iterative mask-based refinement framework under a cosine corruption schedule, enabling multi-step denoising of peptide sequences and more expressive modeling of long-range, context-dependent residue interactions. Furthermore, it introduces a contrastive learning algorithm based on shared representations to align interacting proteins and peptides while repelling non-interacting ones. Trained with joint reconstruction and contrastive losses, the model generates peptides via iterative mask refinement and amino-acid-constrained top-k sampling. Collectively, this paradigm shifts couples token-level reconstruction with interaction-aware representation learning, yielding more effective sequence generation under binding constraints and a flexible sampling process across the peptide sequence space.

##### ADMET properties prediction

Taking a compound SMILES string as input, this task aims to accurately predict ADMET properties. The prediction is formulated as either a classification or regression problem, contingent upon the specific endpoint. We utilized FragNet^75^ as the backbone GNN, which hierarchically constructs four complementary representations (bond, atom, fragment, and fragment connection graphs) via bond attribute initialization, atom-to-fragment aggregation, and BRICS-based fragmentation with virtual inter-fragment links. The multi-scale representations are iteratively refined via graph attention message passing, pooled into atom- and fragment-level embeddings, and fused into a unified molecular representation. Ultimately, this representation drives downstream property prediction through self-supervised pretraining and task-specific fine-tuning. Optimizing for classification (AUROC) and regression (RMSE) performance on the validation sets, DrugEvolve evolved the architecture operating on the four pre-constructed molecular graph representations. Specifically, the evolved FragNet-Evo facilitates cross-scale information exchange by adaptively injecting fragment-derived contextual representations into atom-level message passing, enabling hierarchical molecular representations to interact continuously throughout the propagation process. It further stabilizes deep graph representation learning through learnable residual information flow, adaptive feature gating, and normalization. This ensures more effective feature preservation and integration across successive message-passing layers. Moreover, FragNet-Evo extends conventional additive graph pooling by incorporating complementary mean-, max-, and attention-based aggregation pooling mechanisms, yielding more expressive graph-level compound representations while preserving the additive molecular signal. Altogether, these architectural refinements improve cross-scale molecular information integration, stabilize hierarchical representation learning and enhance graph-level representation expressiveness, thereby improving predictive performance across diverse ADMET classification and regression tasks.

#### 4.2.3 Preclinical Study

##### Animal toxicology assessment

This task aims to predict compound-induced animal pathology profiles from historical *in vivo* studies, capturing multi-dimensional physiological responses to facilitate chemical and drug safety assessment. Given molecular descriptors and treatment conditions (e.g., dose and exposure duration), the model predicts a multi-dimensional pathology profile comprising hematological and clinical chemistry measurements. We established AnimalMLP as the evolutionary baseline, utilizing molecular descriptors and exposure conditions as inputs. Specifically, molecular descriptors were standardized and dimensionally reduced via PCA, while a time-dose reference profile was derived from the training compounds. An MLP then predicted compound-specific corrections to the reference profile, which were combined to yield the final pathology profile. Using the benchmark from a previous study^78^, DrugEvolve optimized AnimalMLP based on Cosine similarity and RMSE metrics. The evolved AnimalMLP-Evo advances the original framework primarily by introducing a multi-view molecular representation strategy. This approach integrates two pretrained SMILES-based chemical language models (ChemBERTa-77M-MLM and ChemBERTa-zinc-base-v1^96^) and molecular fingerprints alongside traditional descriptors. Furthermore, a molecular retrieval module was developed to identify analogous training compounds and transfer their residual response profiles via similarity-weighted aggregation. These complementary prediction signals were integrated through a weighted fusion framework, enabling the model to capture both learned molecular-response relationships and global chemical similarity patterns, providing a more comprehensive characterization of compound-specific responses.

#### 4.2.4 Clinical Trial

##### Drug-drug interaction

This task aims to characterize the interaction between two drugs. Using the benchmark dataset provided by MeTDDI^81^, the model takes a pair of drug molecules as input to either predict the specific interaction mechanism among four categories (classification) or estimate the resulting pharmacokinetic (PK) fold change (regression). The baseline model, MeTDDI, takes a pair of drug molecules as input, representing each drug using complementary atom-level and motif-level molecular graphs. Weight-shared Transformer encoders extract multi-scale structural features by integrating atom and motif representations, followed by a co-attention module that models drug interactions. The resulting interaction representations are then utilized for metabolic DDI mechanism classification or PK fold change prediction. DrugEvolve simultaneously utilized classification benchmarks (using AUROC, AUPRC, and ACC metrics) and regression benchmarks (using RMSE and Pearson correlation coefficient) as evolutionary objectives. While preserving the original input-output interface, DrugEvolve was used to evolve the entire model architecture. As a result, the evolved MeTDDI-Evo preserves the shared atom-motif backbone while redesigning the downstream interaction modeling pipeline. Instead of relying solely on co-attention representations, MeTDDI-Evo incorporates a substructure-aware structural representation branch by jointly modeling atom-level features, molecular substructures, and their spatial relationships. These features are then incorporated via an orthogonal residual fusion module, allowing structural information to be integrated into pairwise interaction representations. In summary, the evolved MeTDDI-Evo captures chemically meaningful inter-drug interaction patterns by combining substructure representations with their spatial organization, thereby driving more effective DDI prediction.

##### Drug side-effect prediction

Taking a drug-side effect pair as input, this task is formulated as a multi-task learning problem that jointly performs a primary classification to predict side effect induction and a secondary regression to estimate its occurrence frequency. SDPred^84^ was selected as the baseline architecture. This model encodes both global and block-wise local representations, generates pairwise interaction maps through outer-product operations, extracts interaction features via convolutional neural networks, and fuses the resulting representations to perform joint classification and regression. DrugEvolve autonomously evolved the SDPred architecture by optimizing against a composite objective of classification (AUROC, AUPRC) and regression (MAE, RMSE) metrics across SIDER random-split and unseen-drug split validation sets. Specifically, the evolved SDPred-Evo replaces independent block-specific encoders with shared residual block encoders, introduces low-dimensional interaction projections to construct compact interaction maps before convolutional feature extraction, and redesigns the prediction module into task-specific classification and regression branches. These architectural changes aim to improve parameter efficiency, mitigate redundant high-dimensional interactions, and decouple the optimization of classification and regression, thereby facilitating side-effect prediction.

## Acknowledgements

The National Natural Science Foundation of China (82373790); The Natural Science Foundation of Zhejiang (RG25H300001); Central Guidance on Local Science and Technology Development Fund of the Hebei Provience (226Z2605G); The AI for Science Programs of Shanghai Municipal Commission of Economy and Informatization (2025-GZL-RGZN-BTBX-01013). Supported by Zhejiang University Innovation Institutes for Artificial Intelligence in Medicine, and Information Technology Center of Zhejiang University.

## Author Contributions

F.Z., P.F.L., and M.J.M. conceived and designed the entire study. F.Z., P.F.L., M.J.M., and Z.M.Z. supervised the algorithm development and wrote manuscript. Z.M.Z., Y.N., Y.X.L., W.X.X., and Z.H.X. developed the DrugEvolve system. Z.Y.Z., H.Y., Z.M.Z., Q.X.Y., and Y.H.L. finished the tasks of target identification. Z.M.Z., Y.N., B.L., W.H.J., Y.L.R. and Y.M.W. finished the tasks of drug discovery. Z.M.Z., Y.L., and T.T.M. finished the tasks of the preclinical stage. H.C.S., Y.T.Q., and Y.X.L. finished the tasks of clinical stage. Z.M.Z. and M.J.M. created figures and tables.

## Data Availability

The benchmark datasets used in this study are available on GitHub (https://github.com/GAIR-NLP/DrugEvolve/tree/main/applications/dataset). Source data are provided with this paper.

## Code Availability

The source code for DrugEvolve, together with the baseline and evolved algorithms for all tasks, is available on GitHub (https://github.com/GAIR-NLP/DrugEvolve).

