## Supplementary Information for "An AI System for Autonomous Algorithm Evolution in Drug Development"

##### Supplementary Discussions

**Discussion S1**. Mechanistic insights into evolutionary search in DrugEvolve……………...….46

##### Supplementary Tables

### Supplementary Methods

**Method S1**. Pipeline configurations

**LLM configurations for each functional module.** DrugEvolve supports flexible assignment of foundation models to individual functional modules. The configuration described below represents the recommended default setting used in this study, rather than a mandatory architectural constraint, and may be adapted according to task complexity, model availability, computational budget and domain-specific requirements. To maximize computational efficiency while exploiting the complementary strengths of frontier foundation models, DrugEvolve adopts a heterogeneous model allocation strategy across its distributed multi-role agent architecture. Within the Researcher domain, OpenAI GPT-5.4 serves as the *Generator*, responsible for synthesizing novel algorithmic hypotheses and optimization strategies from historical evolutionary knowledge. To enforce algorithmic novelty with minimal computational overhead, OpenAI GPT-5 mini is deployed as the *Inspector*, performing lightweight, high-throughput redundancy screening to ensure that newly proposed solutions remain conceptually distinct from previously explored designs. Following successful inspection, OpenAI GPT-5.5 acts as the *Implementer*, leveraging its strong software-engineering and code-generation capabilities to translate abstract algorithmic concepts into high-fidelity executable implementations. During the subsequent validation and assessment stages, OpenAI GPT-5.2 is employed as both the *Judger* and the *Analyst*. In these roles, it conducts multi-dimensional performance evaluation, performs retrospective performance-attribution analysis to identify the architectural factors underlying success or failure, and generates comprehensive interpretability reports for subsequent verification by human scientists.

**General principles for scoring function design**. We designed an evolutionary scoring framework to guide the sampling process by prioritizing candidate nodes with the greatest potential for further evolution and propagation. The framework primarily relies on rule-based performance scores, in which task-specific scoring functions are defined according to the evaluation metrics most relevant to each task. This component directly measures the performance change associated with an algorithm before and after evolution, thereby providing an explicit and quantitative basis for comparison. For tasks evaluated in this study, the evolutionary score was determined exclusively by the rule-based performance score. An evolutionary search process driven exclusively by quantitative performance metrics can be inherently myopic and may encourage reward hacking, a limitation that has been extensively discussed in the AI safety literature[1]. DrugEvolve additionally provides an optional *LLM-as-judge* module for complementary qualitative assessment. This module evaluates the rationality of the algorithmic design, model complexity and consistency between the proposed motivation, implementation and observed performance changes. The resulting *LLM-as-judge* score was retained as a complementary reference for interpreting evolutionary outcomes and supporting subsequent analysis by the *Analyst* and human researchers.

**Computational resources.** All evolutionary experiments reported in this study were conducted using NVIDIA A100 GPUs with 40 GB of memory. DrugEvolve is hardware-agnostic and does not impose specific requirements on GPU type. In practice, the computational resources required for each evolutionary task are determined primarily by the baseline model, dataset scale, and training and evaluation settings. Therefore, the framework can be deployed on alternative GPU configurations.

**Method S2**. Task-specific scoring framework for evolutionary optimization

#### S2.1 Target Identification

##### *S2.1.1 Cancer gene module detection*

To evaluate candidate models for cancer gene module detection, we constructed a composite scoring function that prioritizes ranking performance while incorporating complementary threshold-dependent measures of classification reliability. The scoring framework uses AUPRC and AUROC as the principal metrics, with precision and MCC providing auxiliary contributions only when ranking performance is maintained.

Let $x_{AUPRC}$, $x_{AUROC}$, $x_{precision}$ and $x_{MCC}$ denote the candidate AUPRC, AUROC, Precision and MCC, respectively, and let $b_{AUPRC}$, $b_{AUROC}$, $b_{precision}$ and $b_{MCC}$ denote the corresponding values obtained by the original CGMega baseline. The candidate and baseline ranking statistics were defined as

$$R_c = 0.70x_{AUPRC} + 0.30x_{AUROC} \quad (1)$$

$$R_b = 0.70b_{AUPRC} + 0.30b_{AUROC} \quad (2)$$

AUPRC was assigned the larger weight because it more directly reflects ranking quality under class imbalance. The ranking statistic was transformed into a bounded core score using a sigmoid mapping centered at the baseline:

$$s_{\text{rank}} = 5 + 0.9 \times \left(\frac{10}{1 + \exp[-12(R_c - R_b)]} - 5\right) \quad (3)$$

This transformation assigns a score of 5 to baseline-equivalent ranking performance and maps improvements or degradations smoothly above or below this reference point. The factor of 0.9 slightly contracts the effective score range, reducing the influence of small ranking fluctuations on evolutionary selection.

To distinguish models with similar ranking performance, auxiliary changes in Precision and MCC were calculated as

$$\Delta_{precision} = x_{precision} - b_{precision} \quad (4)$$

$$\Delta_{MCC} = x_{MCC} - b_{MCC} \quad (5)$$

The precision component was defined as

$$s_{precision} = \begin{cases} 1.5\Delta_{precision}, & \Delta_{precision} < 0, \\ 2.0\Delta_{precision}, & 0 \le \Delta_{precision} < 0.03, \\ 0.06 + 4.0(\Delta_{precision} - 0.03), & 0.03 \le \Delta_{precision} < 0.10, \\ 0.34 + 7.0(\Delta_{precision} - 0.10), & \Delta_{precision} \ge 0.10. \end{cases} \quad (6)$$

The MCC component used the same improvement schedule, with an additional penalty for negative MCC values:

$$s_{MCC} = \begin{cases} -1 + 2x_{MCC}, & x_{MCC} < 0, \\ 1.5\Delta_{MCC}, & x_{MCC} \ge 0 \text{ and } \Delta_{MCC} < 0, \\ 2.0\Delta_{MCC}, & 0 \le \Delta_{MCC} < 0.03, \\ 0.06 + 4.0(\Delta_{MCC} - 0.03), & 0.03 \le \Delta_{MCC} < 0.10, \\ 0.34 + 7.0(\Delta_{MCC} - 0.10), & \Delta_{MCC} \ge 0.10. \end{cases} \quad (7)$$

These piecewise transformations provide modest rewards for marginal improvements and stronger rewards for substantial gains, while penalizing deteriorated precision or MCC. Negative MCC is penalized more strictly because it indicates predictions worse than random association. The auxiliary decision score was defined as

$$s_{\text{decision}} = 0.60s_{precision} + 0.40s_{MCC} \quad (8)$$

Because threshold-dependent metrics should not compensate for degraded ranking quality, the decision score was modulated by a ranking-preserving gate. Using the sigmoid function

$$\sigma(z) = \frac{1}{1 + \exp(-z)} \quad (9)$$

the gate was defined as

$$g = \sigma\big(10(R_c - R_b)\big)\sigma(30[x_{AUPRC} - (b_{AUPRC} - 0.01)])\sigma(30[x_{AUROC} - (b_{AUROC} - 0.005)]) \quad (10)$$

This gate suppresses precision- and MCC-based rewards when the weighted ranking statistic deteriorates, or when AUPRC and AUROC fall materially below their baseline values. An explicit ranking penalty was additionally introduced:

$$\mathcal{P}_{\text{rank}} = 8 \times \max(b_{\text{AUPRC}} - x_{\text{AUPRC}}, 0) + 4 \times \max(b_{AUROC} - x_{AUROC}, 0) \quad (11)$$

The final evolutionary score was calculated as

$$S = \text{clip}_{[0,10]}(S_{\text{rank}} + gS_{\text{decision}} - \mathcal{P}_{\text{rank}}) \quad (12)$$

The objective rewards simultaneous improvements in ranking ability and threshold-dependent classification reliability, while preventing models from obtaining a high fitness score by sacrificing AUPRC or AUROC in exchange for isolated gains in Precision or MCC.

##### *S2.1.2 Gene-disease association prediction*

For the gene-disease association prediction task, we used a baseline-referenced score based on AUROC obtained from a randomly selected validation split of the DisGeNET-EVAL[2] dataset. Since AUROC serves as the primary evaluation metric for binary association prediction, the scoring function is formulated directly upon the AUROC without introducing additional metric aggregation. Specifically, the original FusionGDA[2] model is assigned a reference score of 5.0, while the theoretical optimum (AUROC = 1.0) corresponds to the maximum score of 10.0.

Let $x_{AUROC}$ denote the validation AUROC achieved by the candidate architecture, and let $b_{AUROC}$ denote the validation AUROC of the original FusionGDA baseline. The unbounded baseline-referenced score was defined by linear interpolation as

$$S^* = 5 + 5 \times \frac{x_{AUROC} - b_{AUROC}}{1 - b_{AUROC}} \quad (13)$$

To maintain numerical stability and bounded interpretability, the final score was clipped to the interval [0,10]:

$$S = \mathrm{clip}_{[0,10]}(S^*) \quad (14)$$

Under this formulation, models performing below the baseline receive scores lower than 5.0, whereas improved architectures approach the upper bound as AUROC converges toward perfect discrimination. The score therefore provides a continuous optimization objective that preserves the magnitude of incremental AUROC improvements relative to a fixed reference model.

##### *S2.1.3 Protein druggable sites annotation*

To enable consistent comparisons across heterogeneous biomolecular interaction prediction tasks, we used a baseline-relative scoring framework that quantifies model utility according to improvements over a fixed reference method. Six representative benchmark tasks, CarbPI, DPI, PepPI, PPI, RPI and SMPI sites, were included. For each task, the corresponding baseline model was assigned a reference score of 5.0, establishing a common calibration point between performance improvement and degradation.

Because evaluation metrics differ in numerical range, practical sensitivity and biological relevance, raw performance differences were not aggregated directly. For task $t$ and metric $m$, let $x_{t,m}$ and $b_{t,m}$ denote the candidate and baseline values, respectively. The raw improvement was calculated as

$$\Delta_{t,m} = x_{t,m} - b_{t,m} \quad (15)$$

Each raw improvement was then normalized by a predefined metric-specific scaling factor $\lambda_m$:

$$g_{t,m} = \frac{\Delta_{t,m}}{\lambda_m} \tag{16}$$

The normalization constants were fixed before evolutionary optimization according to the expected numerical variation and practical difficulty of improving each metric. Metrics such as AUROC, MCC, AUPRC and F1-score, which typically show smaller numerical fluctuations, were assigned smaller normalization constants, whereas Recall and Precision were assigned larger constants to prevent minor variations from producing disproportionate score changes.

For each benchmark task, only the predefined subset of biologically relevant metrics contributed to the final score. The weighted task-level improvement was computed as

$$G_t = \sum_{m \in \mathcal{M}_t} w_{t,m}\, g_{t,m} \tag{17}$$

$$\sum_{m \in \mathcal{M}_t} w_{t,m} = 1 \tag{18}$$

The task-specific score was then defined as

$$S_t = \mathrm{clip}_{[0,10]}(5 + 5G_t) \tag{19}$$

The overall score was computed as the arithmetic mean across the six benchmark tasks:

$$S = \frac{1}{|\mathcal{T}|} \sum_{t \in \mathcal{T}} S_t \tag{20}$$

All metric selections, normalization constants and task-specific weights were specified before model evaluation and remained fixed throughout all experiments. This design prevents post hoc adjustment and ensures that improvements in informative and difficult metrics, such as MCC and F1-score, exert greater influence than equivalent numerical changes in secondary indicators. The complete task-specific metric selections, weighting schemes and normalization constants are summarized in **Supplementary Table S7**.

#### S2.2 Drug Discovery

##### *S2.2.1 Molecular docking*

To guide the evolution of EquiBind[3] architectures toward improved molecular docking performance, each candidate architecture was trained from scratch and evaluated under a standardized validation protocol. Instead of selecting the best single validation epoch, which can introduce substantial variance when metrics fluctuate during training, the evaluation procedure used late-window averaging over the final $K$ epochs.

For metric $m$, the late-window candidate estimate was defined as

$$\bar{x}_m = \frac{1}{K} \sum_{t=T-K+1}^{T} x_{m,t}\,, \qquad K = 10 \tag{21}$$

where $T$ denotes the total number of completed training epochs and $K$ denotes the averaging window. The corresponding baseline estimate is denoted by $\bar{b}_m$. This procedure reduces sensitivity to transient fluctuations and provides a more stable fitness signal for evolutionary selection.

Candidate architectures were evaluated across three equally weighted metric families: ligand pose accuracy, binding-site localization and ligand conformational fidelity. Direction-adjusted relative changes were defined as

$$\Delta_m = \begin{cases} \dfrac{\bar{x}_m - \bar{b}_m}{|\bar{b}_m|}, & \text{if higher values indicate better performance} \\ \dfrac{\bar{b}_m - \bar{x}_m}{|\bar{b}_m|}, & \text{if lower values indicate better performance} \end{cases} \tag{22}$$

The ligand pose accuracy term combined the fraction of validation complexes with ligand RMSD below 5 Å and the median ligand RMSD across the validation set:

$$\Delta_{\text{pose}} = 0.5\frac{\bar{x}_{\text{RMSD<5}} - \bar{b}_{\text{RMSD<5}}}{|\bar{b}_{\text{RMSD<5}}|} + 0.5\frac{\bar{b}_{\text{medRMSD}} - \bar{x}_{\text{medRMSD}}}{|\bar{b}_{\text{medRMSD}}|} \tag{23}$$

The binding-site localization term used the fraction of validation complexes with centroid distance below 2 Å:

$$\Delta_{\text{site}} = \frac{\bar{x}_{\text{centroid<2}} - \bar{b}_{\text{centroid<2}}}{|\bar{b}_{\text{centroid<2}}|} \tag{24}$$

The conformational fidelity term used Kabsch RMSD after optimal rigid-body alignment, where lower values indicate better agreement between predicted and native ligand conformations:

$$\Delta_{\text{conf}} = \frac{\bar{b}_{\text{Kabsch}} - \bar{x}_{\text{Kabsch}}}{|\bar{b}_{\text{Kabsch}}|} \tag{25}$$

The overall docking improvement was calculated by equally weighting the three metric families:

$$G = \frac{\Delta_{\text{pose}} + \Delta_{\text{site}} + \Delta_{\text{conf}}}{3} \quad (26)$$

The final evolutionary score was obtained by linearly scaling the overall improvement around a baseline-equivalent score of 5:

$$S = \text{clip}_{[0,10]}(5 + K \times G), \qquad K = 25 \quad (27)$$

An architecture matching the baseline across the aggregated metrics therefore receives a score of 5. Improvements relative to the baseline increase the score above 5, whereas overall degradation reduces it below 5. Candidates that failed during model construction, training, result generation or validation metric extraction were assigned a score of 0, including runs with execution timeouts, missing result files, unavailable metrics or invalid values such as $NaN$.

##### *S2.2.2 Drug-target interactions*

To construct a unified evaluation framework for drug-target interaction (DTI) prediction, heterogeneous metrics were first mapped into a common normalized utility space. Let $x_m$ denote the candidate value of metric $m$ and let $\tilde{x}_m$ denote its normalized utility. For threshold-independent discrimination, AUROC was normalized by anchoring random performance at zero and perfect discrimination at one:

$$\tilde{x}_{\text{AUR}OC} = \text{clip}_{[0,1]}\left(\frac{x_{\text{AUROC}} - 0.5}{0.5}\right) \quad (28)$$

For precision-recall evaluation, AUPRC was normalized relative to the empirical positive rate $p$, thereby accounting for the pronounced class imbalance in DTI benchmarks:

$$\tilde{x}_{\text{AUPRC}} = \text{clip}_{[0,1]}\left(\frac{x_{\text{AUPRC}} - p}{1 - p}\right) \quad (29)$$

For MCC, which naturally spans [-1,1], a linear rescaling was used:

$$\tilde{x}_{\text{MCC}} = \text{clip}_{[0,1]}\left(\frac{x_{\text{MCC}} + 1}{2}\right) \quad (30)$$

All remaining metrics, including accuracy, recall, precision, specificity, and F1, were already bounded under their standard probabilistic interpretation and were therefore clipped directly.

The composite DTI score was then calculated as a weighted aggregation of normalized metrics. Metrics were grouped into robustness-oriented indicators (MCC and F1), ranking-quality indicators (AUROC and AUPRC), and threshold-dependent classification indicators (Accuracy, Recall, Precision and Specificity), with group-level weights of 40%, 30% and 30%, respectively.

$$S = 10 \times \mathrm{clip}_{[0,1]}\left(\sum_{m\in\mathcal{M}} w_m\,\tilde{x}_m\right), \qquad \sum_{m\in\mathcal{M}} w_m = 1 \tag{31}$$

This formulation transforms heterogeneous DTI metrics into a bounded 0-10 utility scale while preserving the relative importance of robustness, ranking quality and decision-level classification reliability.

##### *S2.2.3 De novo molecular generation*

To evaluate candidate models for molecular generation, we constructed a baseline-relative scoring function using four complementary metrics: FCD, Scaff, IntDiv2 and Novelty. These metrics jointly assess distributional similarity, scaffold coverage, internal diversity and novelty of generated molecules. For metric $m$, $x_m$ denotes the candidate value and $b_m$ denotes the CharRNN[4] baseline value.

The direction-adjusted relative change was defined as

$$\Delta_m = \begin{cases} \dfrac{x_m - b_m}{|b_m|}, & \text{if higher values indicate better performance,} \\ \dfrac{b_m - x_m}{|b_m|}, & \text{if lower values indicate better performance.} \end{cases} \tag{32}$$

Each relative change was transformed into a bounded metric-level score:

$$s_m = \mathrm{clip}_{[0,10]}\big(5 + 5\tanh(3\Delta_m)\big) \tag{33}$$

Under this transformation, performance identical to the baseline corresponds to a score of 5, whereas improvements and degradations are mapped smoothly above and below this reference point. The hyperbolic tangent function limits the influence of extreme relative changes while preserving sensitivity near the baseline.

The four metric-level scores were then combined using a weighted mean, with weights of 0.30, 0.25, 0.20 and 0.25 assigned to FCD, Scaff, IntDiv2 and Novelty, respectively:

$$S_{\mathrm{raw}} = \frac{\sum_{m\in\mathcal{M}} w_m\, s_m}{\sum_{m\in\mathcal{M}} w_m} \tag{34}$$

Because diversity- and novelty-related statistics can become unreliable when a generator produces a substantial proportion of invalid SMILES, a validity-dependent safeguard was introduced:

$$S_{\mathrm{valid}} = \begin{cases} S_{\mathrm{raw}}\dfrac{v}{v_{\mathrm{min}}}, & v < v_{\mathrm{min}}, \\ S_{\mathrm{raw}}, & v \geq v_{\mathrm{min}}, \end{cases} \qquad v_{\mathrm{min}} = 0.8 \tag{35}$$

Here, $v$ denotes the proportion of valid generated SMILES. The final molecular generation score was clipped to $[0,10]$:

$$S = \mathrm{clip}_{[0,10]}(S_{\mathrm{valid}}) \tag{36}$$

This objective rewards simultaneous improvements in molecular distribution matching, scaffold coverage, diversity and novelty, while penalizing degenerate solutions that achieve favorable secondary metrics at the expense of chemical validity.

##### *S2.2.4 Target-specific peptide binder design*

For target-specific peptide sequence design, the score was formulated as a bounded, baseline-relative utility that prioritizes sequence-level similarity while conditionally rewarding sequence diversity. PepMLM was used as the reference model[5] , with performance on the PepBench dataset serving as the optimization objective. Let $x_{\mathrm{sim}}$ and $x_{\mathrm{div}}$ denote the candidate mean similarity and sequence diversity, respectively, and let $b_{\mathrm{sim}}$ and $b_{\mathrm{div}}$ denote the corresponding PepMLM baseline values.

Similarity was treated as the dominant component because preservation of target-relevant sequence characteristics is the primary requirement for peptide generation. The similarity term was transformed using a steep sigmoid function centered at the baseline value:

$$s_{\mathrm{sim}} = 5 + 0.9\left(\frac{10}{1 + \exp[-10(x_{\mathrm{sim}} - b_{\mathrm{sim}})]} - 5\right) \tag{37}$$

This mapping assigns approximately 5 to baseline-level similarity and maps improvements or degradations smoothly within a compressed 0-10 range. Sequence diversity was evaluated using a piecewise reward term:

$$s_{\mathrm{div}} = \begin{cases} 0.3(x_{\mathrm{div}} - b_{\mathrm{div}}), & x_{\mathrm{div}} < 0.80 \\ 3.0(x_{\mathrm{div}} - 0.80) + 0.06, & 0.80 \le x_{\mathrm{div}} < 0.95 \\ 3.0 \times 0.15 + 0.06 + 8.0(x_{\mathrm{div}} - 0.95), & x_{\mathrm{div}} \ge 0.95 \end{cases} \tag{38}$$

This design attenuates diversity rewards in low-diversity regions and amplifies them only after diversity reaches a higher operating range, where additional increases more plausibly reflect meaningful exploration of the generated sequence space.

To prevent diversity from compensating for insufficient target consistency, the diversity contribution was regulated by a similarity-dependent gate:

$$g = \frac{1}{1 + \exp[-8(x_{\mathrm{sim}} - b_{\mathrm{sim}})]} \tag{39}$$

Because the gate increases primarily when candidate similarity exceeds the baseline, diversity gains are rewarded mainly when target-consistent sequence resemblance is maintained. The dataset-level score was therefore defined as

$$S = \mathrm{clip}_{[0,10]}(s_{\mathrm{sim}} + g\, s_{\mathrm{div}}) \tag{40}$$

The resulting score should be interpreted as an optimization utility rather than a universal benchmark metric. When multiple datasets are evaluated, the final objective is obtained by macro-averaging dataset-level scores with equal weighting, thereby preventing any individual benchmark subset from dominating the overall evaluation.

##### *S2.2.5 ADMET properties prediction*

To ensure fair comparison among candidate architectures, each evolved model was evaluated on all 17 ADMET prediction tasks using an identical two-stage training protocol. In the first stage, the evolved molecular encoder was pretrained on the training set through a fixed three-dimensional geometric prediction objective, which remained unchanged throughout evolution. In the second stage, the pretrained encoder was independently fine-tuned on each downstream ADMET task. AUROC was used for the 12 classification tasks and RMSE for the 5 regression tasks.

Because AUROC is maximized whereas RMSE is minimized, the evolutionary objective was formulated in terms of direction-adjusted relative improvement over the baseline model. For classification task $i$, the relative improvement was defined as

$$\Delta_i^{\mathrm{cls}} = \frac{x_i^{\mathrm{AUROC}} - b_i^{\mathrm{AUROC}}}{\left|b_i^{\mathrm{AUROC}}\right|} \tag{41}$$

For regression task $j$, the relative improvement was defined as

$$\Delta_j^{\mathrm{reg}} = \frac{b_j^{\mathrm{RMSE}} - x_j^{\mathrm{RMSE}}}{\left|b_j^{\mathrm{RMSE}}\right|} \tag{42}$$

If a candidate architecture failed to complete training or produced invalid predictions for a task, the corresponding relative improvement was assigned the maximum penalty $\Delta = -1$.

Relative improvements were averaged separately over classification and regression tasks:

$$\bar{\Delta}_{\mathrm{cls}} = \frac{1}{N_{\mathrm{cls}}} \sum_{i=1}^{N_{\mathrm{cls}}} \Delta_i^{\mathrm{cls}} \tag{43}$$

$$\bar{\Delta}_{\text{reg}} = \frac{1}{N_{\text{reg}}} \sum_{j=1}^{N_{\text{reg}}} \Delta_j^{\text{reg}} \tag{44}$$

The two task groups were then combined with equal importance:

$$\bar{\Delta} = \frac{\bar{\Delta}_{\text{cls}} + \bar{\Delta}_{\text{reg}}}{2} \tag{45}$$

The final evolutionary score was computed as

$$S = \text{clip}_{[0,10]}(5 + 100\bar{\Delta}) \tag{46}$$

This scoring framework assigns a baseline score of 5.0, such that architectures outperforming the baseline receive scores greater than 5.0, whereas inferior architectures receive lower scores. Averaging classification and regression performance independently before combining them prevents the larger number of classification tasks from dominating the objective. Expressing performance as relative improvement also enables AUROC and RMSE to contribute comparably despite their different numerical ranges.

##### S2.3 Preclinical Study

###### *S2.3.1 Animal toxicology assessment*

To evaluate candidate models for animal clinical pathology prediction, we designed a scoring function for four compound-level data splits: Random, Structure, ATC and Time. Each split was evaluated independently using three complementary metrics: median RMSE, mean RMSE and median cosine similarity. Median RMSE was treated as the primary optimization target because it reflects the typical prediction error across treatment conditions and is less sensitive to outlier compounds. Mean RMSE was included as a secondary error-based criterion to penalize models with large errors in difficult conditions, whereas median cosine similarity was used to capture the preservation of the overall clinical pathology response pattern.

For benchmark partition $j$, let $x_{j,\text{RMSEmed}}$, $x_{j,\text{RMSEmean}}$ and $x_{j,\text{CosMed}}$ denote the candidate model values for median RMSE, mean RMSE and median cosine similarity, respectively. The corresponding baseline values obtained from benchmark are denoted by $b_{j,\text{RMSEmed}}$, $b_{j,\text{RMSEmean}}$ and $b_{j,\text{CosMed}}$. Because RMSE is minimized whereas cosine similarity is maximized, the direction-adjusted relative changes were defined as

$$\Delta_{j,\text{RMSEmed}} = \frac{b_{j,\text{RMSEmed}} - x_{j,\text{RMSEmed}}}{\max(|b_{j,\text{RMSEmed}}|, \epsilon)} \tag{47}$$

$$\Delta_{j,\text{RMSEmean}} = \frac{b_{j,\text{RMSEmean}} - x_{j,\text{RMSEmean}}}{\max(|b_{j,\text{RMSEmean}}|, \epsilon)} \tag{48}$$

and

$$\Delta_{j,\text{CosMed}} = \frac{x_{j,\text{CosMed}} - b_{j,\text{CosMed}}}{\max(|b_{j,\text{CosMed}}|, \epsilon)} \tag{49}$$

where $\epsilon = 10^{-8}$ was used only to avoid numerical instability. Each relative change was then transformed into a bounded metric-level score using

$$s(\Delta_{j,m}; \tau_m) = \text{clip}_{[0,10]}\left[5 + 5\tanh\left(\frac{\Delta_{j,m}}{\tau_m}\right)\right] \tag{50}$$

Under this transformation, performance identical to the AnimalMLP baseline receives a score of 5, improvements over the baseline receive scores above 5, and degradations receive scores below 5. The scale parameter $\tau_m$ controls the sensitivity of the score to relative changes. For the two RMSE-based metrics, $\tau_{\text{RMSEmed}} = \tau_{\text{RMSEmean}} = 0.04$, whereas for cosine similarity, $\tau_{\text{CosMed}} = 0.002$, reflecting the much narrower numerical range of cosine similarity values near one. The raw benchmark score was calculated as a weighted aggregation of the three metric-level scores:

$$S_{j,\text{raw}} = 0.60\, s(\Delta_{j,\text{RMSEmed}}; 0.04) + 0.25\, s(\Delta_{j,\text{RMSEmean}}; 0.04) + 0.15\, s(\Delta_{j,\text{CosMed}}; 0.002) \tag{51}$$

The larger weight assigned to median RMSE emphasizes robust condition-level error reduction, while the mean RMSE and cosine similarity terms provide additional pressure to reduce large errors and preserve the global clinical pathology response pattern. To prevent incomplete prediction outputs from receiving inflated scores, a coverage-dependent safeguard was applied. Let $q_j$ denote the fraction of evaluated treatment conditions with valid predictions in benchmark partition $j$. The final partition-level evolutionary score was defined as

$$S_j = clip_{[0,10]}(\gamma_j S_{j,\text{raw}}) \tag{52}$$

where

$$\gamma_j = 1 - (1 - q_j)1\{q_j < 0.999\} \tag{53}$$

denotes the coverage adjustment factor. The coverage adjustment further ensures that candidate models are favored only when they produce valid predictions across essentially all required compound-dose-time treatment conditions. The overall evolutionary score was calculated as the arithmetic mean of the partition-level scores across the Random, Structure, ATC and Time splits.

#### S2.4 Clinical Trial

##### *S2.4.1 Drug-drug interactions*

To provide a unified optimization objective for drug-drug interaction (DDI) prediction, we define a hierarchical performance score that jointly evaluates both classification and regression tasks. During evolutionary optimization, model performance is assessed exclusively on validation data using eight evaluation metrics spanning three validation settings, including classification scenarios (unseen one drug and unseen two drug settings) and one regression validation scenario. Since these metrics differ in numerical scale, optimization difficulty, and directionality, each metric is first transformed into a bounded utility score relative to the performance of the original MeTDDI[6] architecture.

For each evaluation metric $m$, the baseline model is assigned a reference score of 5.0. Performance improvements above the baseline receive scores greater than 5.0, whereas performance degradation results in scores below 5.0. Rather than adopting a linear transformation, a hyperbolic tangent mapping is employed to provide smooth saturation for large performance gains while preserving sensitivity around the baseline,

$$s_m = 5 + 5 \times \tanh(k\Delta_m) \tag{54}$$

where $k$ is a scaling coefficient controlling the sensitivity of the score function, and

$$\Delta_m = \begin{cases} \dfrac{x_m - b_m}{|b_m|}, & \text{if higher values indicate better performance} \\ \dfrac{b_m - x_m}{|b_m|}, & \text{if lower values indicate better performance} \end{cases} \tag{55}$$

Here, $b_m$ denotes the corresponding validation metric achieved by the original MeTDDI architecture. In this study, the scaling coefficient was fixed at $k = 3.0$, providing a balance between sensitivity to modest improvements and saturation for substantially better-performing architectures. The resulting score is further constrained to the interval $[0,10]$ to ensure bounded interpretability throughout the evolutionary optimization.

Building upon the metric-level scores, the three metrics within each classification scenario are averaged to obtain the corresponding scenario score, while the RMSE and Pearson correlation coefficient are averaged to obtain the regression score on the regression dataset. The two classification scenario scores are subsequently averaged to produce the overall classification score. Finally, equal importance is assigned to classification and regression performance, yielding the overall optimization objective:

$$S = \frac{S_{\text{cls}} + S_{\text{reg}}}{2} \tag{56}$$

##### *S2.4.2 Side-effect prediction*

To encourage architectures that generalize beyond individual data partitioning strategies, the side-effect prediction objective jointly considered validation performance under the random pair split and the unseen-drug split. The random pair split measures predictive accuracy on randomly sampled drug-side effect associations, whereas the unseen-drug split provides a more stringent assessment of generalization to previously unseen drugs. The final score assigns a higher weight to the unseen-drug split to prioritize cross-drug generalization during architecture search.

Within each validation split, model performance was evaluated from both classification and regression perspectives. AUROC and AUPRC were used for classification, whereas RMSE and MAE were used for quantitative side-effect frequency prediction. Because RMSE and MAE are minimization objectives, they were converted into bounded utility scores:

$$u_{\text{RMSE}} = \frac{1}{1 + x_{\text{RMSE}}} \tag{57}$$

$$u_{\text{MAE}} = \frac{1}{1 + x_{\text{MAE}}} \tag{58}$$

The score for each validation split was defined as

$$S_{\text{split}} = 10(0.35x_{\text{AUROC}} + 0.35x_{\text{AUPRC}} + 0.15u_{\text{RMSE}} + 0.15u_{\text{MAE}}) \tag{59}$$

The larger weights on AUROC and AUPRC preserve optimization pressure on robust discrimination, while the RMSE and MAE utilities maintain sensitivity to quantitative frequency prediction. The final side-effect prediction score was computed as

$$S = 0.30S_{\text{pair}} + 0.70S_{\text{drug}} \tag{60}$$

where $S_{\text{pair}}$ and $S_{\text{drug}}$ denote the validation scores under the random pair split and unseen-drug split, respectively. By assigning 70% of the overall score to the unseen-drug split, DrugEvolve explicitly biases the evolutionary search toward architectures with stronger generalization to unseen compounds while balancing classification accuracy and quantitative side-effect prediction performance.

**Method S3**. Prompting strategies for evolutionary search

To investigate how architectural constraints affect evolutionary search, we designed two implementation prompting schemes: a framework-constrained prompt and a framework-free prompt. The two prompts retained the same task objective and output requirements but differed in whether the supplied baseline implementation was treated as an architectural scaffold. A comparative analysis of their effects on evolutionary performance and design diversity is provided in **Supplementary Discussion S1**. The conceptual designs of the two prompting schemes are illustrated below using protein druggable-site annotation as a representative task.

##### S3.1 Framework-constrained implementation prompt

| Prompt 1. Framework-constrained implementation prompt |
| --- |
| Your task is to implement the proposed algorithm for protein druggable site annotation while preserving full compatibility with the existing training and evaluation pipeline.<br>A baseline implementation is provided below to define the required public interfaces, method signatures, input/output behavior, checkpoint handling, and framework-visible execution semantics.<br>For this implementation setting, the supplied code should be treated as both an **interface template and an architectural scaffold**. The high-level organization and principal computational flow of the existing `Encoder`, `Decoder`, `Predictor`, `Trainer`, and `Tester` components should remain recognizable. The proposed algorithm should therefore be implemented primarily by modifying the internal computations of these components rather than by replacing the overall architecture.<br>In particular:<br>• The overall module organization should remain recognizable.<br>• The principal data flow should remain compatible with the supplied architecture; for example, protein-level representations should be processed by an encoder-like component, local features should be processed by a decoder- or classifier-like component, and final predictions should be produced through the `Predictor`.<br>• You may modify layer types, feature transformations, attention mechanisms, pooling strategies, fusion operations, and loss computation within the existing components.<br>• You may introduce auxiliary private functions or submodules when necessary.<br>• You should not remove or fundamentally bypass the principal roles of the existing architectural components.<br>All framework-visible public interfaces, method signatures, input/output formats, tensor shapes, and invocation semantics must remain unchanged. The implementation must remain compatible with model construction, training-time loss computation, evaluation-time prediction, checkpoint saving and loading, and downstream metric aggregation.<br>In short, the supplied implementation serves as both a compatibility template and an architectural scaffold. The proposed algorithm should be realized through controlled modifications within this scaffold rather than through a complete architectural redesign. |

##### S3.2 Framework-free implementation prompt

##### Prompt 2. Framework-free implementation prompt

Your task is to implement the proposed algorithm for protein druggable site annotation while preserving full compatibility with the existing training and evaluation pipeline.

A baseline implementation is provided below to define the required public interfaces, method signatures, input/output behavior, checkpoint handling, and framework-visible execution semantics.

For this implementation setting, the supplied code should be treated only as an **interface and compatibility template**, rather than as an architectural scaffold. The internal organization and computational flow of the existing `Encoder`, `Decoder`, `Predictor`, `Trainer`, and `Tester` components do not need to be preserved. The proposed algorithm may therefore be implemented through substantial architectural redesign, provided that all framework-visible interfaces and input/output contracts remain unchanged.

In particular:

- You may redesign the overall model organization and computational flow.
- You may replace, merge, or remove internal architectural components.
- You may introduce a fundamentally different modeling paradigm when appropriate.
- You may redesign how protein context is represented, how local or residue-centered evidence is constructed, and how protein-wide information is aggregated.
- You may redesign how local and global representations are fused, how predictions are generated, and how the loss function is formulated.
- Components named `Encoder` or `Decoder` do not need to be retained internally unless required for framework compatibility.

All framework-visible public interfaces, method signatures, input/output formats, tensor shapes, and invocation semantics must remain unchanged. The implementation must remain compatible with model construction, training-time loss computation, evaluation-time prediction, checkpoint saving and loading, and downstream metric aggregation.

In short, the supplied implementation serves only as a compatibility template and does not restrict the internal modeling choices.

**Method S4**. Evolutionary candidate sampling strategy

##### S4.1 Task-specific sampling design and notation

DrugEvolve maintains an *Evolutionary Database* containing the complete history of autonomous algorithm development. For a drug-development task $q$, the database after evolutionary iteration $t$ is denoted by

$$\mathcal{D}_t^{(q)} = \left\{x_1^{(q)}, x_2^{(q)}, \dots, x_{N_t}^{(q)}\right\} \quad (61)$$

where $x_i^{(q)}$ represents a completed evolutionary candidate. Each candidate record contains

$$x_i^{(q)} = (p_i, g_i, \mathbf{m}_i, F_i, a_i, \ell_i, n_i) \quad (62)$$

where $p_i$ is the algorithmic proposal and scientific rationale, $g_i$ is the executable program, $\mathbf{m}_i$ contains task-specific evaluation metrics, $F_i$ is the fitness score, $a_i$ is the analysis report, $\ell_i$ records the evolutionary lineage, and $n_i$ is the number of previous retrieval events associated with the candidate.

DrugEvolve provides two alternative sampling strategies: the upper confidence bound (UCB)1-based sampling[7] and island-based Multi-dimensional Archive of Phenotypic Elites (MAP-Elites)-style sampling[8]. For each task $q$, a single sampling strategy

$$P_q \in \{\text{UCB1}, \text{Island}\} \quad (63)$$

is selected before evolution according to the task objective, evaluation budget, and expected heterogeneity of the algorithmic search space. The selected strategy remains fixed throughout the corresponding evolutionary run. In this study, Island-based MAP-Elites sampling was used for protein druggable-site annotation, whereas UCB1-based sampling was used for all other tasks. At the beginning of an evolutionary iteration, the task-specific sampler retrieves a candidate set

$$\mathcal{S}_t^{(q)} = \text{Sample}_{P_q}\left(\mathcal{D}_t^{(q)}; \Theta_{\text{sample}}^{(q)}\right) \quad (64)$$

The retrieved set contains a parent candidate to be modified and, when required, additional reference candidates that provide successful and unsuccessful evolutionary experience to the *Researcher*.

##### S4.2 Task-specific evolutionary search pipeline

**Algorithm S1** summarizes the task-level assignment and execution of evolutionary search in DrugEvolve. For each drug-development task $q$, the task-specific evaluation function $\text{Evaluate}_q(\cdot)$ executes a candidate algorithm under the corresponding training and validation protocol, and returns the validation results $\mathbf{m}_i^{(q)}$, including the raw metrics required by that task. The validation results are then passed to a task-specific scoring function, denoted as $\mathrm{Score_q}(\cdot)$, which converts the raw validation metrics into a scalar evolutionary fitness value:

$$F_q(x_i) = \mathrm{Score_q}\left(\mathbf{m}_i^{(q)}\right) \tag{65}$$

DrugEvolve does not apply a single universal scoring equation across all tasks. Instead, each task uses a predefined fitness function tailored to its evaluation criteria and optimization goal. Detailed scoring functions for all tasks are described in **Supplementary Method S2**.

For each task, DrugEvolve initializes the evolutionary database with the baseline algorithm, evaluates the baseline using the corresponding task protocol, converts the resulting metrics into a scalar fitness score, and then applies the preassigned sampling policy throughout the evolutionary run. **Algorithm S1** summarizes this task-level workflow.

---

**Algorithm S1** Task-Specific Evolutionary Search Pipeline

---

**Input**:
Task $q$
Baseline algorithm $x_0$
Preassigned sampling policy $P_q$
Task-specific evaluation function $\mathrm{Evaluate}_q(\cdot)$
Evolutionary budget $T_q$
Sampler-specific hyperparameters $\Theta_q$
**Output**:
Best evolved algorithm $x_{best}$
Task-specific Evolutionary Database $D_q$

$D_q$ ← InitializeDatabase($x_0$)
$\mathbf{m_0}$ ←$\mathrm{Evaluate}_q(x_0)$
$F_q(x_0) = \mathrm{Score}(\mathbf{m_0})$
$\mathrm{H}_\mathbf{0}$ ← $\mathrm{Analyze}_q(x_0, \mathbf{m}_0)$
$\mathrm{Store}\left(D_q, x_0, \mathbf{m_0}, F_q(x_0), \mathrm{H}_\mathbf{0}\right)$

$P$ ← $P_q$
**for** $t$ = 1 to $T_q$ **do**
  **if** $P$ = UCB1 **then**
    $\mathcal{C}_t^{(q)}$ ← SampleUCB1($\mathcal{D}_t^{(q)}$, $\Theta_q$)
  **else if** $P$ = Island-MAP-Elites **then**
    $\mathcal{C}_t^{(q)}$ ← SampleIslandArchive($\mathcal{D}_t^{(q)}$, $\Theta_q$)
  **end if**
  $D_q$ ← EvolveAndEvaluateCandidates($q$, $\mathcal{C}_t^{(q)}$, $D_q$)
**end for**
$x_{best}$ ← GetBestAlgorithm($D_q$)

16: **Return** $x_{best}$, $D_q$

##### S4.3 UCB1-based evolutionary candidate sampling

UCB1 is an uncertainty-aware selection strategy that balances exploitation of candidates with strong observed utility and exploration of candidates that have been selected less frequently. It may be particularly suitable when candidates can be ranked using a comparable scalar selection utility and the search is expected to benefit from progressively refining promising evolutionary lineages while retaining opportunities to revisit underexplored alternatives. At each sampling step, candidates that have never been retrieved are given priority. If the number of unvisited candidates is sufficient to satisfy the requested sample size, candidates are selected uniformly at random and without replacement from this unvisited subset. Otherwise, all candidates are ranked using a UCB1 score that combines normalized selection utility with an exploration bonus:

$$U_i(t) = \hat{F}_i(t) + c_q\sqrt{\frac{\ln\left(1 + N_{\text{total}}(t)\right)}{1 + n_i(t)}} \tag{66}$$

where, at step $t$, $\hat{F}_i(t)$ is the min-max normalized selection utility, $n_i(t)$ is the number of previous retrieval events associated with candidate $i$, $c_q$ is the task-specific exploration coefficient, and

$$N_{\text{total}}(t) = \sum_{j=1}^{N_t} n_j\,(t) \tag{67}$$

is the total number of retrieval events across all available candidates. The fitness term favors candidates with strong empirical performance, whereas the confidence term gives a larger exploration bonus to candidates that have been retrieved less frequently. The first term favors candidates with stronger empirical utility, whereas the second assigns a larger exploration bonus to candidates that have been retrieved less frequently. The candidates with the highest UCB1 scores are selected without replacement to form the retrieved candidate set. Their retrieval counts are subsequently updated for use in later sampling steps. The designation of one retrieved candidate as the evolutionary parent and the use of the remaining candidates as contextual references are performed outside the UCB1 sampler, as summarized in **Algorithm S2**.

**Algorithm S2** UCB1-Based Evolutionary Candidate Sampling

**Input**:

Evolutionary candidate set $\mathcal{D}_t^{(q)}$
Number of candidates requested $K_q$
Exploration coefficient $c_q$
Selection utility $F_q^{\text{sel}}(x_i)$ and previous-selection count $n_i(t)$

**Output**:
Selected candidate set $\mathcal{C}_t^{(q)}$

**def** SampleUCB1($\mathcal{D}_t^{(q)}$, $K_q$, $c_q$):
  **if** $\mathcal{D}_t^{(q)}$ is empty **then**
    **return** $\varnothing$
  **end if**
  *# Do not request more candidates than are currently available.*
  $K \leftarrow \min\left(K_q, \left|\mathcal{D}_t^{(q)}\right|\right)$
  *# Summarize previous sampling activity and identify candidates that have never been selected.*
  $N_{\text{visit}}(t) \leftarrow \sum_{x_j \in \mathcal{D}_t^{(q)}} n_j\,(t)$
  $\mathcal{U}_0 \leftarrow \left\{x_i \in \mathcal{D}_t^{(q)} : n_i(t) = 0\right\}$
  *# When enough unvisited candidates exist, sample only from them to guarantee initial exploration.*
  **if** $|\mathcal{U}_0| \geq K$ **then**
    $\mathcal{C}_t^{(q)} \leftarrow \text{UniformSampleWithoutReplacement}(\mathcal{U}_0, K)$
    $\text{MarkSelected}\left(\mathcal{C}_t^{(q)}\right)$
    **return** $\mathcal{C}_t^{(q)}$
  **else**
    *# Otherwise, place utilities on a common 0-1 scale before combining quality and exploration.*
    $F_{\min} \leftarrow \text{MinimumSelectionUtility}(\mathcal{D}_t^{(q)})$
    $F_{\max} \leftarrow \text{MaximumSelectionUtility}(\mathcal{D}_t^{(q)})$
    $\Delta_F \leftarrow F_{\max} - F_{\min}$ ; use 1 when all utilities are identical.
    **for** each candidate $x_i \in \mathcal{D}_t^{(q)}$ **do**
      **if** $n_i(t) = 0$ **then**
        $U_i(t) \leftarrow +\infty$
        *# An infinite score ensures that any remaining unvisited candidate is ranked first.*
      **else**
        Compute the normalized selection utility $\hat{F}_i$
        Compute the exploration bonus $B_i(t)$
        $U_i(t) \leftarrow \hat{F}_i + B_i(t)$
      **end if**
    **end for**
    $\mathcal{C}_t^{(q)} \leftarrow \text{TopKWithoutReplacement}\left(\mathcal{D}_t^{(q)}, U_i(t), K\right)$
  **end if**
  *# Rank all candidates by UCB1 score and return the highest-scoring unique candidates.*

MarkSelected($\mathcal{C}_t^{(q)}$)
**return** $\mathcal{C}_t^{(q)}$

---

##### S4.4 Island-based evolutionary candidate sampling

An island- and archive-based sampler is a diversity-aware selection strategy that combines rotating emphasis on different subpopulations with a Multi-dimensional Archive of Phenotypic Elites (MAP-Elites)-style archive. MAP-Elites is a quality-diversity approach that partitions candidates into behavioral niches according to predefined descriptors and retains the highest-performing candidate, referred to as the elite, within each occupied niche[8]. Rather than preserving only a single globally best candidate, this mechanism maintains multiple high-quality candidates representing different regions of the behavioral space. This sampling strategy may be particularly suitable when the algorithmic search space contains heterogeneous or mechanistically distinct solution families and the search is expected to benefit from preserving multiple promising design directions.

For each candidate, predefined behavioral features, such as algorithmic complexity and design diversity, are normalized using persistently updated feature ranges and discretized to define a niche coordinate. A shared global archive retains the candidate with the highest selection utility observed in each occupied niche. Consequently, a candidate may remain evolutionarily valuable even when it does not achieve the highest utility across the entire population, provided that it is the strongest candidate within a distinct behavioral niche. At each sampling step, candidates assigned to the current island form the preferred sampling pool. If the current island contains no candidates, or if its unselected candidates are exhausted before the requested sample size is reached, the sampler falls back to the globally available candidate set. Candidates are selected without replacement through a mixture of uniform exploration, global archive-based exploitation, and utility-weighted sampling. The active island advances in round-robin order after each sampling call. No separate periodic migration operation is implemented; cross-island information sharing instead occurs through both the shared global archive, which allows archived elites from any island to contribute during exploitation, and the global fallback mechanism used when the current-island pool cannot complete the requested sample. The complete island-based candidate-sampling procedure is described in **Algorithm S3**.

---

**Algorithm S3** Island-Based Sampling

---

**Input**:

Evolutionary candidate set $\mathcal{D}_t^{(q)}$

Number of candidates requested $K_q$ and number of islands $M_q$
Current-island state $m_t$ and global niche archive $\mathcal{A}$
Feature dimensions $\mathcal{G}_q$, number of bins $B_q$, and persistent feature statistics
Exploration and exploitation probabilities $p_{\text{explore}}$ and $p_{\text{exploit}}$
Selection utility $F_q^{\text{sel}}(x_i)$ and small positive constant $\varepsilon = 10^{-6}$

**Output**:
Selected candidate set $\mathcal{C}_t^{(q)}$
Current-island state updated in place

**def** FeatureCoordinates($x$):
    *# Convert configured behavioral features into a discrete niche coordinate.*
    **if** $\mathcal{B}_q = \varnothing$ **then**
        **return None**
    **end if**
    $\mathbf{z} \leftarrow [\,]$
    **for each** $b \in \mathcal{B}_q$ **do**
        $v \leftarrow$ FeatureValue$(x, b)$
        **if** $v \notin \mathbb{R}$ **then return None**
        $h_b^{\min} \leftarrow \min\left(h_b^{\min}, v\right)$; $h_b^{\max} \leftarrow \max(h_b^{\max}, v)$
        $\Delta_b \leftarrow h_b^{\max} - h_b^{\min}$
        $\alpha_b \leftarrow 0.5$ **if** $\Delta_b \le 10^{-12}$, otherwise $\alpha_b \leftarrow \left(v - h_b^{\min}\right)/\Delta_b$
        $z_b \leftarrow \min\left(L_q - 1, \max\left(0, \lfloor \alpha_b L_q \rfloor\right)\right)$; **append** $z_b$ to $\mathbf{z}$
    **end for**
    **return** $\mathbf{z}$

**def** RegisterCandidate($x$):
    *# Update the shared niche archive and assign an island when the node is added.*
    $\mathbf{z}(x) \leftarrow$ FeatureValue$(x)$
    **if** $\mathbf{z}(x)$ is not None **then**
        store $\mathbf{z}(x)$ as the niche metadata of $x$
            **if** $\mathcal{A}_t[\mathbf{z}(x)] = \varnothing$ or $F_q^{\text{sel}}(x) > \text{Utility}(\mathcal{A}_t[\mathbf{z}(x)])$ **then**
                $\mathcal{A}_t[\mathbf{z}(x)] \leftarrow \left(\text{id}(x), F_q^{\text{sel}}(x)\right)$
            **end if**
    **end if**
    **if** $m(x)$ **is undefined then**
        **if** $\mathbf{z}(x)$ is not None **then** $m(x) \leftarrow \mathit{Hash}32\left(\mathbf{z}(x)\right) \bmod M_q$
        **else if** $x$ **has a parent then** $m(x) \leftarrow m_t$
        **else** $m(x) \leftarrow (\text{id}(x)$ if defined, otherwise $0) \bmod M_q$
    **end if**

**def** SampleIslandArchive($\mathcal{D}_t^{(q)}, K_q$):

  **if** $\mathcal{D}_t^{(q)}$ is empty **then**

    **return** $\varnothing$

  **end if**

  *# Prefer the current island, but fall back to all available candidates when needed.*

  $\mathcal{P} \leftarrow \{x \in \mathcal{D}_t^{(q)}: m(x) = m_t\}$

  **if** $\mathcal{P} = \varnothing$ **then**

    $\mathcal{P} \leftarrow \mathcal{D}_t^{(q)}$

  **end if**

  $K_q \leftarrow \min(K_q, |\mathcal{D}_t^{(q)}|)$

  $\mathcal{C}_t^{(q)} \leftarrow \varnothing$

  *# Accumulate unique candidates until the requested sample size is reached.*

  **while** $|\mathcal{C}_t^{(q)}| < K_q$ **do**

    $\mathcal{G} \leftarrow \mathcal{D}_t^{(q)} \setminus \mathcal{C}_t^{(q)}$; **if** $\mathcal{G} = \varnothing$ **then break**

    $\mathcal{I} \leftarrow \mathcal{P} \setminus \mathcal{C}_t^{(q)}$; $\mathcal{Q} \leftarrow \mathcal{I}$ **if** $\mathcal{I} \neq \varnothing$, otherwise $\mathcal{G}$

    sample $r \sim \mathcal{U}(0,1)$

    *# Exploration gives every candidate in the active pool equal probability.*

    **if** $r < p_{\text{explore}}$ **then** $x^* \sim \text{Uniform}(\mathcal{Q})$

    **else if** $r < p_{\text{explore}} + p_{\text{exploit}}$ **then**

    *# Archive-based exploitation is global rather than restricted to the current island.*

      $\mathcal{E} \leftarrow \{x \in \mathcal{G}: \text{id}(x) \in \text{ArchiveIDs}(\mathcal{A}_t)\}$

      **if** $\mathcal{E} \neq \varnothing$ **then** $x^* \sim \text{Uniform}(\mathcal{E})$

      **else** $x^* \leftarrow \text{argmax}_{x \in \mathcal{G}} F_q^{\text{sel}}(x)$

    **else**

    *# Utility shifting preserves relative preferences while preventing zero or negative sampling weights.*

      $F_{\min} \leftarrow \min_{y \in \mathcal{Q}} F_q^{\text{sel}}(y)$

      $w(y) \leftarrow F_q^{\text{sel}}(y) - F_{\min} + \varepsilon$

      sample $x^*$ from $\mathcal{Q}$ with probability $w(x^*) / \sum_{y \in \mathcal{Q}} w(y)$

    **end if**

    $\mathcal{C}_t^{(q)} \leftarrow \mathcal{C}_t^{(q)} \cup \{x^*\}$

  **end while**

  *# Rotate island emphasis and record the retrieval events.*

  $m_{t+1} \leftarrow (m_t + 1) \bmod M_q$; update state["current_island"]

  MarkSelected$\left(\mathcal{C}_t^{(q)}\right)$

  **return** $\mathcal{C}_t^{(q)}$

**Method S5**. Detailed descriptions of the datasets used across all drug-development tasks

#### S5.1 Target Identification

##### *S5.1.1 Cancer gene module detection*

Following the experimental protocol of CGMega, we evaluated the model on the MCF-7 human breast cancer cell line using a curated multi-omics dataset comprising genomic, epigenomic and high-throughput chromosome conformation capture (Hi-C)-derived features together with protein-protein interaction networks[9]. The training-validation data were divided into ten folds, with nine folds used for training and the remaining fold for validation in each round, while an independent test set was kept fixed across all rounds. Detailed statistics of the dataset composition and class distribution are summarized in **Supplementary Table S8**.

##### *S5.1.2 Gene-disease association prediction*

Following the experimental protocol of FusionGDA, we adopted the DisGeNET-EVAL benchmark introduced by the prior study[2]. DisGeNET-EVAL was constructed from DisGeNET[10] (version 2023), which contains 1,041,587 experimentally curated and literature-supported gene-disease associations involving 16,622 genes and 28,873 diseases. Protein sequences were retrieved from the STRING database[11] by matching gene symbols, whereas disease descriptions were obtained from the Medical Genome Database Extended Form (MGDEF)[12] using disease names. For evaluation, an equal number of negative gene-disease pairs were randomly sampled without overlapping known associations. Predictive performance was assessed using five independent random data partitions, where the model was independently trained and evaluated on each split. In addition to the DisGeNET-based benchmarks, we included the Therapeutics Data Commons (TDC)[13] GDA dataset as an external evaluation set. TDC provides a standardized binary classification setting with a 1:1 ratio of positive and negative gene-disease pairs, where negative samples are randomly generated based on positive associations.

##### *S5.1.3 Protein druggable site annotation*

**Protein-protein interaction (PPI) site.** The PPI site benchmark suite comprises four widely used datasets: PPI-Train352/PPI-Test70 (DeepPPISP[14]), PPI-Train9982/PPI-Test355 (DELPHI[15]), PPI-Train335/PPI-Test60 (GraphPPIS[16]), and an additional independent test set, PPI-Test315. All datasets are constructed under strict non-redundancy criteria, with sequence identity typically filtered to no more than 25%. PPI-Train352 and PPI-Test70 are derived from DeepPPISP[14], where binding residues are annotated based on reductions in solvent accessibility upon complex formation. PPI-Train9982/PPI-Test355 are curated from BioLip[17], in which binding residues are defined according to atomic distance criteria; a subset of the training data is used for validation, and structural annotations in the training split are partially incomplete. The GraphPPIS dataset follows the same sequence identity filtering protocol and provides an additional external evaluation setting for benchmarking.

**Peptide-protein interaction (PepPI) site.** PepPI-Train1154/PepPI-Test125 was obtained from from SPRINT-Str[18], whereas PepPI-Train640/PepPI-Test639 is derived from a previously published study[19]. Both datasets follow standard preprocessing pipelines, with sequence identity reduced to no more than 30% using BLAST-based clustering to prevent redundancy between training and test sets, thereby ensuring a reliable evaluation of peptide-protein interaction site prediction performance[20].

**DNA-protein interaction (DPI) site.** The DPI benchmarks include DPI-Train573/DPI-Test129 from GraphBind[21], together with an additional independent test set, DPI-Test181. All datasets are derived from BioLip and filtered to ensure sequence identity of no more than 30% between training and test sets. Binding residues are defined based on atomic proximity between DNA molecules and protein residues using van der Waals distance criteria[22]. DPI-Test181 serves as an external evaluation set to assess model generalization performance.

**RNA-protein interaction (RPI) site.** The RPI benchmark, also introduced in GraphBind[21], consists of 495 training proteins and 117 test proteins. Data preprocessing procedures and binding-site definitions are consistent with those used in the DPI benchmarks. Sequence identity is strictly constrained to below 30% to reduce redundancy. A pronounced class imbalance is observed, in which non-binding residues substantially outnumber binding residues.

**Small molecule-protein (SMPI) site.** The SMPI benchmark is constructed from the sc-PDB database[23], initially comprising 17,594 protein-ligand complexes. After mapping structures to UniProt sequences and removing redundancy at a sequence identity threshold of no more than 30%, a non-redundant set of 2,324 proteins is obtained. The dataset is partitioned into 1,628 training, 348 validation and 348 test proteins. Binding residues are defined as residues having any atom within 6.5 Å of a ligand atom.

**Carbohydrate-protein interaction (CarbPI) site.** The CarbPI dataset is derived from the CAPSIF study[24] and consists of 517 training, 129 validation, and 162 test proteins. All structures have a resolution better than 3.0 Å, and sequence identity between any pair of proteins is constrained to below 30%. Binding residues are defined as residues having any heavy atom within 4.2 Å of a carbohydrate atom.

**Ion-binding site.** The LigBind datasets were constructed in a previous study[25], comprising ligand-specific datasets, pre-training datasets, and unseen-ligand test sets derived from the February 2021 release of the BioLiP database[17]. CD-HIT was applied to remove redundant protein sequences and to strictly control sequence identity between training and test protein chains to below 30%[26]. Binding residues were defined according to atomic proximity between ligands and protein residues. The unseen-ligand test set, which contains 16 novel ligands, serves as an external evaluation set to assess model generalization performance to previously unseen ligand types[27]. The GPSites benchmark datasets were derived from LABind[27], which retains six target ligands, including ATP, HEM, $Zn^{2+}$, $Ca^{2+}$, $Mg^{2+}$, and $Mn^{2+}$. A strict chronological data splitting strategy was adopted, in which proteins released before January 1, 2021 for small molecules and before January 1, 2020 for metal ions were allocated to the training sets. Entries released after these cut-off dates were used to construct independent test sets. This procedure was further constrained by the same requirement as in the LigBind benchmark, namely that all ligands appearing in the test sets must also have been present during the training phase.

**Covalent site.** To construct datasets for training and validating the covalent binding site prediction task, CovalentInDB 2.0 was adopted as the primary data source[28]. After removing redundant protein entries, the remaining unique records were randomly partitioned to support robust model training and evaluation. A curated set of approved covalent drugs was further analyzed to characterize the physicochemical and interaction properties underlying established covalent binding patterns. In addition, the PDBbind v2020 dataset, which comprises 19,443 PDB entries, was employed to investigate the distribution and prevalence of covalent-binding residues across diverse protein binding pockets[29]. Finally, representative protein targets and high-quality protein-ligand complexes were systematically selected to construct a large-scale molecular library, thereby enabling rigorous model validation and providing a valuable resource for downstream covalent drug development. The Cys dataset was constructed from CovalentInDB 2.0[28]. Protein sequences and covalently ligand-bound cysteine sites were extracted from the corresponding PDB entries, followed by sequence consolidation and redundancy removal using MMseqs2 clustering (30% sequence identity, 80% coverage). The resulting non-redundant dataset contained 234 representative protein sequences with 1,302 cysteine residues, including 238 positive sites annotated as covalently ligand-bound cysteines and 1,064 negative sites without covalent ligand-binding annotations.

**Catalytic site.** Uni14230 and Uni3175 were utilised as described in prior work[30]. These datasets were originally constructed from UniProt[31], M-CSA[32], and SwissProt[32] annotations, and include both manually curated and computationally predicted catalytic residue labels. To reduce sequence redundancy, sequences sharing more than 60% identity were removed. In addition, six benchmark datasets reported in earlier studies were used for external evaluation[33-36]. These datasets include family-, superfamily-, and fold-level enzyme benchmarks, as well as the NN and PC datasets, which were designed to cover structurally and functionally diverse enzymes based on SCOP and CATH classifications[35,36]. To address the limitations of existing benchmark datasets for evaluating catalytic residue prediction, a prior study[37] introduced a benchmark dataset named CataloDB. This dataset was constructed based on enzyme annotations in SwissProt and includes experimentally supported catalytic residue annotations as well as reviewed enzyme sequences with resolved structures. Redundant sequences sharing more than 90% sequence identity were removed using MMseqs2[38]. Sequence-based and structure-based clustering were subsequently applied to define a stringent test set. The resulting test set was further constrained to ensure that both sequence and structural identity relative to the training set were below 30%, with structural similarity assessed using Foldseek[39]. In addition, sequences overlapping with six commonly used benchmark datasets were excluded from the CataloDB training set to ensure consistency with prior evaluation settings.

**Cryptic site.** The CryptoSites dataset used in this study was collected from CryptoBank[40] and consists exclusively of cryptic holo protein structures. To ensure adequate structural complexity, protein sequences shorter than 50 amino acids were excluded. Binding sites were defined as residues containing at least one atom within 5 Å of any atom of a cryptic ligand, and these annotations were converted into binary residue-level labels for subsequent classification tasks. For protein chains associated with multiple ligand annotations, labels were integrated using a residue-wise maximum operation to generate a single binding-site annotation for each chain. To prevent sequence identity leakage between data partitions, protein sequences were clustered using MMseqs2[38] with a minimum sequence identity threshold of 20%. The independent test set was constructed by sequentially selecting the smallest connected components until approximately 10% of the remaining sequence pool had been allocated. The validation set was subsequently generated by randomly sampling connected components from the remaining pool until another approximately 10% had been assigned, whereas all remaining components were used for training. This component-based partitioning strategy ensured that no sequence in the test set shared more than 20% sequence identity with any sequence in the training or validation sets. The final dataset comprised 7,931 protein sequences, including 6,345 sequences in the training set and 793 sequences each in the validation and independent test sets.

**Post-translational modification (PTM) site**. The PTM dataset was obtained from DeepMVP[41], which curated experimentally validated post-translational modification annotations from PTMAtlas to establish residue-level prediction tasks for major PTM types. The dataset includes phosphorylation, methylation, N-glycosylation, acetylation, ubiquitination, and sumoylation, with candidate residues defined according to the biochemical specificity of each modification. Specifically, phosphorylation sites are restricted to serine, threonine, and tyrosine residues; methylation sites to lysine and arginine residues; N-glycosylation sites to asparagine residues; and acetylation, ubiquitination, and sumoylation sites to lysine residues. For each PTM type, experimentally validated modification sites annotated in PTMAtlas are treated as positive samples. Negative samples are selected from the corresponding candidate residue types within the same proteins after removing all known PTM sites recorded in PTMAtlas, UniProt[42], PhosphoSitePlus (PSP)[43], and PLMD[44]. The complete dataset is partitioned into training, validation, and independent test sets, comprising 81%, 9%, and 10% of the samples, respectively. To further investigate the potential impact of sequence similarity between training and test proteins on model generalization, an additional protein-level re-splitting protocol is employed. Under this protocol, test peptides exhibiting sequence similarity greater than 90%, 80%, or 70% to any peptide in the training set are progressively excluded. This strategy enables a systematic assessment of model performance under increasingly stringent sequence similarity constraints and provides a more rigorous evaluation of generalization capability.

Detailed statistics for all benchmark datasets used in protein druggable site annotation, including the numbers of proteins, residues, binding residues, non-binding residues, and the proportion of binding residues in each data split, are summarized in **Supplementary Table S9-14**.

##### S5.2 Drug Discovery

###### *S5.2.1 Molecular docking*

**PDBbind.** Following the experimental protocol of the prior study[3], we adopted its temporally split PDBbind v2020 benchmark[29], in which complexes released in 2019 or later were reserved for testing and earlier complexes were used for training and validation. We then independently processed the downloaded protein-ligand complexes for EquiBind by excluding ligands that could not be parsed by RDKit[45], preparing receptor structures with OpenBabel[46], and retaining protein chains containing atoms within 10 Å of the ligand; when no chain met this criterion, the closest valid protein chain was retained. After preprocessing and removal of complexes that could not be successfully processed, the final dataset contained 16,379 training, 968 validation and 363 test complexes.

**Posebusters.** To assess performance on more recent and previously unseen systems, we additionally evaluated the models on the PoseBusters Benchmark Set[47]. This PDB-derived release contains 428 protein-ligand complexes and provides processed receptor structures, crystallographic ligand poses and independently generated ligand starting conformations, enabling evaluation of docking accuracy, physical plausibility and out-of-distribution generalization.

**Astex Diverse Set.** We used the Astex Diverse Set introduced by *Hartshorn et al.*[48], a curated benchmark derived from X-ray crystal structures in the Protein Data Bank[49]. The dataset comprises 85 diverse, high-quality protein-ligand complexes spanning distinct protein targets and drug-like ligands, with experimentally well-supported binding poses, and is widely used for evaluating protein-ligand docking performance.

##### *S5.2.2 Drug-target interaction prediction*

**BindingDB.** The BindingDB benchmark employed in this study was derived from a previous work[50] and has been widely used to evaluate computational methods for compound-protein interaction prediction. It contains 33,772 positive interaction pairs and 27,486 negative pairs collected from the BindingDB database[51]. The original benchmark provides predefined training, validation, and test partitions under three evaluation settings: *random-split, unseen-drug, and unseen-protein*. In the unseen-drug setting, all compounds appearing in the test set are excluded from the training set. Similarly, in the unseen-protein setting, all proteins in the test set are excluded from training. By eliminating entity-level overlap between the training and test sets, these evaluation protocols provide a more rigorous assessment of model generalization by requiring transferable interaction patterns rather than memorized entity representations.

**Human-cold.** The Human dataset, originally constructed by *Tsubaki et al.*[52], comprises positive drug-target interaction pairs collected from DrugBank[53] 4.1 and Matador[54], together with high-confidence negative samples generated through a systematic screening procedure. As noted in TransformerCPI[50], the dataset may be affected by hidden ligand bias, whereby predictive performance is driven largely by compound-specific characteristics rather than genuine drug-target interaction patterns. As a result, evaluation based on conventional random splits can yield overly optimistic estimates and may not accurately reflect prospective predictive performance. To mitigate this issue, a cold-pair splitting strategy was adopted in DrugBAN[55]. Specifically, 5% and 10% of drug-target interaction pairs were randomly allocated to the validation and test sets, respectively, and all drugs and proteins associated with these pairs were excluded from the training set. Consequently, neither the drugs nor the proteins contained in the validation and test sets were observed during training, preventing models from relying on memorized representations of previously seen compounds or targets. Dataset statistics and split composition for BindingDB and Human-cold are summarized in **Supplementary Table S15**.

###### *S5.2.3 De novo molecular generation*

**Molecular Sets (MOSES)**: We used the Molecular Sets (MOSES) benchmark dataset introduced by *Polykovskiy et al.*[56], which was constructed from the ZINC Clean Leads collection[57]. The original compounds were filtered to retain lead-like molecules with restricted molecular weight, lipophilicity, rotatable-bond count, atom types and ring sizes, followed by medicinal chemistry and PAINS filtering[58]. MOSES provides 1,584,664 molecules for training, 176,075 molecules for random testing and 176,226 molecules for scaffold-based testing. The scaffold-based test set (TestSF) contains Bemis-Murcko[59] scaffolds absent from the training set, enabling evaluation of generalization to unseen molecular frameworks. Molecular structures were represented as SMILES strings.

###### *S5.2.4 Target-specific peptide binder design*

**PepBench.** PepBench was assembled as a literature-derived, non-redundant benchmark for evaluating peptide binder design on unseen targets[60]. To minimize homology bias and prevent information leakage between the benchmark and the model development sets, complexes whose target proteins shared more than 40% sequence identity with targets in the training or validation sets were excluded. Sequence identity was computed using MMseqs2[38]. To ensure high-quality sequence inputs for downstream modeling, we further removed entries containing unknown amino acid residues, while incorporating corresponding structural data from the PDB to enrich the complex annotations. The final dataset comprises 4,147 training complexes, 114 validation complexes, and 103 protein-peptide complexes in the test set.

**PepNN.** The PepNN benchmark dataset was derived from PepMLM[5], which was curated from the PepNN and Propedia databases[61,62]. To reduce redundancy across the two sources, duplicate entries were first removed. The remaining protein sequences were clustered using MMseqs2[38] with a sequence similarity threshold of 0.8. For complexes in which protein sequences were assigned to the same cluster and shared identical peptide binder sequences, only a single representative complex was retained. To further mitigate homology bias, an additional manual curation step was performed to remove protein sequences with high sequence identity above an 80% threshold. We further excluded entries containing unknown amino-acid residues to ensure sequence reliability. The final dataset comprised 7,148 protein-peptide complexes, which were divided into 5,585 complexes for training, 1,415 for validation, and 148 complexes for testing.

##### *S5.2.5 ADMET property prediction*

A total of 17 ADMET datasets were collected from the admetSAR 3.0 database[63], covering the five major ADMET categories: absorption, distribution, metabolism, excretion, and toxicity. The absorption endpoints included pKa, oral bioavailability (F50%, binarized using 50% as the threshold), and Caco-2 permeability. The distribution endpoints included OATP1B1 inhibition and OCT2 inhibition. The metabolism endpoints comprised CYP2D6 inhibition, CYP2C9 substrate, CYP2B6 inhibition, and human liver microsomal stability (HLM). The excretion endpoints included plasma clearance (CLp) and mean residence time (MRT). The toxicity endpoints included hERG liability, Ames mutagenicity, FDAMDD, mouse carcinogenicity, androgen receptor (AR) activity, and acute oral toxicity. For hERG, an IC50 threshold of 1 μM was used, with compounds above the threshold labeled as non-cardiotoxic and those below as potentially cardiotoxic. All datasets were split using a scaffold-based partitioning strategy based on Murcko scaffold decomposition. Molecular structures were standardized from SMILES representations, and canonical Murcko[59] scaffolds were extracted to define scaffold groups. Compounds sharing the same scaffold were assigned to the same group. Groups were then allocated to training, validation, and test sets in a greedy manner under an 8:1:1 ratio, ensuring that scaffold overlap across splits was eliminated. The final benchmark contained 12 classification datasets and 5 regression datasets. Detailed dataset statistics and split information are provided in **Supplementary Table S16-17**.

#### S5.3 Preclinical Study

##### *S5.3.1 Animal toxicology assessment*

We used a local clinical pathology dataset derived from the Open TG-GATEs[64] data used in AnimalGAN[65]. The dataset comprises 138 compounds, 1,637 compound-dose-time treatment conditions and 1,826 Mordred[66] molecular descriptors, together with 8,078 treated-animal records and 2,775 vehicle-control records in the official split file. To enable feature generation, we supplemented the compound annotations with canonical and isomeric SMILES retrieved from PubChem[67]. Model applicability was assessed using four compound-level partitions of the same dataset, each assigning approximately 110 compounds to training and 28 compounds to testing while retaining all treatment conditions of a given compound within a single partition: a random compound split (Random), a structure-similarity split that held out the compounds with the lowest median pairwise similarity for testing (Structure), a therapeutic-class split based on

Anatomical Therapeutic Chemical categories (ATC, https://www.whocc.no/atc_ddd_index/), and a temporal split separating earlier or non-drug-like compounds from later-approved drugs (Time).

#### S5.4 Clinical Trial

##### *S5.4.1 Drug-drug interaction prediction*

Following the benchmark and evaluation protocol of MeTDDI, we adopted the datasets released by MotifGT-DTI[68]. For the classification task, metabolic DDIs were collected from DrugBank[69] (version 5.1.8). Model performance was evaluated under two complementary generalization settings, namely the unseen-one-drug and unseen-two-drugs splits, following the original benchmark protocol. For the regression task, we used the pharmacokinetic benchmark released by MeTDDI, which was derived from U.S. Food and Drug Administration (FDA) drug labels[70]. The task predicts the area under the plasma concentration-time curve fold change (AUC FC) of the victim drug to quantify the pharmacokinetic effect of the perpetrator. Detailed statistics for each data split are provided in **Supplementary Table S18**.

##### *S5.4.2 Drug side-effect prediction*

We utilized the SIDER benchmark dataset, referred to as the SIDER dataset in this work, which was published with SDPred[71]. Its core drug-side effect frequency matrix was first derived from Side effect Resource (SIDER)[72] 4.1 database by *Galeano et al.*[73], then integrated with compound annotations extracted from STITCH[74] and DrugBank[69]. After removing drugs that could not be matched across databases, the final dataset contains 757 drugs, 994 side effects, and 37,366 observed drug side-effect frequency pairs. Side-effect frequencies are categorized into five ordinal levels ranging from very rare to very frequent. To support joint classification and quantitative frequency prediction, additional negative drug side-effect pairs were incorporated following the SDPred preprocessing protocol, yielding 74,774 labelled pairs in total. To comprehensively evaluate model generalization, we considered two complementary data splitting strategies. In the random split, drug side-effect pairs were randomly partitioned into training (52,341 pairs), validation (7,477 pairs), and test (14,956 pairs) sets with a ratio of 7:1:2, allowing the same drug to appear across different splits. In the unseen-drug split, drugs were partitioned with the same ratio, resulting in 53,548, 7,368, and 13,858 drug side-effect pairs in the training, validation, and test sets, respectively, while ensuring that all pairs associated with each drug were assigned to a single split. This setting provides a more stringent evaluation of model generalization to previously unseen drugs.

**Method S6**. Definitions of evaluation metrics

##### S6.1 Classification task

Model performance was comprehensively evaluated using a diverse set of classification metrics, including accuracy (ACC), precision, recall, specificity, area under the receiver operator characteristic curve (AUROC), area under the precision-recall curve (AUPRC), F1 score, maximum F1 score ($F_{max}$) and Matthews correlation coefficient (MCC). Together, these metrics characterize predictive performance from multiple perspectives, encompassing classification accuracy, ranking ability, class balance, overlap between predicted and observed labels. The mathematical expressions used for their calculation are presented below:

$$\mathrm{ACC} = \frac{\mathrm{TP} + \mathrm{TN}}{\mathrm{TP} + \mathrm{TN} + \mathrm{FP} + \mathrm{FN}} \tag{68}$$

$$\mathrm{Precision} = \frac{\mathrm{TP}}{\mathrm{TP} + \mathrm{FP}} \tag{69}$$

$$\mathrm{Recall} = \frac{\mathrm{TP}}{\mathrm{TP} + \mathrm{FN}} \tag{70}$$

$$\mathrm{Specificity} = \frac{TN}{TN + FP} \tag{71}$$

$$\mathrm{F1} = \frac{2 \times Precision \times Recall}{Precision + Recal} \tag{72}$$

$$\mathrm{F_{max}} = \max_{\theta} \frac{2 \times Precision_{\theta} \times Recall_{\theta}}{Precision_{\theta} + Recall_{\theta}} \tag{73}$$

$$\mathrm{MCC} = \frac{\mathrm{TP} \times \mathrm{TN} - \mathrm{FP} \times \mathrm{FN}}{\sqrt{(\mathrm{TP} + \mathrm{FP}) \times (\mathrm{TP} + \mathrm{FN}) \times (\mathrm{TN} + \mathrm{FP}) \times (\mathrm{TN} + \mathrm{FN})}} \tag{74}$$

where $TP$, $TN$, $FP$ and $FN$ correspond to the numbers of true positives, true negatives, false positives, and false negatives, respectively. $\theta$ denotes the classification threshold. $F_{max}$ is defined as the maximum F1 score obtained by scanning all possible decision thresholds, thereby providing a threshold-optimized measure of the best achievable balance between precision and recall.

##### S6.2 Regression task

**Mean Absolute Error (MAE).** To evaluate the average magnitude of prediction errors, we employed the mean absolute error (MAE) between the predicted and reference values. MAE quantifies prediction accuracy by calculating the mean of the absolute differences between paired observations. MAE assigns equal weight to all prediction errors without squaring the deviations, making it less sensitive to large errors and outliers while providing a direct measure of the average prediction error. Lower MAE values indicate closer agreement between predicted and reference values. The metric is defined as follows:

$$\mathrm{MAE} = \frac{1}{n}\sum_{i=1}^{n}|\hat{y}_i - y_i| \tag{75}$$

where $n$ denotes the total number of evaluated samples, $\hat{y}_i$ represents the predicted value of the $i$-th sample, and $y_i$ denotes the corresponding reference value.

**Mean Squared Error (MSE).** We employed the mean squared error (MSE) between the predicted and reference values. MSE measures the average of the squares of the errors, representing the average squared difference between the estimated values and the actual value. By squaring the errors, MSE heavily penalizes larger discrepancies, making it a highly effective and rigorous metric when avoiding large deviations is a critical priority. Lower MSE values indicate superior model accuracy. The metric is defined as follows:

$$\mathrm{MSE} = \frac{1}{n}\sum_{i=1}^{n}(\hat{y}_i - y_i)^2 \tag{76}$$

where $n$ denotes the total number of evaluated samples, $\hat{y}_i$ represents the predicted value of the $i$-th sample, and $y_i$ denotes the corresponding reference value.

**Root Mean Square Error (RMSE).** To evaluate the numerical accuracy of the predicted values, we employed the root mean square error (RMSE) between the predicted and reference values. RMSE quantifies the average magnitude of prediction errors by calculating the square root of the mean squared differences between paired observations. Because prediction errors are squared prior to averaging, RMSE assigns greater weight to larger deviations and is therefore particularly sensitive to substantial prediction errors and outliers. Lower RMSE values indicate closer agreement between predicted and reference values. The metric is defined as follows:

$$\mathrm{RMSE} = \sqrt{\frac{1}{n}\sum_{i=1}^{n}(\hat{y}_i - y_i)^2} \tag{77}$$

where $n$ denotes the total number of evaluated samples, $\hat{y}_i$ represents the predicted value of the $i$-th sample, and $y_i$ denotes the corresponding reference value.

**Pearson correlation coefficient ($R_p$).** To evaluate the consistency between predicted and reference values, we calculated the Pearson correlation coefficient $R_p$. In contrast to error-based metrics that primarily measure absolute numerical deviations, $R_p$ quantifies the strength and direction of the linear relationship between predicted and reference values. This metric assesses the extent to which the predicted values reproduce the variation patterns and linear trends of the reference values across samples. Higher $R_p$ values indicate stronger positive linear agreement between predictions and references. The metric is defined as follows:

$$R_p = \frac{\sum_{i=1}^{N}(\hat{y}_i - \bar{\hat{y}})(y_i - \bar{y})}{\sqrt{\sum_{i=1}^{N}(\hat{y}_i - \bar{\hat{y}})^2} \cdot \sqrt{\sum_{i=1}^{N}(y_i - \bar{y})^2}} \tag{78}$$

where $N$ denotes the total number of evaluated samples, $\hat{y}_i$ and $y_i$ represent the predicted and reference values of the $i$-th sample, respectively, and $\bar{\hat{y}}$ and $\bar{y}$ denote the mean values of the predicted and reference variables.

**Coefficient of determination ($R^2$).** To assess overall predictive performance and goodness of fit, we employed the coefficient of determination ($R^2$). $R^2$ evaluates how well the predicted values explain the variance of the observed data by comparing the residual variance with the total variance of the observations. Consequently, larger prediction errors, including systematic bias, reduce the $R^2$ score. The value ranges from negative infinity to 1.0, where 1.0 indicates perfect agreement between predicted and observed values, whereas values below 0 indicate that the model performs worse than a naive predictor that always outputs the mean of the observed data. $R^2$ is calculated as:

$$R^2 = 1 - \frac{\sum_{i=1}^{N}(y_i - \hat{y}_i)^2}{\sum_{i=1}^{N}(y_i - \bar{y})^2} \tag{79}$$

where $N$ is the dataset size, $\hat{y}_i$ and $y_i$ signify the predicted and experimental values for each sample $i$, respectively, and $\bar{y}$ is the average of the observed value.

**Spearman's Rank Correlation Coefficient (ρ).** To evaluate the monotonic relationship between predicted and reference values, we calculated the Spearman rank correlation coefficient (ρ). ρ measures the strength and direction of the monotonic association between the ranked values of the predictions and reference data. By operating on ranks rather than raw values, this metric is robust to outliers and non-linear monotonic transformations, making no assumptions regarding the normality of the underlying distributions. The value ranges from -1.0 to 1.0, where

1.0 indicates a perfect positive monotonic relationship, and -1.0 represents a perfect negative monotonic relationship. The metric is defined as follows:

$$\rho = 1 - \frac{6\sum_{i=1}^{n} d_i^2}{n(n^2 - 1)} \tag{80}$$

where $n$ denotes the total number of evaluated samples, and $d_i$ represents the difference between the ranks of the predicted and reference values of the $i$-th sample.

**Cosine Similarity.** Cosine similarity measures the similarity between two non-zero vectors by calculating the cosine of the angle between them. Given two vectors $X = (X_1, \dots, X_p)$ and $Y = (Y_1, \dots, Y_p)$, cosine similarity is defined as follows:

$$\text{Cosine Similarity}(X, Y) = \frac{\sum_{i=1}^{p} X_i Y_i}{\sqrt{\sum_{i=1}^{p} X_i^2}\sqrt{\sum_{i=1}^{p} Y_i^2}} \tag{81}$$

where $p$ denotes the dimensionality of the vectors, and $X_i$ and $Y_i$ represent their $i$-th components. The metric ranges from $-1$ to $1$, where $1$ indicates that the two vectors point in the same direction, $0$ indicates that they are orthogonal, and $-1$ indicates that they point in opposite directions. Because the calculation normalizes the vectors by their magnitudes, cosine similarity primarily captures directional or pattern similarity rather than differences in absolute scale.

##### S6.3 Molecular docking

**Ligand root-mean-square deviation (L-RMSD).** To quantify the overall accuracy of the predicted ligand binding pose, we calculated the ligand root-mean-square deviation (L-RMSD) between the predicted ligand coordinates and the corresponding experimentally determined bound structure. In contrast to an RMSD calculated after structural superposition, L-RMSD was evaluated directly in the receptor coordinate frame and therefore jointly captures errors in binding-site localization, ligand orientation and internal conformation. Hydrogen atoms were excluded before evaluation. To account for molecular symmetries and differences in atom ordering, the minimum RMSD over all chemically valid symmetry-equivalent atom mappings was used. The corresponding formulation is provided below:

$$\text{L-RMSD} = \min_{\pi \in \Pi} \sqrt{\frac{1}{N}\sum_{i=1}^{N} \left\| \hat{x}_i - x_{\pi(i)} \right\|_2^2} \tag{82}$$

where $\hat{x}_i \in R^3$ denotes the predicted coordinate of ligand heavy atom $i$, $x_{\pi(i)}$ denotes the coordinate of its corresponding atom in the experimentally determined structure under symmetry-equivalent mapping $\pi$, $N$ is the number of ligand heavy atoms, and $\Pi$ is the set of chemically valid atom correspondences. Lower L-RMSD values indicate more accurate recovery of the complete bound pose.

**Centroid distance.** To separately assess whether a docking method identified the correct binding region, we calculated the Euclidean distance between the geometric centroids of the predicted and experimentally determined ligand poses. Unlike L-RMSD, centroid distance is insensitive to the detailed orientation and internal conformation of the ligand and primarily reflects translational displacement from the native binding pocket. The ligand centroid was calculated as the unweighted mean of all ligand heavy-atom coordinates:

$$\hat{c} = \frac{1}{N}\sum_{i=1}^{N} \hat{x}_i \tag{83}$$

$$c = \frac{1}{N}\sum_{i=1}^{N} x_i \tag{84}$$

and the centroid distance was defined as:

$$d_{\text{centroid}} = \left|\left|\hat{c} - c\right|\right|_2 \tag{85}$$

where $\hat{c}$ and $c$ denote the geometric centroids of the predicted and reference ligand poses, respectively, and $N$ is the number of ligand heavy atoms. A smaller centroid distance indicates more accurate localization of the ligand within the receptor, whereas a large value indicates that the ligand was placed in an incorrect or substantially displaced binding region.

**Kabsch-aligned root-mean-square deviation (Kabsch RMSD).** To quantify differences in ligand conformation independently of global translation and rotation, we calculated the minimum RMSD obtainable after optimal rigid-body superposition of the predicted and reference ligand structures using the Kabsch algorithm. This procedure identifies the rotation and translation that minimize the squared distances between corresponding ligand atoms while preserving the internal geometry of each structure. Kabsch-RMSD therefore primarily reflects discrepancies in ligand bond rotations and internal conformation rather than errors in binding-site placement or global orientation. The metric was calculated as:

$$\text{Kabsch-RMSD} = \min_{\pi\in\Pi} \min_{R\in SO(3), t\in R^3} \sqrt{\frac{1}{N}\sum_{i=1}^{N} \left|\left|R\hat{x}_i + t - x_{\pi(i)}\right|\right|_2^2} \quad (86)$$

subject to $R^T R = I$ and $det(R) = 1$, where $R$ in $SO(3)$ is the optimal rotation matrix, $t$ is the optimal translation vector, and the remaining variables are defined as above. Lower Kabsch-RMSD values indicate greater agreement between the internal conformations of the predicted and reference ligands.

##### S6.4 De novo molecular generation

**Fréchet ChemNet Distance (FCD)[75].** To evaluate the similarity between the generated and reference molecular distributions, we employed the Fréchet ChemNet Distance (FCD). FCD is calculated from activations of the penultimate layer of ChemNet, a neural network trained to predict biological activities from canonical SMILES representations. These activations encode both chemical and biological characteristics of the molecules. FCD compares the means and covariance matrices of the ChemNet activation distributions obtained from the generated and reference sets. Lower FCD values indicate closer agreement between the two molecular distributions and therefore better overall generation quality. The metric is defined as follows:

$$\text{FCD}(G, R) = |\mu_G - \mu_R|_2^2 + \text{Tr}\left(\Sigma_G + \Sigma_R - 2(\Sigma_G\Sigma_R)^{\frac{1}{2}}\right) \quad (87)$$

where $G$ and $R$ denote the generated and reference molecular sets, respectively; $\mu_G$ and $\mu_R$ are the corresponding mean ChemNet activation vectors; and $\Sigma_G$ and $\Sigma_R$ are their full covariance matrices.

**Scaffold Similarity (Scaff).** To evaluate the agreement between the scaffold distributions of the generated and reference molecular sets, we employed scaffold similarity (Scaff). Bemis-Murcko scaffolds[59] retain the ring systems of a molecule and the linker fragments connecting those rings, while removing peripheral side chains. Following MOSES, scaffolds were extracted using the RDKit implementation, which additionally treats carbonyl groups attached to rings as part of the scaffold. Scaff compares the occurrence frequencies of all scaffolds present in either the generated or reference set using cosine similarity. Higher Scaff values indicate that the generated set more closely reproduces the scaffold-frequency distribution of the reference set, whereas lower values indicate that particular chemotypes are underrepresented, overrepresented or absent. The metric is defined as follows:

$$\text{Scaff}(G,R) = \frac{\sum_{s\in S} c_s(G)c_s(R)}{\sqrt{\sum_{s\in S} c_s(G)^2}\sqrt{\sum_{s\in S} c_s(R)^2}} \tag{88}$$

where $S$ denotes the union of scaffolds occurring in the generated or reference set, and $c_s(A)$ denotes the number of occurrences of scaffold s among molecules in set $A$.

**Novelty.** To assess whether the model generates molecules beyond those observed during training, we employed the novelty metric. Novelty is defined as the fraction of valid generated molecules that are absent from the training set. It therefore provides a direct measure of the model's ability to produce previously unseen molecular structures. Low novelty indicates that the model frequently reproduces training molecules and may consequently reflect overfitting, whereas higher novelty indicates greater exploration beyond the training data.

**Internal Diversity (IntDiv$\boldsymbol{p}$)[76].** To assess structural diversity within the generated molecular set, we employed internal diversity (IntDiv). This metric summarizes pairwise Tanimoto similarities between generated molecules and converts them into a diversity score, thereby detecting mode collapse, in which a generative model produces only a restricted range of molecular structures while neglecting other regions of chemical space. The parameter $p$ determines the order used to aggregate pairwise similarities. Following MOSES, we report both IntDiv1 and IntDiv2, corresponding to $p = 1$ and $p = 2$, respectively. Higher values indicate greater chemical diversity within the generated set. The metric is defined as follows:

$$\text{IntDiv}_p(G) = 1 - \left(\frac{1}{|G|^2}\sum_{m_1,m_2\in G} T(m_1,m_2)^p\right)^{\frac{1}{p}} \tag{89}$$

where $G$ denotes the valid generated molecular set, $|G|$ is the number of molecules in $G$, and $T(m_1, m_2)$ represents the Tanimoto similarity between the molecular fingerprints of molecules $m_1$ and $m_2$.

**Uniqueness (Unique@K).** To evaluate whether the model repeatedly generates the same molecular structures, we employed the uniqueness metric. Uniqueness measures the fraction of distinct molecules among the first $K$ valid generated molecules. Following MOSES, we report Unique@k and Unique@10k by setting $K = 1000$ and $K = 10000$, respectively. If fewer than $K$ valid molecules are available, the metric is calculated over all valid generated molecules. Higher values indicate lower duplication and a reduced tendency toward mode collapse.

##### S6.5 Target-specific peptide binder design

**Pseudo-perplexity (PPL).** Following PepMLM[5], we adopted pseudo-perplexity (PPL) as a sequence-level metric to evaluate the quality and plausibility of the generated peptide binders. Unlike conventional perplexity, which is computed over the entire sequence, the PPL calculation was restricted to the generated binder region. Specifically, each residue in the binder sequence was masked individually, and the probability of correctly recovering the masked residue was estimated conditioned on all remaining binder residues and the target protein context. The pseudo-perplexity was then calculated as the exponential of the average negative log-likelihood across all positions in the binder sequence. Lower PPL values indicate that the generated peptide is more consistent with the distribution learned by the underlying protein language model and therefore exhibits higher sequence plausibility. The corresponding formulation is provided below:

$$\mathrm{PPL}(b) = \exp\left\{-\frac{1}{m}\sum_{i=1}^{m}\log P\left(b_i \mid b_{j\neq i}, p\right)\right\} \tag{90}$$

where b denotes the binder sequence, m represents the binder length, and $\log P\left(b_i \middle| b_{j\neq i}, p\right)$ is the conditional probability assigned to residue $b_i$ given the remaining binder residues and the target protein sequence $p$.

**Similarity.** To evaluate the preservation of sequence patterns and functional motifs in generated peptides, we measured sequence similarity between the designed and native peptide sequences. Rather than relying solely on exact residue correspondence, this metric captures broader biochemical conservation by accounting for substitutions with related physicochemical properties. Sequence similarity was computed using the Smith-Waterman local alignment algorithm[77], which identifies the highest-scoring alignment between two sequences and remains robust to terminal extensions, truncations, and local positional shifts. The resulting alignment score was normalized to facilitate comparison across peptide targets of different lengths. Higher similarity values indicate greater conservation of sequence features relative to the native peptide. The corresponding formulation is provided below:

$$\mathrm{Similarity} = \frac{\mathrm{Score}_{\mathrm{opt}}\left(s_{\mathrm{gen}}, s_{\mathrm{ref}}\right)}{\max\left(L_{\mathrm{gen}}, L_{\mathrm{ref}}\right)} \tag{91}$$

where $\mathrm{Score}_{\mathrm{opt}}$ represents the optimal alignment score derived from a pairwise aligner, and $L_{\mathrm{gen}}$ and $L_{\mathrm{ref}}$ denote the lengths of the generated and reference sequences, respectively. This metric accounts for the relative positioning of residues and penalizes gaps and mismatches, providing a rigorous measure of the model's ability to reproduce the essential sequence features and biochemical signatures of the target peptides.

**Sequence Diversity.** For each target, pairwise similarities among the generated peptide sequences were calculated using global sequence alignment with the BLOSUM62 substitution matrix. The alignment score between two sequences was normalized by the geometric mean of their respective self-alignment scores, and the corresponding pairwise distance was defined as one minus the normalized similarity. The resulting distance matrix was subjected to single-linkage hierarchical clustering with a distance threshold of 0.4. Sequence diversity was calculated as the number of resulting clusters divided by the total number of generated sequences, and the final value was averaged across all targets.

**The Interface Predicted TM-score (ipTM).** Following structure prediction with AlphaFold3[78], the interface predicted TM-score (ipTM) was used to assess the confidence and structural quality of the predicted peptide-receptor complexes. Unlike global structure confidence metrics, ipTM specifically evaluates the accuracy of inter-chain interactions by measuring the predicted relative positioning and packing of the peptide ligand with respect to the receptor. The score is derived from the Predicted Aligned Error (PAE) matrix and aggregates confidence estimates across residue pairs located at the protein-peptide interface. Higher ipTM values indicate greater confidence in the predicted binding mode and a more reliable interface geometry, whereas lower values suggest increased uncertainty in the relative arrangement of the interacting chains.

### Supplementary Discussions

**Discussion S1**. Mechanistic insights into evolutionary search in DrugEvolve.

Using protein druggable-site annotation as a representative task, we compared four foundation models, including Gemini 3.5 Flash, Claude Sonnet 4.6, GPT-5.5, and Seed 2.0 Pro, as unified system-level controllers of DrugEvolve. To enable a rapid and standardized comparison, the same model was assigned to all functional agents within each configuration. The performance distributions across ten evolutionary runs for each foundation model are shown in **Supplementary Figure S13a**, where results are reported conditional on 10 runs for clarity of visualization. We observe that the GPT-5.5 configuration achieves the highest peak performance, indicating a stronger observed performance ceiling in evolutionary trajectories. Claude Sonnet 4.6 exhibits a slightly higher average performance within the above-baseline-score subset but produces a larger number of low-performance outcomes. In contrast, Gemini 3.5 Flash and Seed 2.0 Pro show fewer below-baseline-score cases but also exhibit a more limited high-performance tail, suggesting more conservative but less exploratory behavior. Overall, these results suggest that the choice of foundation model may influence the stability and performance distribution of evolutionary search. Importantly, these observations are specific to the evaluated task and evolutionary configuration in DrugEvolve, and should be interpreted as empirical behavior under this experimental setting. In practice, different foundation models may be flexibly adopted depending on desired exploration characteristics and application requirements.

Sampling is a critical component of evolutionary search, particularly in the evolution of site-prediction models, where the search process must accommodate heterogeneous task categories and multiple evaluation metrics[79]. To assess how sampling strategies influence evolutionary outcomes, we compared two sampling frameworks: UCB1-based sampling and island-based sampling. Within the UCB framework, we evaluated two variants, one using only an aggregated score and the other jointly using the aggregated score and task-specific auxiliary scores. These variants were compared with island-based sampling[80]. For each strategy, we selected the top-10 evolutionary cases and conducted a comparative trajectory analysis (**Supplementary Figure S13b**). Among the selected top-performing trajectories, island-based sampling exhibited the strongest performance in the evolution of unified site-prediction models. The approach preserved a broader set of candidate lineages throughout evolution while simultaneously achieving higher final scores, indicating improved maintenance of population diversity. We speculate that such diversity preservation is particularly beneficial in heterogeneous optimization settings involving multiple task categories and evaluation objectives, where early convergence to locally favorable solutions may impede discovery of more generalizable algorithmic designs.

We further observed that prompt design may influence both the performance stability and exploratory breadth of DrugEvolve. To examine this effect, we compared two prompting schemes: a framework-constrained setting, in which DrugEvolve was instructed to modify architectures within the baseline framework, and a framework-free setting, in which no explicit architectural scaffold was imposed (**Supplementary Method S3**). We analyzed the mean and variance of the resulting scores, together with the spatial distribution of semantic embeddings of the generated design proposals after t-SNE dimensionality reduction (**Supplementary Figure S13c, d**). The framework-constrained prompts produced evolved models with lower score variance, consistent with more stable performance, and their semantic embeddings appeared more tightly clustered in the two-dimensional t-SNE projection. In contrast, framework-free prompts yielded higher score variance but slightly higher mean performance, with a visually broader distribution of embeddings. Although the t-SNE projection does not provide a quantitative measure of architectural diversity, this pattern is consistent with framework-free prompting exploring a broader range of design concepts under the evaluated setting.

Finally, to assess whether DrugEvolve maintains consistency between algorithmic conception and executable implementation, we compared the design motivations proposed by the generator with the corresponding code produced by the implementer across a subset of representative cases (**Supplementary Table S19**). In most cases, the implemented models faithfully instantiate the intended algorithmic designs, including core architectural modifications and their expected functional roles. A small number of cases exhibit partial simplification during implementation, likely reflecting practical considerations such as computational tractability, training stability, or compatibility with the existing modeling pipeline. Importantly, these simplifications do not substantially deviate from the original design rationale. Overall, this analysis indicates a high degree of alignment between the proposed algorithmic ideas and their realized implementations, supporting the effectiveness of the proposed design-implementation pipeline.

**Table S1**. Comprehensive summary of the 11 tasks, baseline, and computational costs evaluated by DrugEvolve across four drug development stages.

| Stage | Task | Evaluation scale | Benchmark settings | Baseline | Baseline Reference | Rounds | Total single-GPU runtime (h) |
|---|---|---|---|---|---|---|---|
| Target identification | Cancer gene module detection | MCF-7 breast cancer multi-omics dataset with 10-fold evaluation | 1 | CGMega | *Nat Commun*. 15, 5997 (2024) | 151 | 361 |
| | Gene-disease association prediction | DisGeNET-EVAL with five random splits and TDC as an independent test set | 2 | FusionGDA | *Brief Bioinform*. 25, bbae380 (2024) | 3 | 1 |
| | Protein druggable-site annotation | 79 tests spanning noncovalent, covalent, catalytic, cryptic, and PTM sites | 79 | ALLSites | *Adv Sci*. 13, e16530 (2026) | 392 | 2254 |
| Drug discovery | Molecular docking | PDBbind, PoseBusters, and Astex dataset | 3 | EquiBind | *International conference on machine learning* 20503-20521 (PMLR, 2022) | 24 | 864 |
| | Drug-target interaction prediction | BindingDB under random, unseen-drug, and unseen-protein settings, together with the Human-cold benchmark | 4 | DrugBAN | *Nat Mach Intell*. 5, 126-136 (2023) | 21 | 21 |
| | De novo molecular generation | MOSES dataset with test and testSF splits | 2 | CharRNN | *ACS Cent Sci*. 4, 120-131 (2018) | 2 | 3 |
| | Target-specific peptide binder design | PepBench and PepNN dataset | 2 | PepMLM | *Nat Biotechnol*. 44, 1002-1010 (2026) | 3 | 5 |
| | ADMET property prediction | 12 classification and 5 regression endpoints | 17 | FragNet | *J Am Chem Soc*. 148, 9930-9950 (2026) | 156 | 312 |
| Preclinical study | Animal toxicology assessment | One dataset evaluated under four compound-level splitting strategies | 4 | AnimalMLP | Developed in this study | 10 | 4 |
| Clinical trial | Drug-drug interaction prediction | 2 classification settings and 1 regression benchmark | 3 | MeTDDI | *Nat Mach Intell*. 6, 1094-1105 (2024) | 139 | 140 |
| | Drug Side-effect prediction | SIDER datasets under 10-fold, unseen-pair, and unseen-drug settings | 3 | SDPred | *Brief Bioinform*. 23, bbab449 (2022) | 150 | 48 |

**Table S2**. Quantitative performance comparison of CGMega and CGMega-Evo on the MCF-7 breast cancer multi-omics dataset.

| **Method** | AUROC | AUPRC | ACC | F1 | MCC | Precision | Recall | Specificity |
|---|---|---|---|---|---|---|---|---|
| CGMega | 0.961 | 0.910 | 0.924 | 0.812 | 0.769 | 0.748 | **0.889** | 0.932 |
| CGMega-Evo | **0.964** | **0.915** | **0.942** | **0.844** | **0.809** | **0.844** | 0.844 | **0.965** |

**Table S3.** Performance evaluation of ALLSites-Evo on the PPI-Test70 dataset. The best performance for each metric is highlighted in bold and the second-best performance is underlined.

| Class | Method | ACC | AUROC | AUPRC | F1 | MCC |
|---|---|---|---|---|---|---|
| Structure-based | IntPred [a] | 0.672 | - | - | 0.332 | 0.165 |
| | SPPIDER [c] | 0.667 | 0.518 | 0.235 | 0.273 | 0.063 |
| | DeepPPISP [b] | 0.655 | 0.671 | 0.320 | 0.397 | 0.206 |
| | EGRET [b] | 0.715 | <u>0.719</u> | <u>0.405</u> | 0.438 | 0.270 |
| Sequence-based | ISIS [a] | 0.622 | - | 0.240 | 0.267 | 0.097 |
| | RF_PPI [a] | 0.598 | - | 0.210 | 0.258 | 0.118 |
| | PSIVER [a] | 0.653 | - | 0.250 | 0.328 | 0.138 |
| | SPRINGS [b] | 0.631 | - | 0.280 | 0.350 | 0.181 |
| | ProNA2020 [c] | **0.741** | - | - | 0.258 | 0.106 |
| | SCRIBER [c] | 0.616 | 0.635 | 0.307 | 0.370 | 0.159 |
| | DELPHI [b] | 0.667 | 0.690 | 0.360 | 0.418 | 0.236 |
| | DLPred [c] | 0.680 | 0.697 | 0.380 | 0.416 | 0.235 |
| | EnsemPPIS [b] | <u>0.732</u> | 0.719 | 0.405 | 0.440 | 0.277 |
| | ALLSites-Evo | 0.705 | **0.766** | **0.449** | **0.475** | **0.321** |

*Note*: [a] Results reported by DeepPPISP. [b] Results generated by reproducing the source code. [c] Results obtained by using the web server. ProNA2020 only makes binary predictions, and its AUROC and AUPRC are not calculated.

**Table S4.** Performance evaluation of ALLSites-Evo on the PPI-Test355 dataset. All the comparison methods use only protein sequences. The best performance for each metric is highlighted in bold and the second-best performance is underlined.

| Method | ACC | AUROC | AUPRC | F1 | MCC |
|---|---|---|---|---|---|
| SPRINGS [a] | 0.811 | 0.608 | 0.178 | 0.211 | 0.103 |
| DLPred [b] | 0.835 | 0.724 | 0.272 | 0.308 | 0.214 |
| SCRIBER [b] | 0.838 | 0.719 | 0.275 | 0.322 | 0.230 |
| DELPHI [a] | 0.848 | 0.746 | 0.326 | 0.364 | 0.278 |
| EnsemPPIS [a] | 0.821 | 0.770 | 0.354 | 0.385 | 0.291 |
| ALLSites-Evo | **0.858** | **0.828** | **0.455** | **0.470** | **0.392** |

*Note*: [a] Results generated by reproducing the source code. [b] Results obtained by using the web server.

**Table S5**. Detailed docking performance comparison of EquiBind and EquiBind-Evo variants on PDBbind, PoseBusters, and Astex Diverse Set benchmarks. The best result for each dataset is depicted in **bold**.

| Dataset | Model | Ligand RMSD ↓ | | | | | | Centroid Distance ↓ | | | | | | Kabsch RMSD ↓ | |
|---|---|---|---|---|---|---|---|---|---|---|---|---|---|---|---|
| | | | Percentiles ↓ | | | %Below threshold ↑ | | | Percentiles↓ | | | %Below threshold ↑ | | | |
| | | Mean | 25TH | 50TH | 75TH | 5Å | 2Å | Mean | 25TH | 50TH | 75TH | 5Å | 2Å | Mean | Med |
| PDBbind | EquiBind-U | 8.075 | 3.778 | 6.233 | 10.588 | 39.669 | 3.857 | 5.92 | 1.632 | 3.101 | 7.866 | 61.157 | 32.231 | 2.56 | 2.177 |
| | EquiBind | 8.381 | 4.191 | 6.447 | 11.193 | 36.639 | 4.132 | 5.92 | 1.632 | 3.101 | 7.866 | 61.157 | 32.231 | 2.501 | 2.302 |
| | EquiBind-Evo-U | **7.946** | **3.232** | **5.68** | **10.444** | **44.628** | **10.468** | **5.871** | **1.283** | **2.817** | **7.569** | **66.667** | **41.322** | 2.488 | **2.114** |
| | EquiBind-Evo | 8.231 | 3.591 | 5.889 | 11.181 | 41.322 | 7.713 | 5.871 | 1.283 | 2.817 | 7.569 | 66.667 | 41.322 | **2.467** | 2.311 |
| Posebusters | EquiBind-U | 10.892 | 6.71 | 9.521 | 13.816 | 10.514 | 0 | 9.41 | 4.964 | 7.975 | 12.484 | 25.701 | 3.972 | 1.999 | 1.87 |
| | EquiBind | 11.123 | 7.073 | 9.843 | 13.886 | 8.411 | 0 | 9.41 | 4.964 | 7.975 | 12.484 | 25.701 | 3.972 | 1.936 | 1.919 |
| | EquiBind-Evo-U | **10.231** | **5.692** | **8.221** | **13.17** | **19.626** | 0.234 | **8.548** | **3.391** | **5.918** | **12.378** | **41.822** | **7.009** | 1.854 | **1.755** |
| | EquiBind-Evo | 10.467 | 6.022 | 8.592 | 13.285 | 15.654 | **0.467** | 8.548 | 3.391 | 5.918 | 12.378 | 41.822 | 7.009 | **1.842** | 1.77 |
| Astex | EquiBind-U | 8.847 | **4.45** | 7.044 | 10.483 | 29.412 | 0 | 7.421 | 3.082 | 4.991 | 9.002 | 50.588 | 10.588 | 1.685 | 1.631 |
| | EquiBind | 9.086 | 4.661 | 7.397 | 10.789 | 28.235 | 0 | 7.421 | 3.082 | 4.991 | 9.002 | 50.588 | 10.588 | 1.658 | 1.548 |
| | EquiBind-Evo-U | **8.595** | 4.527 | **6.639** | **9.873** | **30.588** | 0 | **7.003** | **2.548** | **4.44** | **8.038** | **55.294** | **16.471** | 1.641 | 1.589 |
| | EquiBind-Evo | 8.827 | 4.894 | 6.958 | 10.067 | 27.059 | 0 | 7.003 | 2.548 | 4.44 | 8.038 | 55.294 | 16.471 | **1.615** | **1.534** |

**Table S6.** Performance comparison of MeTDDI and MeTDDI-Evo on unseen-one-drug and unseen-two-drug scenarios.

| **Model** | Unseen one drug | | | Unseen two drugs | | |
|---|---|---|---|---|---|---|
| | ACC | AUROC | AUPRC | ACC | AUROC | AUPRC |
| MeTDDI | 0.677 | 0.883 | 0.740 | 0.522 | 0.778 | 0.539 |
| MeTDDI-Evo | 0.685 | 0.885 | 0.734 | 0.538 | 0.786 | 0.542 |

**Table S7**. Fitness score design for DrugEvolve algorithm evolution on protein druggable site annotation, including metric selection, weighting schemes, and normalization constants.

| Benchmark | Metric | Weight | Normalization constant |
|---|---|---|---|
| CarbPI | Recall | 0.200 | 0.100 |
| | Precision | 0.200 | 0.100 |
| | F1-score | 0.250 | 0.050 |
| | MCC | 0.350 | 0.040 |
| DPI | AUROC | 0.175 | 0.025 |
| | AUPRC | 0.175 | 0.030 |
| | Recall | 0.075 | 0.100 |
| | Precision | 0.075 | 0.100 |
| | F1-score | 0.200 | 0.050 |
| | MCC | 0.300 | 0.040 |
| PepPI | AUROC | 0.400 | 0.025 |
| | MCC | 0.600 | 0.040 |
| PPI | AUROC | 0.175 | 0.025 |
| | AUPRC | 0.175 | 0.030 |
| | Accuracy | 0.100 | 0.100 |
| | F1-score | 0.250 | 0.050 |
| | MCC | 0.300 | 0.040 |
| RPI | AUROC | 0.200 | 0.025 |
| | Recall | 0.125 | 0.100 |
| | Precision | 0.125 | 0.100 |
| | F1-score | 0.250 | 0.050 |
| | MCC | 0.300 | 0.040 |
| SMPI | Accuracy | 0.150 | 0.100 |
| | Recall | 0.150 | 0.100 |
| | Precision | 0.150 | 0.100 |
| | F1-score | 0.250 | 0.050 |
| | MCC | 0.300 | 0.040 |

**Table S8.** Dataset statistics of the MCF-7 breast cancer multi-omics dataset for cancer gene module detection.

| Dataset | Split | No. genes | No. of cancer-associated genes | No. of non-cancer genes | Ratio of cancer-associated genes (%) |
|---|---|---|---|---|---|
| MCF-7 breast cancer multi-omics dataset | Train+Valid | 1454 | 268 | 1186 | 18.43% |
| | Test | 485 | 90 | 395 | 18.56% |

**Table S9.** Dataset statistics for universal protein binding site prediction across different interaction types.

| Site type | Task | Dataset | No. proteins | No. residues | No. binding residues | No. non-binding residues | Ratio of binding residues (%) |
|---|---|---|---|---|---|---|---|
| PPI | PPI-Train352 | PPI-Train282 | 282 | 59617 | 8972 | 50645 | 15.05 |
| | | PPI-Valid70 | 70 | 16632 | 2232 | 14400 | 13.42 |
| | | PPI-Test70 | 70 | 11791 | 2332 | 9459 | 19.78 |
| | PPI-Train9982 | PPI-Train7986 | 7986 | 3391855 | 342781 | 3049074 | 10.11 |
| | | PPI-Valid1996 | 1996 | 862343 | 84906 | 777437 | 9.85 |
| | | PPI-Test355 | 355 | 95940 | 11467 | 84473 | 11.95 |
| | PPI-335 | PPI-Train268 | 268 | 54105 | 8276 | 45829 | 15.30 |
| | | PPI-Valid67 | 67 | 12261 | 2098 | 10163 | 17.11 |
| | | PPI-Test60 | 60 | 13144 | 2075 | 11069 | 15.79 |
| | | PPI-Test315 | 315 | 65331 | 9355 | 55976 | 14.32 |
| PepPI | PepPI-1154 | PepPI-Train924 | 924 | 222440 | 11940 | 210500 | 5.37 |
| | | PepPI-Valid230 | 230 | 54382 | 3093 | 51289 | 5.69 |
| | | PepPI-Test125 | 125 | 30870 | 1716 | 29154 | 5.56 |
| | PepPI-Train640 | PepPI-Train512 | 512 | 128181 | 6618 | 121563 | 5.16 |
| | | PepPI-Valid128 | 128 | 29181 | 1641 | 27540 | 5.62 |
| | | PepPI-Test639 | 639 | 150330 | 8490 | 141840 | 5.65 |
| SMPI | SMPI-1628 | SMPI-Train1628 | 1628 | 678373 | 61210 | 617163 | 9.02 |
| | | SMPI-Valid348 | 348 | 138926 | 13004 | 125922 | 9.36 |
| | | SMPI-Test348 | 348 | 140821 | 13171 | 127650 | 9.35 |
| CarbPI | Carb-517 | Carb-Train517 | 517 | 186665 | 7207 | 179458 | 3.86 |
| | | Carb-Valid129 | 129 | 49680 | 1815 | 47865 | 3.65 |
| | | Carb-Test162 | 162 | 56928 | 2237 | 54691 | 3.93 |
| DPI | DPI-573 | DPI-Train459 | 459 | 129074 | 11934 | 117140 | 9.25 |
| | | DPI-Valid114 | 114 | 30809 | 2545 | 28264 | 8.26 |
| | | DPI-Test129 | 129 | 37515 | 2240 | 35275 | 5.97 |
| | | DPI-Test181 | 181 | 75258 | 3208 | 72050 | 4.26 |
| RPI site | RPI-Train495 | RPI-Train396 | 396 | 112903 | 11614 | 101289 | 10.29 |
| | | RPI-Valid99 | 99 | 23996 | 2995 | 21001 | 12.48 |
| | | RPI-Test117 | 117 | 37345 | 2031 | 35314 | 5.44 |

**Table S10.** Dataset statistics for ion-binding site prediction across different ion types.

| Site type | Task | Target Ion | Dataset | No. proteins | No. residues | No. binding residues | No. non-binding residues | Ratio of binding residues (%) |
|---|---|---|---|---|---|---|---|---|
| Ion-binding | LigBind | $Ca^{2+}$ | Train | 1154 | 376086 | 6215 | 369871 | 1.65 |
| | | | Valid | 267 | 85066 | 1473 | 83593 | 1.73 |
| | | | Test | 533 | 214683 | 3365 | 211318 | 1.57 |
| | | $Mg^{2+}$ | Train | 1360 | 457415 | 4779 | 452636 | 1.04 |
| | | | Valid | 341 | 111328 | 1250 | 110078 | 1.12 |
| | | | Test | 665 | 291215 | 2490 | 288725 | 0.86 |
| | | $Mn^{2+}$ | Train | 405 | 137546 | 1773 | 135773 | 1.29 |
| | | | Valid | 108 | 38154 | 568 | 37586 | 1.49 |
| | | | Test | 167 | 64330 | 778 | 63552 | 1.21 |
| | | $Zn^{2+}$ | Train | 1420 | 375847 | 6743 | 369104 | 1.79 |
| | | | Valid | 350 | 92247 | 1646 | 90601 | 1.78 |
| | | | Test | 615 | 215861 | 3150 | 212711 | 1.46 |
| | | $Fe^{2+}$ | Train | 86 | 28415 | 351 | 28064 | 1.24 |
| | | | Valid | 22 | 6968 | 102 | 6866 | 1.46 |
| | | | Test | 31 | 9788 | 123 | 9665 | 1.26 |
| | | $Fe^{3+}$ | Train | 181 | 56507 | 878 | 55629 | 1.55 |
| | | | Valid | 56 | 17267 | 246 | 17021 | 1.42 |
| | | | Test | 88 | 30663 | 345 | 30318 | 1.13 |
| | | $Cu^{2+}$ | Train | 107 | 32219 | 435 | 31784 | 1.35 |
| | | | Valid | 22 | 6461 | 84 | 6377 | 1.3 |
| | | | Test | 54 | 16589 | 241 | 16348 | 1.45 |
| | | $Na^{+}$ | Train | 79 | 25408 | 456 | 24952 | 1.79 |
| | | | Valid | 24 | 8116 | 164 | 7952 | 2.02 |
| | | | Test | 23 | 7617 | 199 | 7418 | 2.61 |
| | | $K^{+}$ | Train | 52 | 18938 | 501 | 18437 | 2.65 |
| | | | Valid | 12 | 4209 | 138 | 4071 | 3.28 |
| | | | Test | 17 | 7441 | 147 | 7294 | 1.98 |
| | | $CO_3^{2-}$ | Train | 57 | 18421 | 260 | 18161 | 1.41 |
| | | | Valid | 12 | 5237 | 70 | 5167 | 1.34 |
| | | | Test | 21 | 8684 | 99 | 8585 | 1.14 |
| | | $SO_4^{2-}$ | Train | 288 | 85491 | 1978 | 83513 | 2.31 |
| | | | Valid | 82 | 25100 | 563 | 24537 | 2.24 |
| | | | Test | 31 | 10812 | 230 | 10582 | 2.13 |
| | | $NO_2^{-}$ | Train | 14 | 4695 | 62 | 4633 | 1.32 |
| | | | Valid | 5 | 2441 | 21 | 2420 | 0.86 |
| | | | Test | 7 | 1633 | 43 | 1590 | 2.63 |
| | | $PO_4^{3-}$ | Train | 322 | 100967 | 1997 | 98970 | 1.98 |
| | | | Valid | 71 | 23889 | 450 | 23439 | 1.88 |

| | | | | | | | |
|---|---|---|---|---|---|---|---|
| | | Test | 73 | 26724 | 482 | 26242 | 1.8 |
| | ADP | Train | 316 | 132629 | 4259 | 128370 | 3.21 |
| | | Valid | 83 | 32296 | 1179 | 31117 | 3.65 |
| | | Test | 158 | 80859 | 1903 | 78956 | 2.35 |
| | AMP | Train | 183 | 60894 | 1993 | 58901 | 3.27 |
| | | Valid | 52 | 17894 | 595 | 17299 | 3.33 |
| | | Test | 67 | 26673 | 786 | 25887 | 2.95 |
| | ATP | Train | 265 | 106350 | 3534 | 102816 | 3.32 |
| | | Valid | 71 | 28914 | 990 | 27924 | 3.42 |
| | | Test | 139 | 73985 | 1943 | 72042 | 2.63 |
| | GDP | Train | 111 | 41629 | 1364 | 40265 | 3.28 |
| | | Valid | 16 | 5916 | 194 | 5722 | 3.28 |
| | | Test | 40 | 16198 | 496 | 15702 | 3.06 |
| | GTP | Train | 66 | 28683 | 866 | 27817 | 3.02 |
| | | Valid | 17 | 6619 | 186 | 6433 | 2.81 |
| | | Test | 34 | 15119 | 454 | 14665 | 3 |
| | HEM | Train | 158 | 43495 | 3513 | 39982 | 8.08 |
| | | Valid | 45 | 13100 | 1220 | 11880 | 9.31 |
| | | Test | 78 | 24541 | 1659 | 22882 | 6.76 |
| GPSites | $Ca2^{+}$ | Train | 1240 | 404082 | 6670 | 397412 | 1.65 |
| | | Valid | 314 | 100064 | 1772 | 98292 | 1.77 |
| | | Test | 183 | 66854 | 1034 | 65820 | 1.55 |
| | $Mg^{2+}$ | Train | 1379 | 459659 | 5091 | 454568 | 1.11 |
| | | Valid | 350 | 116073 | 1226 | 114847 | 1.06 |
| | | Test | 235 | 88806 | 893 | 87913 | 1.01 |
| | $Mn^{2+}$ | Train | 443 | 148024 | 2096 | 145928 | 1.42 |
| | | Valid | 104 | 33675 | 460 | 33215 | 1.37 |
| | | Test | 57 | 20419 | 225 | 20194 | 1.1 |
| | $Zn^{2+}$ | Train | 1311 | 381266 | 6037 | 375229 | 1.58 |
| | | Valid | 335 | 93589 | 1690 | 91899 | 1.81 |
| | | Test | 211 | 56020 | 1039 | 54981 | 1.85 |
| | ATP | Train | 285 | 107729 | 4208 | 103521 | 3.91 |
| | | Valid | 62 | 22926 | 907 | 22019 | 3.96 |
| | | Test | 79 | 39459 | 1231 | 38228 | 3.12 |
| | HEM | Train | 141 | 36779 | 3283 | 33496 | 8.93 |
| | | Valid | 35 | 10284 | 739 | 9545 | 7.19 |
| | | Test | 48 | 15618 | 970 | 14648 | 6.21 |

**Table S11.** Dataset statistics for covalent binding site prediction across different covalent binding types.

| Site type | Task | Target Residue | Dataset | No. proteins | No. target residues | No. binding residues | No. non-binding residues | Ratio of binding residues (%) |
|---|---|---|---|---|---|---|---|---|
| Covalent | Covalent | DKHSTCREY | Train | 1545 | 373063 | 1535 | 371528 | 0.41 |
| | | | Valid | 193 | 46115 | 192 | 45923 | 0.42 |
| | | | Test | 194 | 45547 | 193 | 45354 | 0.42 |
| | Cys | C | Train | 188 | 1048 | 191 | 857 | 18.23 |
| | | | Valid | 23 | 134 | 23 | 111 | 17.16 |
| | | | Test | 23 | 120 | 24 | 96 | 20.00 |

**Table S12.** Dataset statistics for catalytic site prediction.

| Site type | Task | Dataset | No. proteins | No. residues | No. binding residues | No. non-binding residues | Ratio of binding residues (%) |
|---|---|---|---|---|---|---|---|
| Catalytic | CataloDB | Train | 4531 | 1860369 | 7452 | 1852917 | 0.4 |
| | | Valid | 504 | 213874 | 813 | 213061 | 0.38 |
| | | Test | 231 | 97113 | 413 | 96700 | 0.43 |
| | Uni14230 | Train | 7896 | 3272883 | 12804 | 3260079 | 0.39 |
| | | Valid | 878 | 359669 | 1407 | 358262 | 0.39 |
| | | Test | 1955 | 748322 | 3175 | 745147 | 0.42 |
| | Benchmark | EF_fold | 93 | 36575 | 165 | 36410 | 0.45 |
| | | EF_family | 196 | 84008 | 369 | 83639 | 0.44 |
| | | EF_superfamily | 123 | 47765 | 219 | 47546 | 0.46 |
| | | PC | 55 | 21238 | 99 | 21139 | 0.47 |
| | | HA_superfamily | 152 | 62783 | 275 | 62508 | 0.44 |
| | | NN | 108 | 45392 | 194 | 45198 | 0.43 |

**Table S13.** Dataset statistics for cryptic site prediction.

| Site type | Task | Dataset | No. proteins | No. residues | No. binding residues | No. non-binding residues | Ratio of binding residues (%) |
|---|---|---|---|---|---|---|---|
| Cryptic | Cryptobank | Train | 6345 | 2262014 | 123520 | 2138494 | 5.46 |
| | | Valid | 793 | 291612 | 15067 | 276545 | 5.17 |
| | | Test | 793 | 229424 | 9429 | 219995 | 4.11 |

**Table S14.** Dataset statistics for post-translational modification site prediction across different modification types.

| Site type | Task | Target Residue | Dataset | No. proteins | No. target residues | No. binding residues | No. non-binding residues | Ratio of binding residues (%) |
|---|---|---|---|---|---|---|---|---|
| PTM | Acetylation | K | Train | 4868 | 219648 | 23892 | 195756 | 10.88 |
| | | | Valid | 1216 | 52580 | 5833 | 46747 | 11.09 |
| | | | Test | 2286 | 121866 | 3239 | 118627 | 2.66 |
| | | | Test90 | 662 | 28505 | 2823 | 25682 | 9.9 |
| | | | Test80 | 648 | 28115 | 2689 | 25426 | 9.56 |
| | | | Test70 | 622 | 27290 | 2544 | 24746 | 9.32 |
| | N-glycosylation | N | Train | 2119 | 64078 | 5405 | 58673 | 8.44 |
| | | | Valid | 530 | 16930 | 1473 | 15457 | 8.7 |
| | | | Test | 677 | 28620 | 771 | 27849 | 2.69 |
| | | | Test90 | 288 | 8359 | 853 | 7506 | 10.2 |
| | | | Test80 | 284 | 8224 | 842 | 7382 | 10.24 |
| | | | Test70 | 276 | 8040 | 821 | 7219 | 10.21 |
| | Methylation | K | Train | 2390 | 141995 | 4393 | 137602 | 3.09 |
| | | | Valid | 595 | 39910 | 1113 | 38797 | 2.79 |
| | | | Test | 1308 | 78915 | 590 | 78325 | 0.75 |
| | | | Test90 | 329 | 19844 | 585 | 19259 | 2.95 |
| | | | Test80 | 327 | 19813 | 576 | 19237 | 2.91 |
| | | | Test70 | 324 | 19649 | 568 | 19081 | 2.89 |
| | | R | Train | 2752 | 129468 | 6865 | 122603 | 5.30 |
| | | | Valid | 689 | 33339 | 1655 | 31684 | 4.96 |
| | | | Test | 828 | 48524 | 947 | 47577 | 1.95 |
| | | | Test90 | 375 | 17241 | 954 | 16287 | 5.53 |
| | | | Test80 | 371 | 17082 | 917 | 16165 | 5.37 |
| | | | Test70 | 370 | 17027 | 891 | 16136 | 5.23 |
| | Phosphorylation | ST | Train | 9660 | 881762 | 128759 | 753003 | 14.6 |
| | | | Valid | 2417 | 219052 | 32902 | 186150 | 15.02 |
| | | | Test | 7479 | 831747 | 17656 | 814091 | 2.12 |
| | | | Test90 | 1286 | 115040 | 16992 | 98048 | 14.77 |
| | | | Test80 | 1275 | 114475 | 16733 | 97742 | 14.62 |
| | | | Test70 | 1261 | 113607 | 16170 | 97437 | 14.23 |
| | | Y | Train | 3506 | 70136 | 8231 | 61905 | 11.74 |
| | | | Valid | 874 | 17754 | 2014 | 15740 | 11.34 |
| | | | Test | 1740 | 47690 | 1081 | 46609 | 2.27 |
| | | | Test90 | 469 | 9344 | 1000 | 8344 | 10.7 |
| | | | Test80 | 462 | 9272 | 970 | 8302 | 10.46 |
| | | | Test70 | 454 | 9126 | 925 | 8201 | 10.14 |
| | Sumoylation | K | Train | 4568 | 202449 | 28142 | 174307 | 13.9 |

| | | | | | | | |
|---|---|---|---|---|---|---|---|
| | | Valid | 1142 | 49300 | 6681 | 42619 | 13.55 |
| | | Test | 2420 | 124857 | 3861 | 120996 | 3.09 |
| | | Test90 | 628 | 29000 | 3680 | 25320 | 12.69 |
| | | Test80 | 623 | 28890 | 3556 | 25334 | 12.31 |
| | | Test70 | 609 | 28321 | 3397 | 24924 | 11.99 |
| Ubiquitination | K | Train | 8377 | 331705 | 76422 | 255283 | 23.04 |
| | | Valid | 2098 | 81091 | 18687 | 62404 | 23.04 |
| | | Test | 6010 | 285448 | 10493 | 274955 | 3.68 |
| | | Test90 | 1146 | 44511 | 10160 | 34351 | 22.83 |
| | | Test80 | 1127 | 43945 | 9787 | 34158 | 22.27 |
| | | Test70 | 1105 | 43252 | 9259 | 33993 | 21.41 |

**Table S15.** Dataset statistics and split composition of drug-target interaction prediction benchmarks.

| Site type | Task | Dataset | No. proteins | No. drugs | No. interaction | No. non-interaction | Ratio of interaction (%) |
|---|---|---|---|---|---|---|---|
| BindingDB | Random | Train | 757 | 43155 | 28237 | 21912 | 56.31 |
| | | Valid | 471 | 5074 | 2830 | 2774 | 50.50 |
| | | Test | 465 | 5013 | 2705 | 2800 | 49.14 |
| | Unseen Drug | Train | 757 | 43155 | 28237 | 21912 | 56.31 |
| | | Valid | 471 | 5074 | 2830 | 2774 | 50.50 |
| | | Test | 361 | 3494 | 2142 | 1649 | 56.5 |
| | Unseen Protein | Train | 757 | 43155 | 28237 | 21912 | 56.31 |
| | | Valid | 471 | 5074 | 2830 | 2774 | 50.50 |
| | | Test | 42 | 2346 | 1249 | 1258 | 49.82 |
| Human | Cold-split | Train | 888 | 1062 | 240 | 1311 | 15.47 |
| | | Valid | 253 | 278 | 180 | 156 | 53.57 |
| | | Test | 475 | 506 | 340 | 332 | 50.60 |

**Table S16.** Dataset statistics for ADMET classification tasks.

| Type | Task | Dataset | No. drugs | No. interaction | No. non-interaction | Ratio of interaction (%) |
|---|---|---|---|---|---|---|
| A | F50% | Train | 1550 | 879 | 671 | 56.71 |
| | | Valid | 193 | 103 | 90 | 53.37 |
| | | Test | 195 | 102 | 93 | 52.31 |
| D | OCT2 inhibition | Train | 536 | 117 | 419 | 21.83 |
| | | Valid | 67 | 15 | 52 | 22.39 |
| | | Test | 68 | 18 | 50 | 26.47 |
| | OATP1B1 inhibition | Train | 1160 | 1030 | 130 | 88.79 |
| | | Valid | 145 | 127 | 18 | 87.59 |
| | | Test | 146 | 118 | 28 | 80.82 |
| M | HLM | Train | 4797 | 1679 | 3118 | 35 |
| | | Valid | 599 | 211 | 388 | 35.23 |
| | | Test | 601 | 236 | 365 | 39.27 |
| | CYP2B6 inhibition | Train | 390 | 153 | 237 | 39.23 |
| | | Valid | 48 | 7 | 41 | 14.58 |
| | | Test | 50 | 22 | 28 | 44 |
| | CYP2D6 inhibition | Train | 14428 | 2849 | 11579 | 19.75 |
| | | Valid | 1803 | 436 | 1367 | 24.18 |
| | | Test | 1804 | 487 | 1317 | 27 |
| | CYP2C9 substrate | Train | 1597 | 442 | 1155 | 27.68 |
| | | Valid | 199 | 44 | 155 | 22.11 |
| | | Test | 201 | 25 | 176 | 12.44 |
| T | Ames mutagenicity | Train | 6376 | 3611 | 2765 | 56.63 |
| | | Valid | 797 | 359 | 438 | 45.04 |
| | | Test | 797 | 402 | 395 | 50.44 |
| | hERG (1μM) | Train | 3594 | 612 | 2982 | 17.03 |
| | | Valid | 449 | 94 | 355 | 20.94 |
| | | Test | 450 | 91 | 359 | 20.22 |
| | FDAMDD | Train | 746 | 322 | 424 | 43.16 |
| | | Valid | 93 | 52 | 41 | 55.91 |
| | | Test | 94 | 54 | 40 | 57.45 |
| | mouse carcinogenicity | Train | 605 | 311 | 294 | 51.4 |
| | | Valid | 75 | 33 | 42 | 44 |
| | | Test | 77 | 27 | 50 | 35.06 |
| | AR | Train | 5015 | 373 | 4642 | 7.44 |
| | | Valid | 626 | 73 | 553 | 11.66 |
| | | Test | 628 | 78 | 550 | 12.42 |

**Table S17.** Dataset statistics for regression tasks in ADMET prediction.

| Type | Task | Dataset | No. drugs | Range |
|---|---|---|---|---|
| A | pKa | Train | 2175 | [-7.15, 18.46] |
| | | Valid | 271 | [-1.70, 16.00] |
| | | Test | 273 | [-1.62, 14.56] |
| | Caco-2 | Train | 1400 | [-7.70, -3.82] |
| | | Valid | 175 | [-7.40, -4.08] |
| | | Test | 175 | [-7.68, -3.78] |
| E | CLp | Train | 928 | [-2.94, 2.40] |
| | | Valid | 116 | [-2.15, 2.00] |
| | | Test | 117 | [-3.03, 1.05] |
| | MRT | Train | 932 | [-3.11, 2.00] |
| | | Valid | 116 | [-3.15, 1.30] |
| | | Test | 117 | [-2.78, 1.22] |
| T | AOT | Train | 10794 | [-5.59, 1.38] |
| | | Valid | 1349 | [-4.84, 0.72] |
| | | Test | 1350 | [-4.84, 1.30] |

**Table S18.** Dataset statistics for drug-drug interaction prediction tasks.

| Type | Dataset | No. pairs | No. drugs |
|---|---|---|---|
| Classification | Train | 213104 | 1117 |
| | Unseen one drug | 114614 | 1278 |
| | Unseen two drugs | 15318 | 264 |
| Regression | Train | 3189 | 786 |
| | Valid | 798 | 487 |
| | Test | 47 | 39 |

**Table S19**. Verification of algorithmic motivation and implementation consistency for evolved models in protein druggable site annotation.

| Method | Judgement | Notes |
| --- | --- | --- |
| ALLSites-Evo-case1 | Follow | The model implements the main processing stages specified by the method: modality-adaptive projection of protein and local descriptors, gated convolution for sequence-context encoding, cross-attention for integrating the two feature streams, and a consensus decoder for final classification. These components preserve the method's intended adaptive contact-context and consensus-prediction workflow. |
| ALLSites-Evo-case2 | Follow | The implementation contains a modality-aware physicochemical hypergraph encoder (MAPHEncoder), which groups and propagates residue information through learned higher-order relations, followed by a calibrated transport-style decoder (CSOTDecoder) that converts encoded residue features into predictions. The encoder-decoder organization is consistent with the proposed modelling strategy. |
| ALLSites-Evo-case3 | Follow | The model derives a pseudo-contact map from sequence embeddings, uses this inferred connectivity in two hypergraph-convolution layers, and combines the resulting context with cross-modal prototype information before prediction. Thus, the code retains the central design principle of exploiting sequence-derived structural neighborhoods for binding inference. |
| ALLSites-Evo-case4 | Follow | The encoder calculates residue-level precision weights and uses gated contextual encoding to form pseudo-orbital sequence features; these features are passed to a universal binding classifier. This directly reflects the proposed selective context encoding and shared prediction framework. |
| ALLSites-Evo-case5 | Follow | The implementation explicitly computes multi-scale physicochemical gradient features, integrates them in an adaptive gradient encoder, and uses a confidence-calibrated decoder that outputs prediction confidence and a binding budget. The associated confidence and budget objectives are |

| | | |
|---|---|---|
| | | also included during training, consistent with the stated sparse, confidence-aware design. |
| ALLSites-Evo-case6 | Follow | The GeodesicEncoder transforms sequence features using a soft geodesic-inspired context operator before classification. This supplies the intended graph-distance-like prior for relating residues beyond immediate sequence adjacency, while retaining a computationally compact implementation. |
| ALLSites-Evo-case7 | Follow | The encoder constructs sequence-derived features intended to represent latent three-dimensional neighborhoods and forwards them to a dedicated binding-site prediction head. The implementation therefore preserves the method's central sequence-to-pseudo-structure-to-prediction workflow. |
| ALLSites-Evo-case8 | Follow | The code provides dedicated modules for rotary positional encoding, protein-only pocket hallucination, multi-scale sequence-context extraction, decoding, and consensus prediction. This modular pipeline implements the proposed sequence-derived pocket representation and consensus-based prediction strategy. |
| ALLSites-Evo-case9 | Partial Follow | The implementation includes multi-scale convolutional sequence encoding, gated feature refinement, and attention pooling, which support the general aim of adaptive context modelling. However, the code does not explicitly instantiate the specified reliability-weighted contact graph. |
| ALLSites-Evo-case10 | Partial Follow | The implementation uses gated convolutional blocks and a protein encoder to learn sequence representations, preserving the initial feature-extraction stage. In contrast, the precision-recall balancing terms, and calibration losses are not explicitly represented and are replaced by weighted cross-entropy training. |

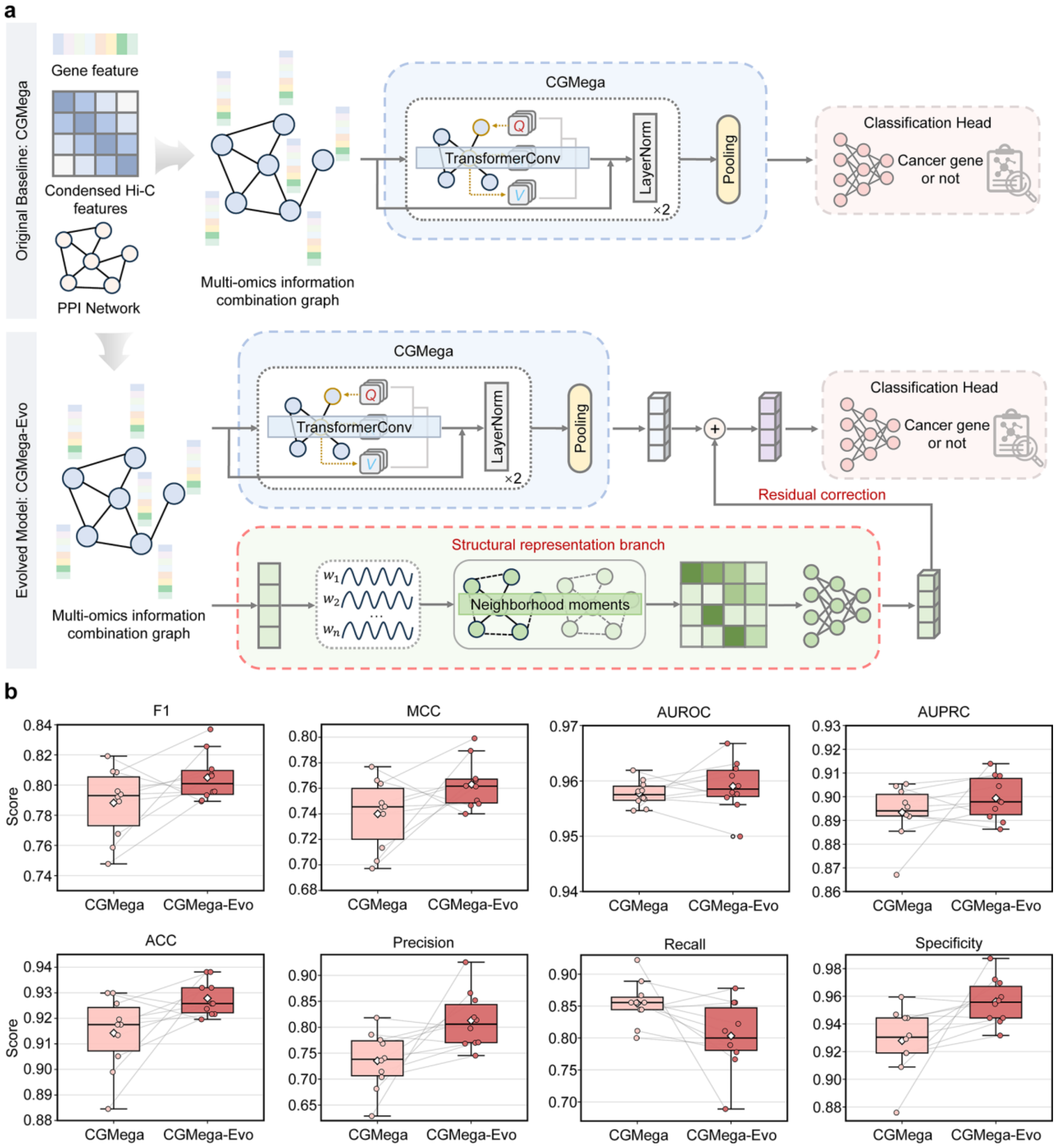

**Figure S1**. Autonomous evolution of CGMega for cancer gene module detection. **a.** Comparison of the original CGMega and evolved CGMega-Evo architectures. CGMega-Evo retains the Transformer backbone while introducing a structural representation branch and a residual correction strategy. **b.** Performance comparison of CGMega and CGMega-Evo across 10-fold evaluations on the MCF-7 breast cancer multi-omics dataset. Box plots show fold-level performance distributions, with individual points representing results from each fold and connecting lines indicating matched folds between the two models. Central lines indicate medians, white diamonds indicate means, boxes indicate the 25th-75th percentiles, and whiskers extend to values within 1.5 × interquartile range.

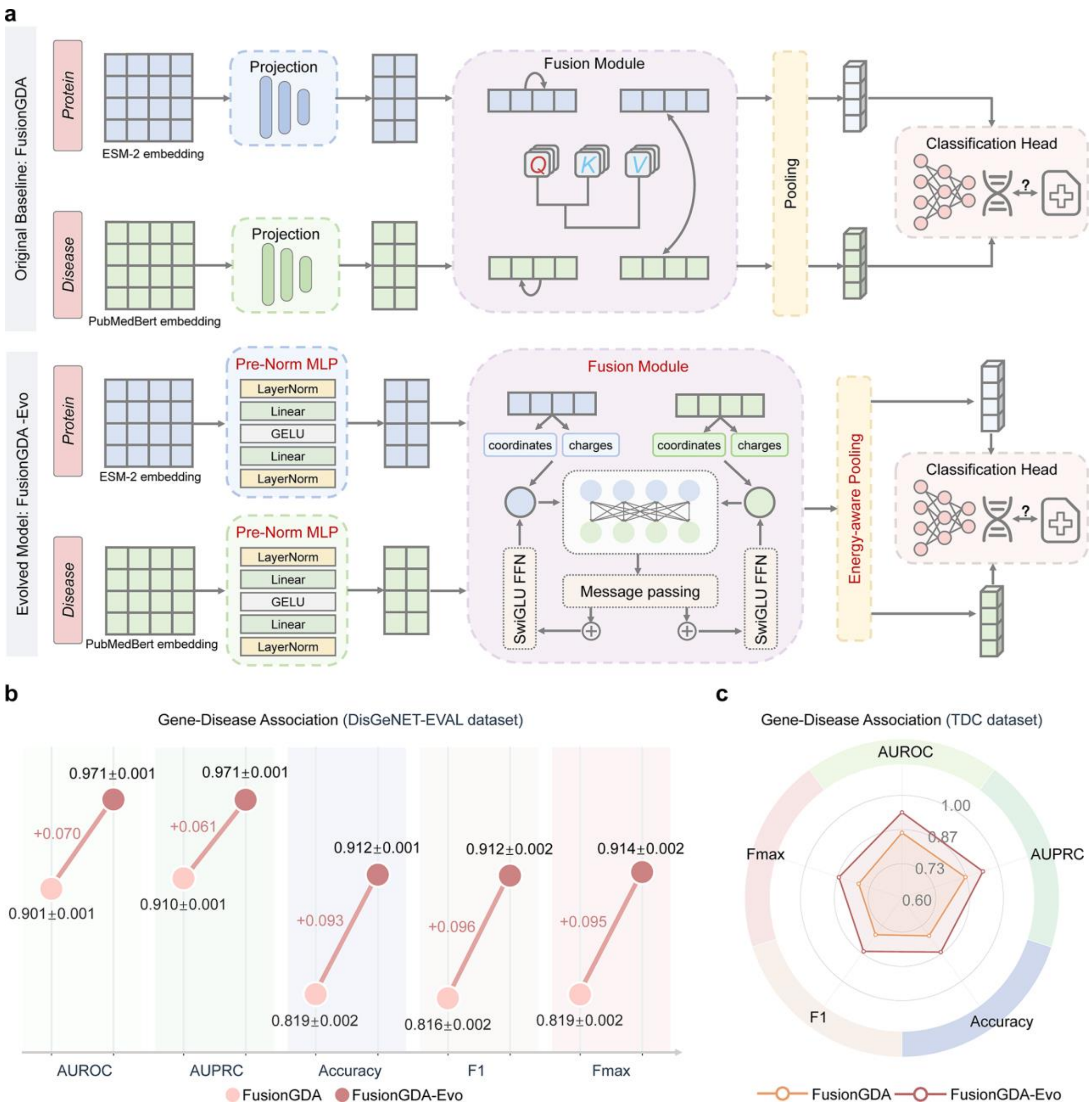

**Figure S2**. Autonomous evolution of FusionGDA for gene-disease association prediction. **a.** Comparison of the original FusionGDA and evolved FusionGDA-Evo architectures. FusionGDA-Evo replaces the original projection layer with a pre-normalized multilayer perceptron and redesigns multimodal fusion using a Hamiltonian interaction framework. **b.** Performance comparison between FusionGDA and FusionGDA-Evo on the DisGeNET-EVAL benchmark, evaluated by AUROC, AUPRC, Accuracy, F1, and Fmax. Values represent mean ± standard deviation across five independent random splits. **c.** Performance comparison between FusionGDA and FusionGDA-Evo on the TDC benchmark, evaluated by AUROC, AUPRC, Accuracy, F1, and Fmax.

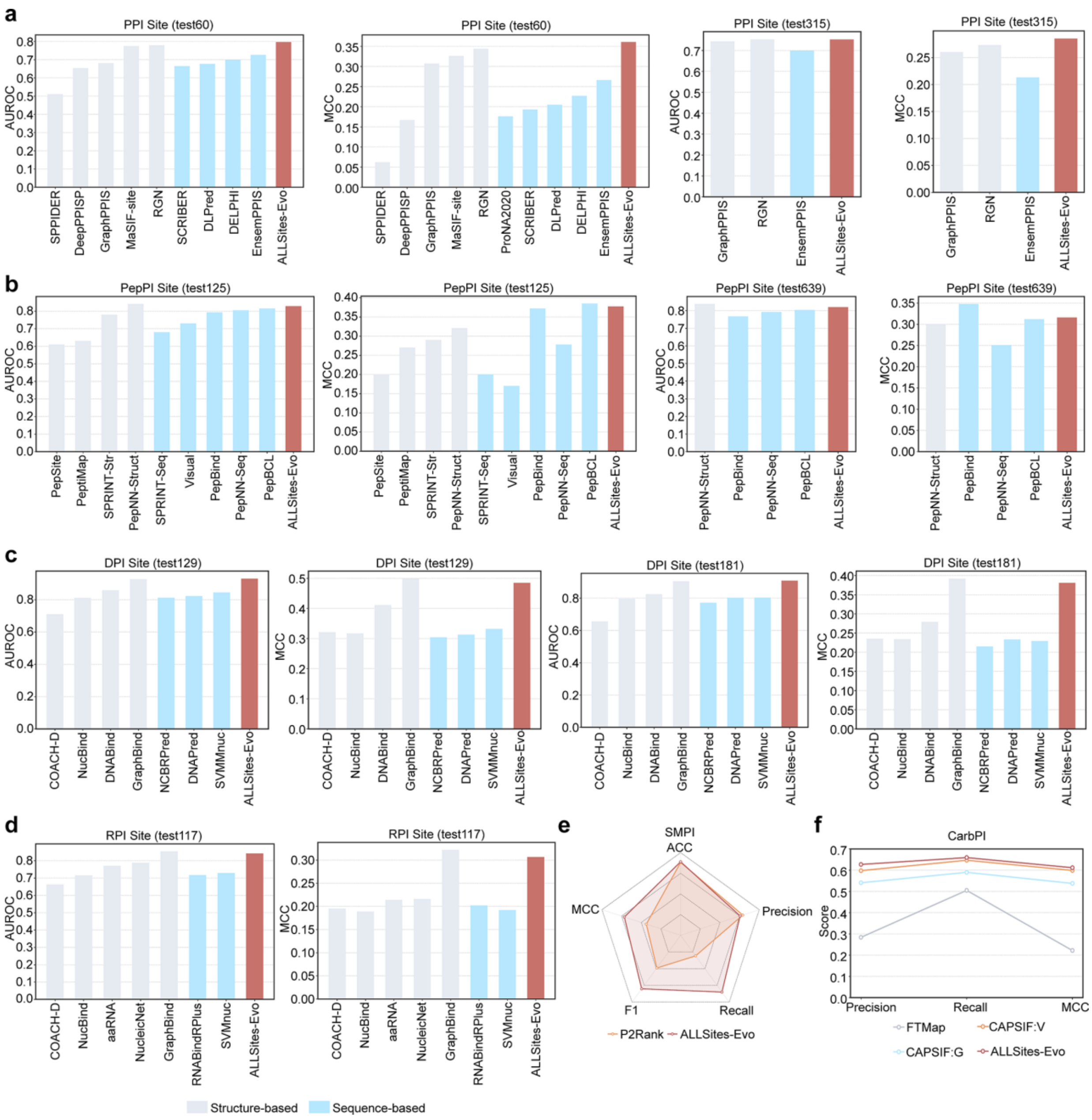

**Figure S3**. Performance comparison of ALLSites-Evo with task-specific methods for protein druggable residue annotation. Structure-based methods are shown in gray and sequence-based methods in blue. **a.** Performance comparison on PPI site prediction using the PPI-Test60 and PPI-Test315 datasets, evaluated by AUROC and MCC. **b.** Performance comparison on PepPI site prediction using the PepPI-Test125 and PepPI-Test639 datasets, evaluated by AUROC and MCC. **c.** Performance comparison on DNA-binding site prediction using the DPI-Test129 and DPI-Test181 datasets, evaluated by AUROC and MCC. **d.** Performance comparison on RNA-binding site prediction using the RPI-Test117 dataset, evaluated by AUROC and MCC. **e.** Performance comparison on small molecule-binding site prediction using the SMPI dataset, shown as radar plots of ACC, Precision, Recall, F1, and MCC. **f.** Performance comparison on carbohydrate-binding site prediction using the CarbPI dataset, evaluated by Precision, Recall, and MCC.

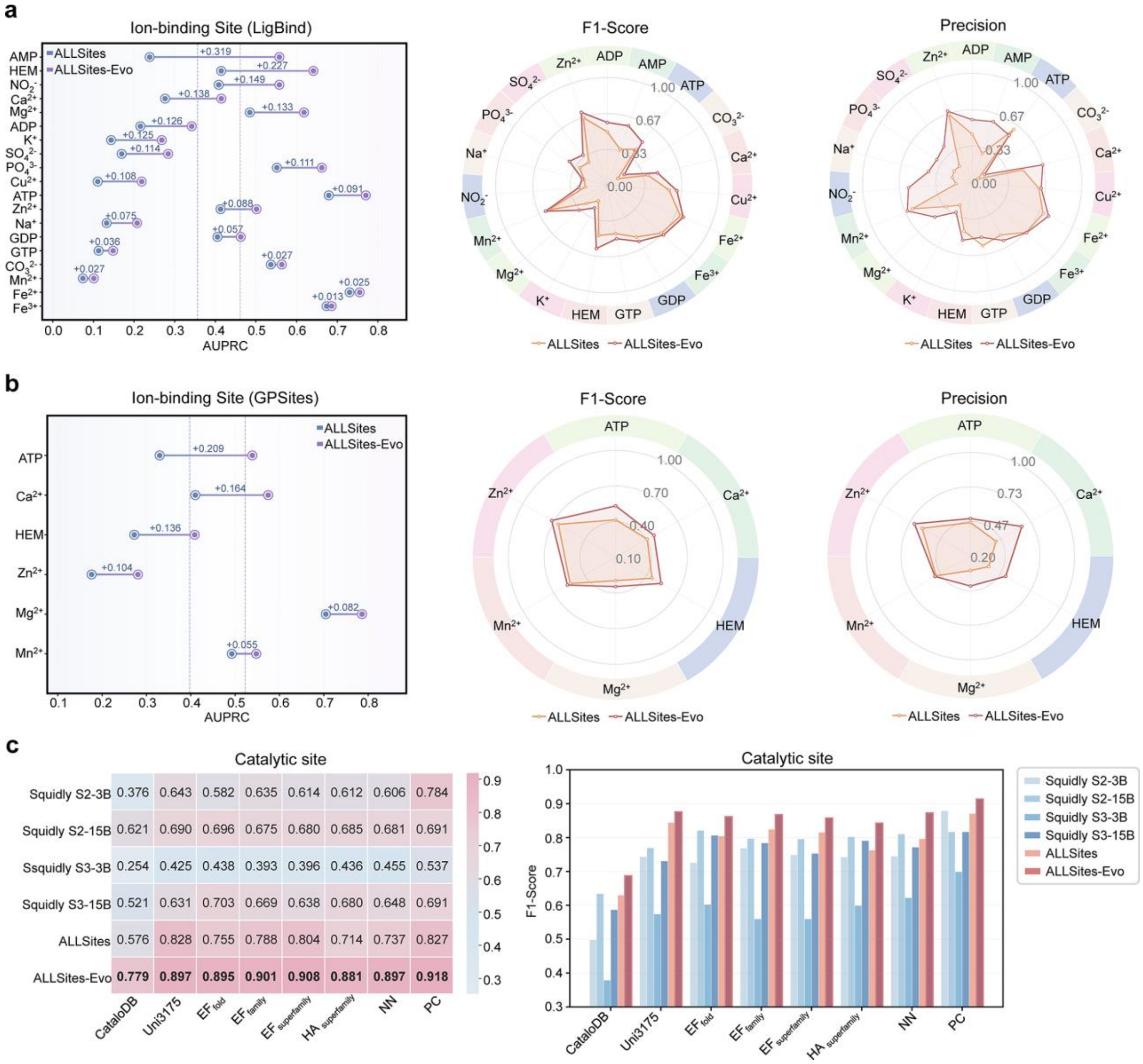

**Figure S4**. Detailed performance comparisons for unseen protein druggable residue annotation tasks. **a.** Performance comparison between ALLSites and ALLSites-Evo for ion-binding site prediction on the LigBind dataset, evaluated by AUPRC (left), F1-score (middle), and precision (right) across different ion and ligand types. **b.** Performance comparison between ALLSites and ALLSites-Evo for ion-binding site prediction on the GPSites dataset, evaluated by AUPRC (left), F1-score (middle), and precision (right) across different ion and ligand types. **c.** Performance comparison of ALLSites and ALLSites-Evo against reproduced Squidly variants across catalytic site benchmarks, including CataloDB, Uni3175, and six additional benchmark datasets. Results are evaluated by precision (left) and F1-score (right).

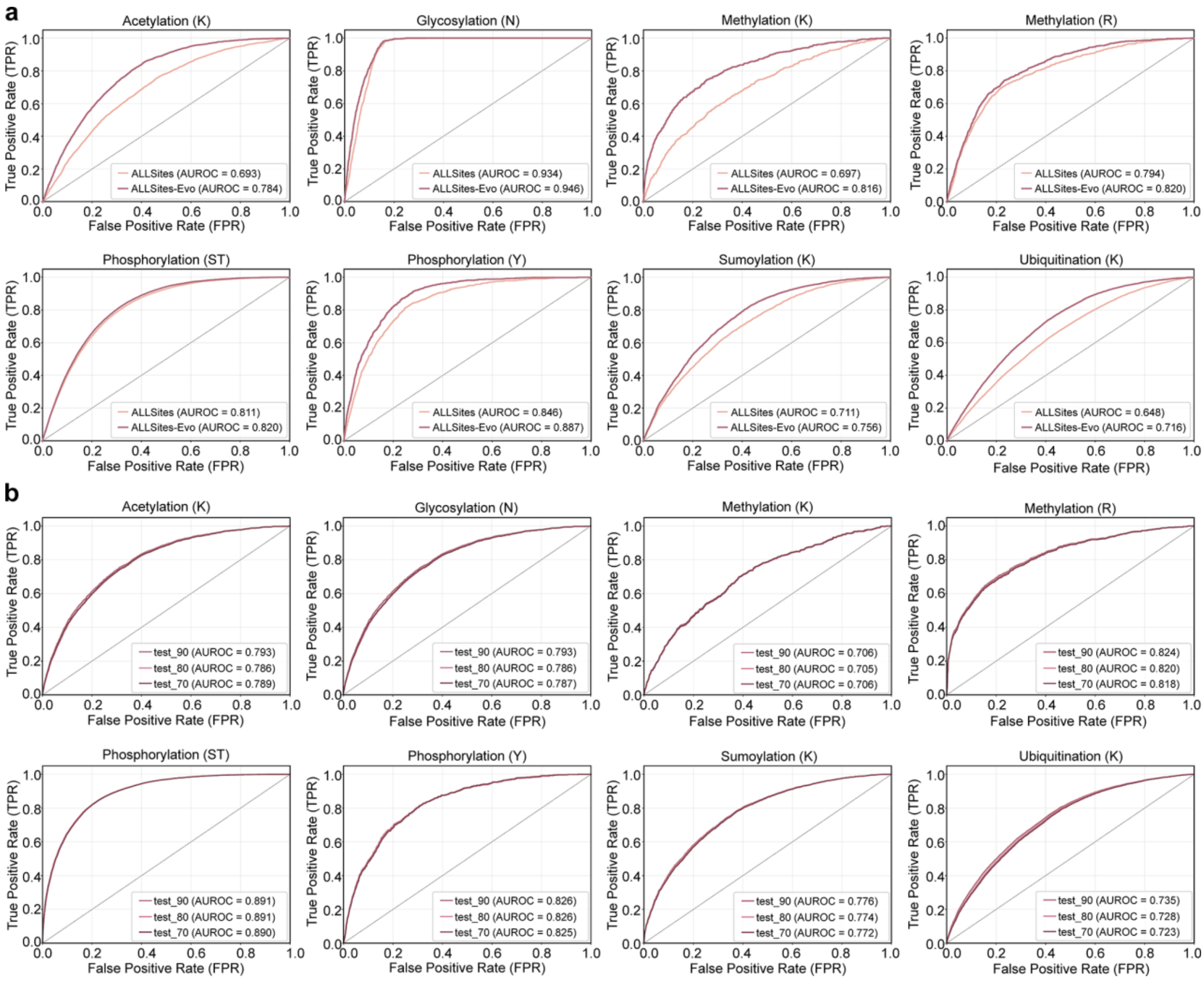

**Figure S5**. Performance of ALLSites-Evo on post-translational modification (PTM) site annotation. **a.** Performance comparison between ALLSites and ALLSites-Evo for PTM site prediction across eight benchmark datasets spanning six PTM types, evaluated by AUROC. **b.** Performance evaluation of ALLSites-Evo for PTM site prediction using datasets constructed under three protein-level sequence similarity thresholds (70%, 80%, and 90%), evaluated by AUROC.

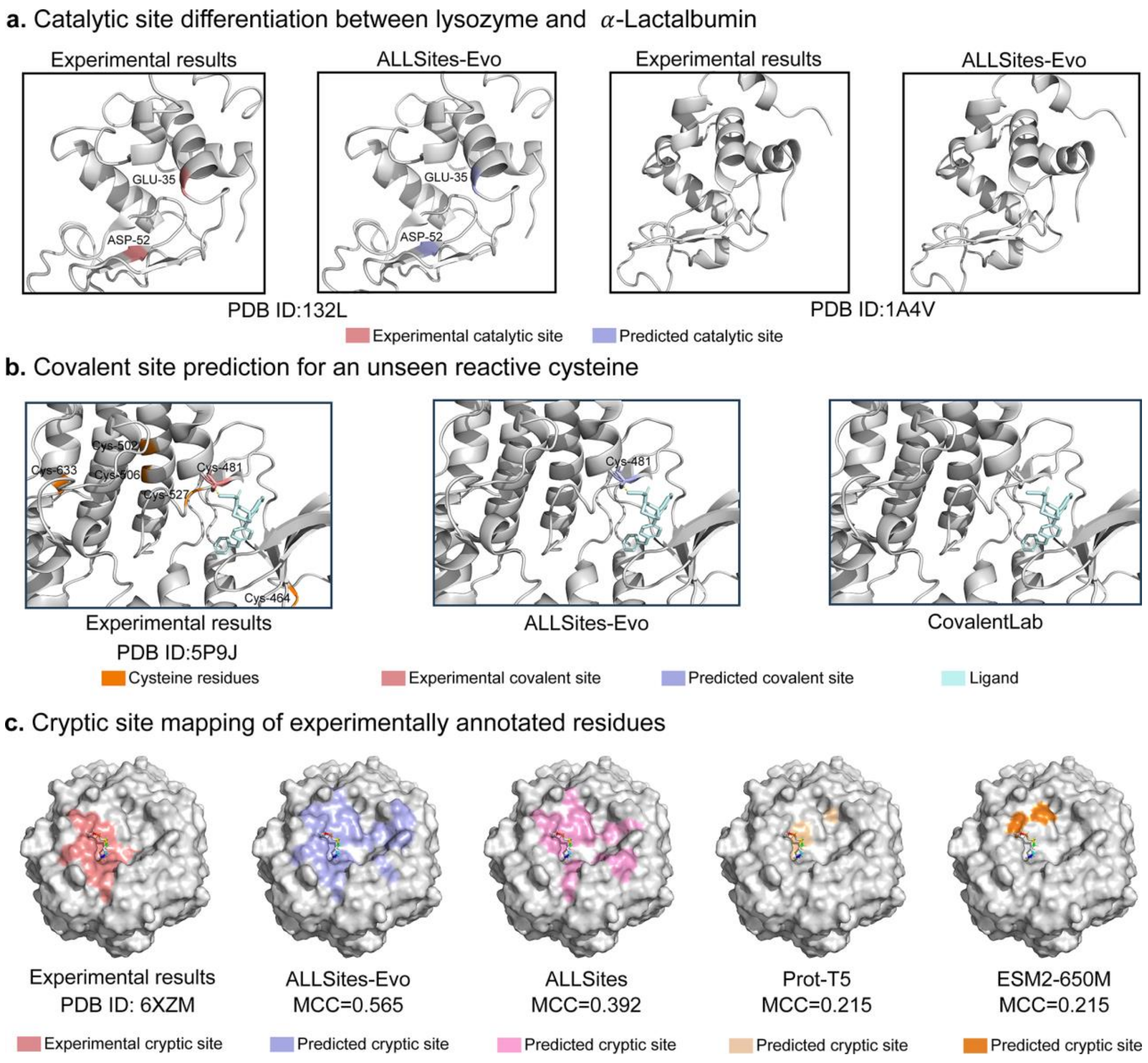

**Figure S6**. Case studies of ALLSites-Evo predictions on diverse druggable residue types. **a.** Catalytic site annotation comparison between lysozyme (PDB ID: 132L) and α-lactalbumin (PDB ID: 1A4V), comparing experimentally annotated sites with ALLSites-Evo predictions. Red and purple indicate experimental and predicted catalytic sites, respectively. **b.** Prediction of an unseen covalent site involving a reactive cysteine (PDB ID: 5P9J), comparing experimental annotations with ALLSites-Evo and CovalentLab predictions. Orange indicates all cysteine residues, red indicates the experimental covalent site, purple indicates the predicted covalent site, and blue indicates the ligand. **c.** Cryptic site mapping of experimentally annotated residues, comparing predictions from ALLSites-Evo, ALLSites, fine-tuned Prot-T5, and fine-tuned ESM-2-650M. Colors indicate experimental sites and predictions from different models.

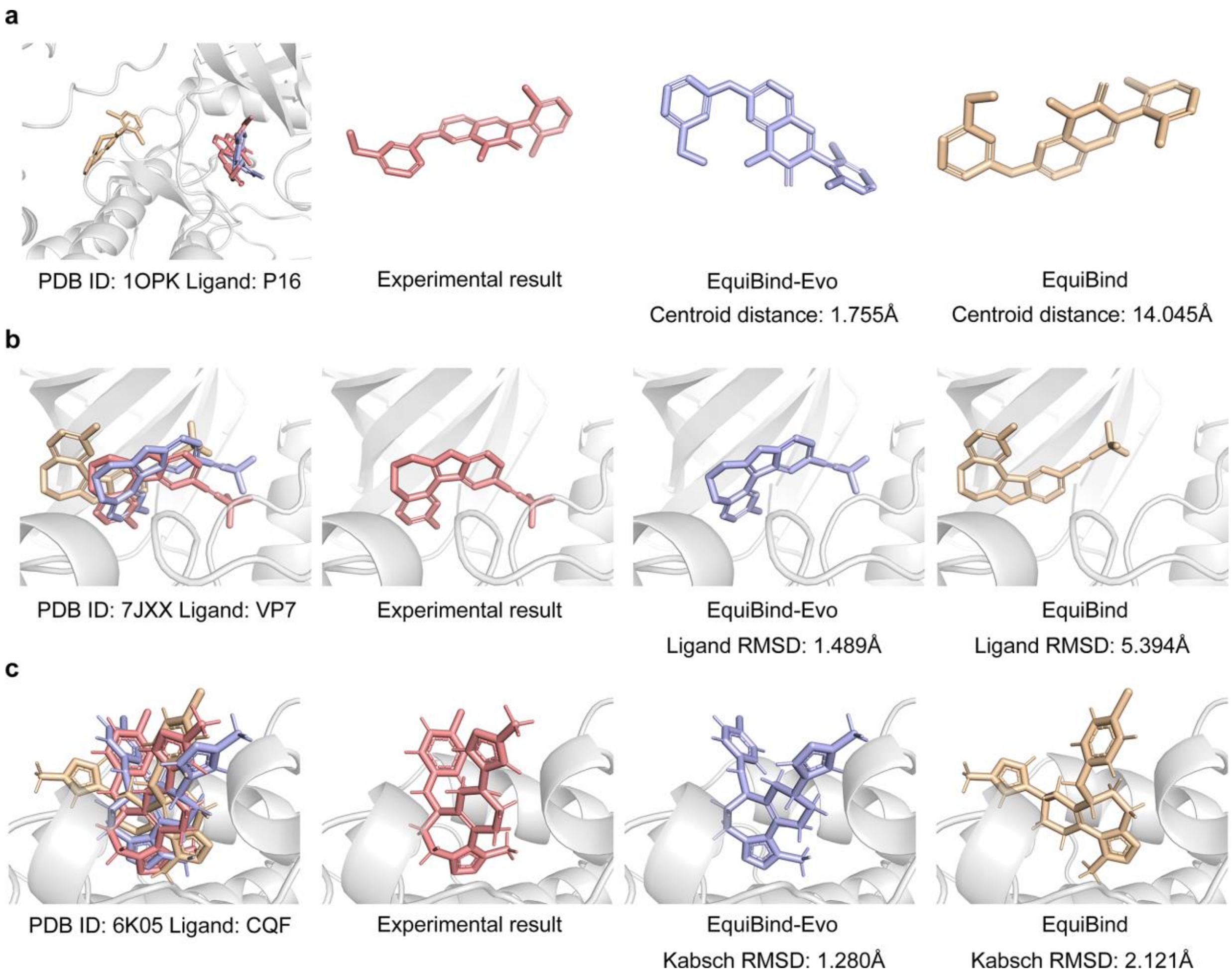

**Figure S7**. Representative case studies illustrating geometric improvements achieved by EquiBind-Evo. **a.** Recovery of the crystallographic binding pocket by EquiBind-Evo on the Astex Diverse Set (PDB ID: 1OPK; ligand: P16). **b.** Improved ligand placement while preserving local conformation on the PoseBusters benchmark (PDB ID: 7JXX; ligand: VP7). **c.** Enhanced ligand conformational refinement on the PDBbind test set (PDB ID: 6K05; ligand: CQF).

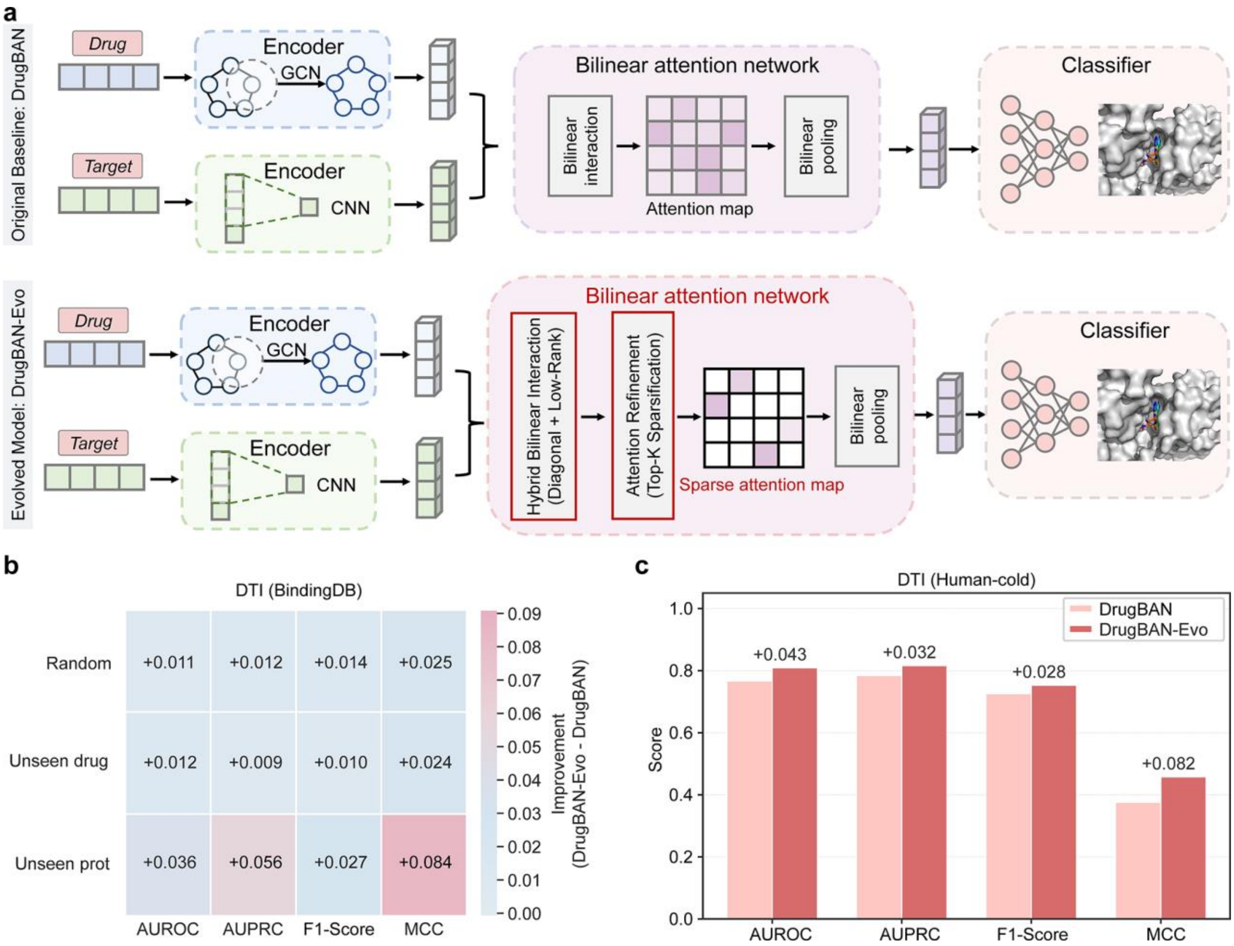

**Figure S8**. Autonomous evolution of DrugBAN for drug-target interaction prediction. **a.** Comparison of the original DrugBAN and evolved DrugBAN-Evo architectures. DrugBAN-Evo redesigns cross-modal interaction modeling by introducing a hybrid bilinear interaction module with Top-K sparse attention. **b.** Performance improvement of DrugBAN-Evo over DrugBAN on the BindingDB benchmark under random, unseen-drug, and unseen-protein splits. Performance differences (DrugBAN-Evo - DrugBAN) are shown for AUROC, AUPRC, F1-score, and MCC. **c.** Performance comparison between DrugBAN and DrugBAN-Evo on the Human-cold dataset, evaluated by AUROC, AUPRC, F1-score, and MCC.

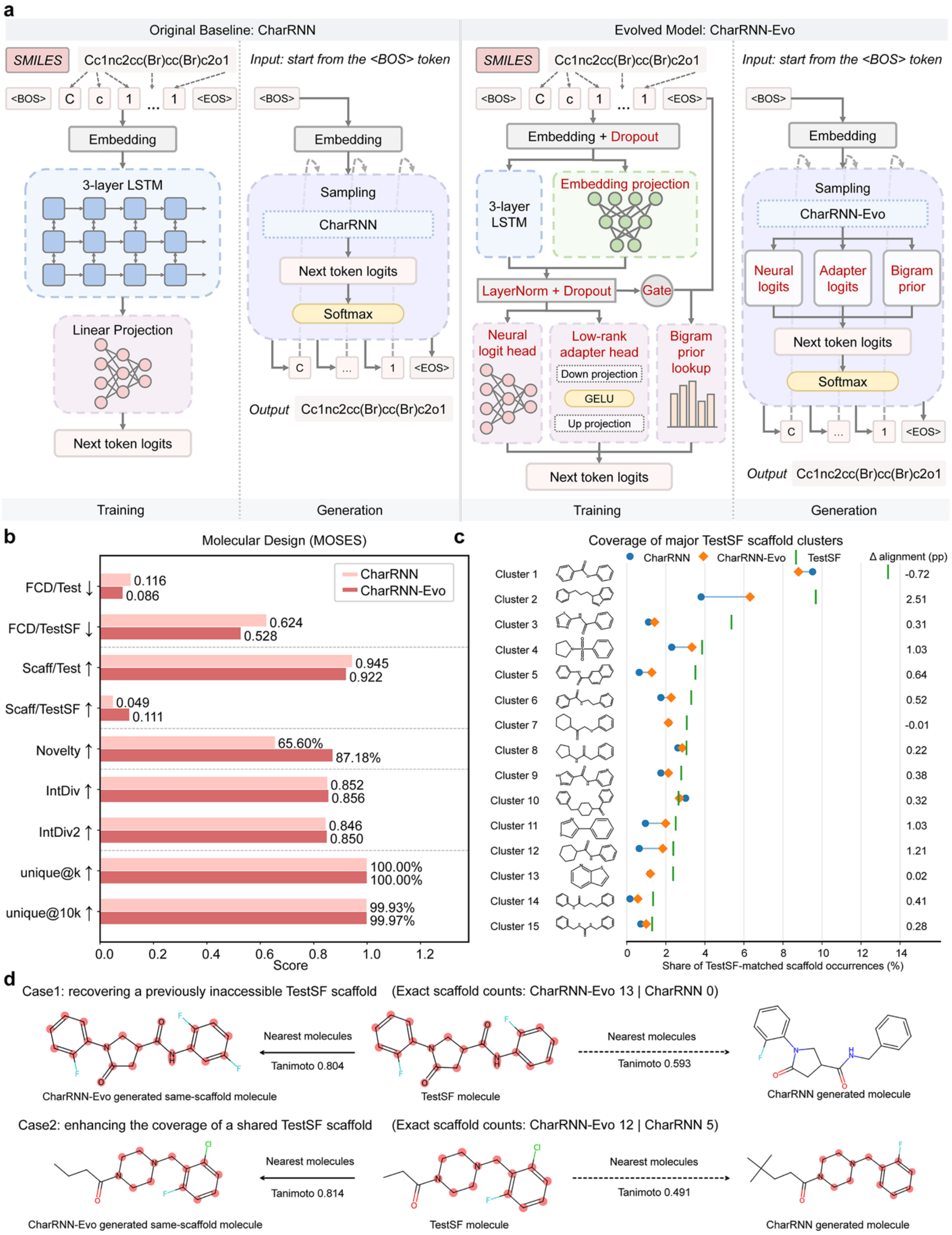

**Figure S9**. Autonomous evolution of CharRNN for de novo molecular generation. **a.** Comparison of the original CharRNN and evolved CharRNN-Evo architectures. CharRNN-Evo redesigns sequence representation and next-token prediction by integrating residual embedding propagation with neural, adapter, and gated statistical prediction heads. **b.** Performance comparison of CharRNN and CharRNN-Evo on the MOSES benchmark using molecular distribution, diversity, and novelty metrics. Results are shown for the random test set (Test) and scaffold-based test set (TestSF), which evaluate generalization under conventional and unseen-scaffold settings, respectively. **c.** Scaffold distribution across major TestSF-derived scaffold clusters. Major scaffold clusters were defined by clustering Bemis-Murcko scaffolds from the MOSES TestSF set using Morgan fingerprint similarity and Butina clustering. The relative abundance of molecules generated by CharRNN and CharRNN-Evo was compared with the TestSF reference distribution across the major scaffold clusters. Positive Δ alignment values indicate that CharRNN-Evo more closely matched the TestSF scaffold distribution. **d.** Representative molecular examples illustrating expanded scaffold coverage and improved scaffold-frequency matching by CharRNN-Evo.

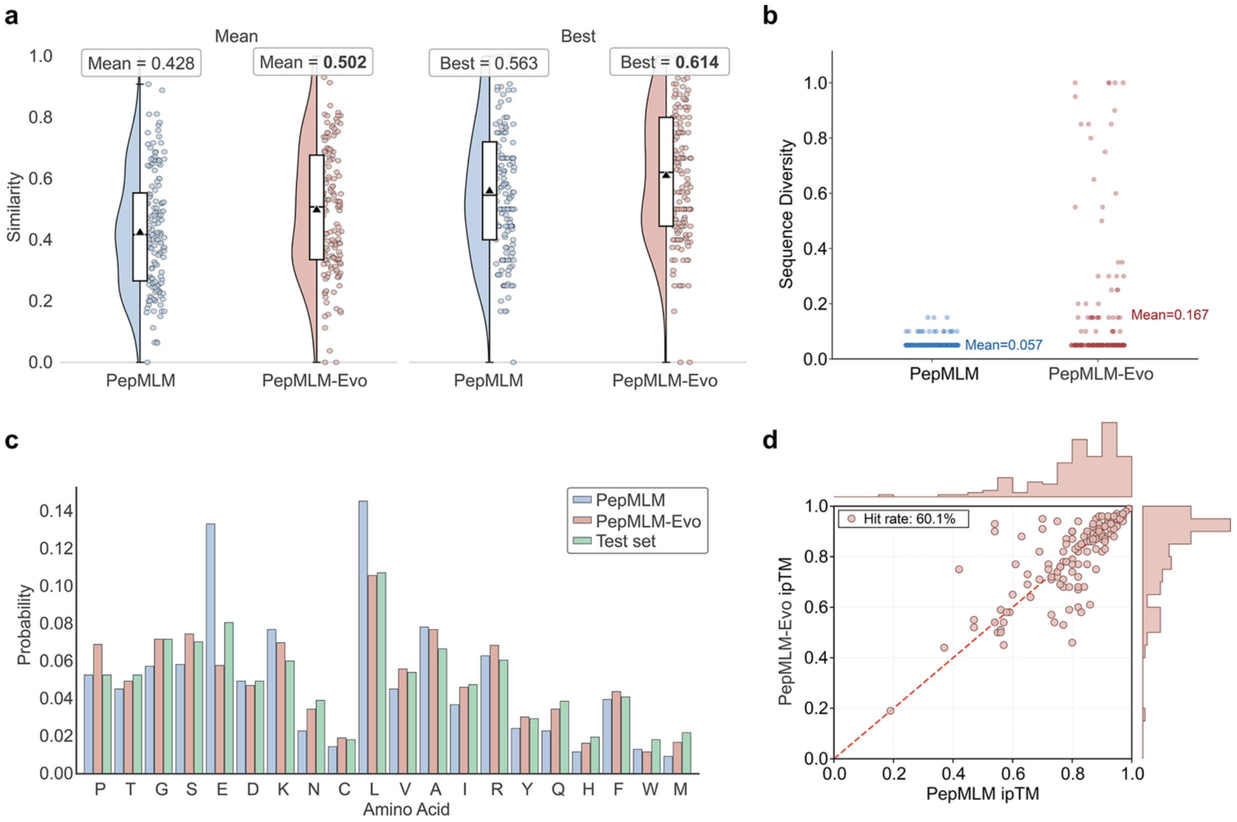

**Figure S10**. Evaluation of PepMLM-Evo and PepMLM on the PepNN benchmark. **a.** Sequence similarity of peptides generated by PepMLM and PepMLM-Evo. Violin plots show the distribution of sequence similarity values, individual dots represent generated peptides, and black triangles indicate mean values. The left groups report the overall mean similarity, whereas the right groups report the mean best similarity, calculated as the average of the highest similarity peptide for each target. **b.** Sequence diversity of peptides generated by PepMLM and PepMLM-Evo. Each point represents the diversity of generated peptides for an individual target, and horizontal lines indicate the mean diversity across targets. **c.** Amino acid frequency distributions of peptides generated by PepMLM (blue) and PepMLM-Evo (red), compared with the PepNN test set (green). **d.** In silico hit-rate evaluation using AlphaFold3-predicted ipTM scores. Each point represents one target-peptide pair. Hits are defined as target-peptide pairs for which the peptide generated by PepMLM-Evo achieves an ipTM score equal to or higher than that generated by PepMLM.

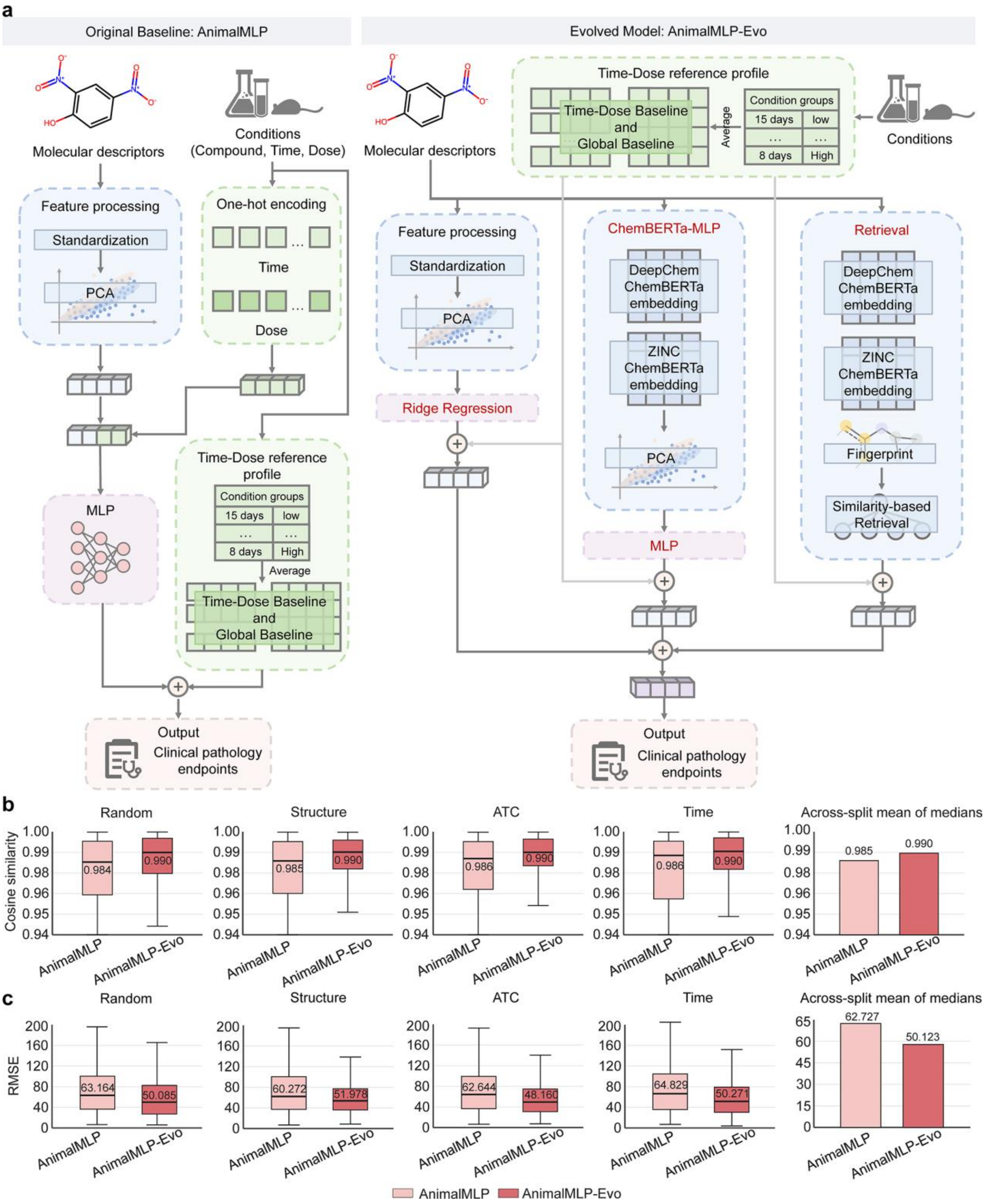

**Figure S11**. Autonomous evolution of AnimalMLP for animal toxicology assessment. **a.** Comparison of the original AnimalMLP and evolved AnimalMLP-Evo architectures. AnimalMLP-Evo enhances molecular representation and response prediction by integrating complementary molecular representations and similarity-based residual correction. **b.** Performance comparison of AnimalMLP and AnimalMLP-Evo using cosine similarity under random, structure, ATC, and temporal compound-level splits, with an across-split summary based on the mean of split-level medians. **c.** Performance comparison of AnimalMLP and

AnimalMLP-Evo using RMSE under random, structure, ATC, and temporal compound-level splits, with an across-split summary based on the mean of split-level medians. Box plots show split-level performance distributions, where central lines indicate medians, white diamonds indicate means, boxes indicate the 25th-75th percentiles, and whiskers extend to the most extreme values within 1.5 × interquartile range.

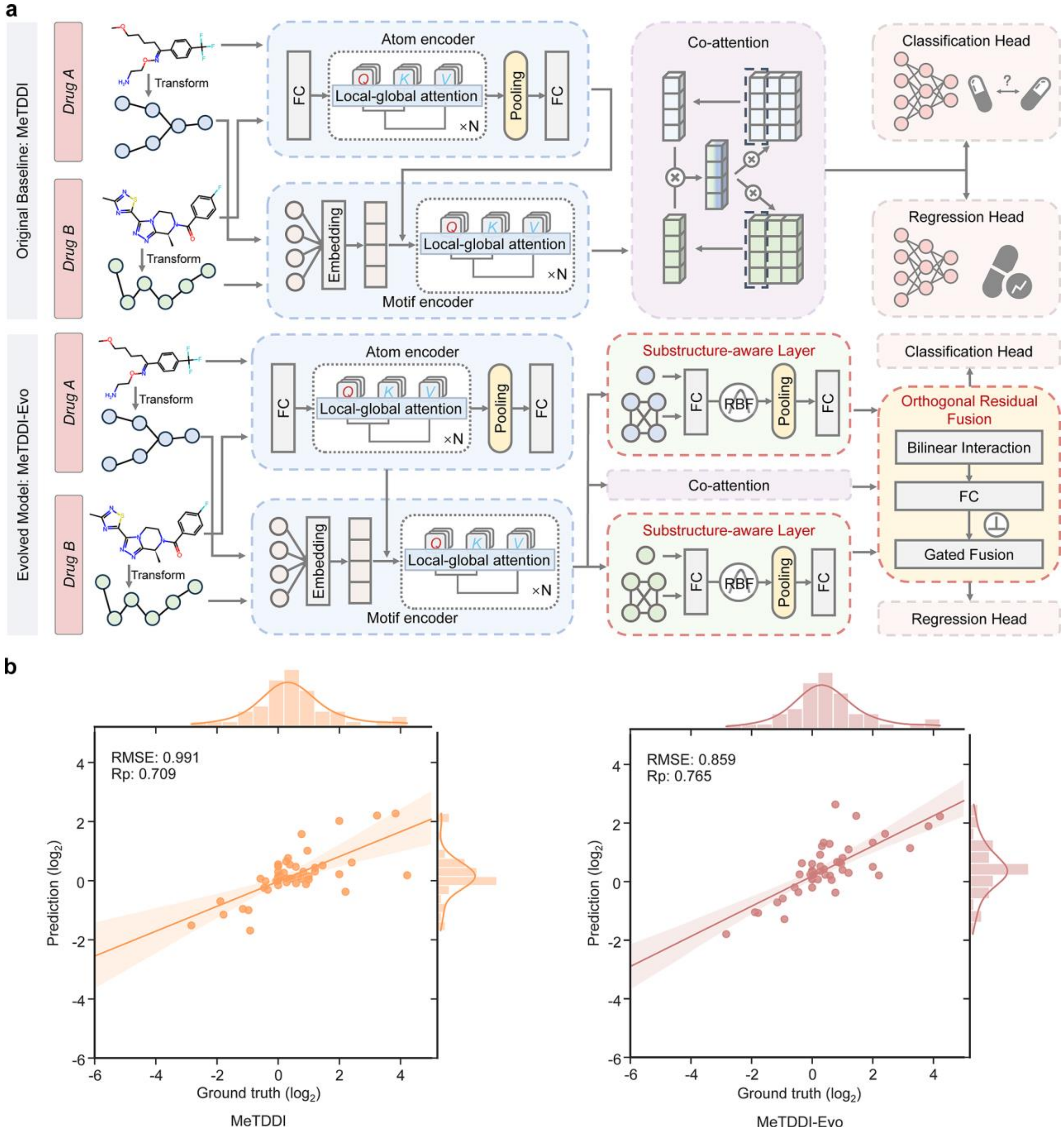

**Figure S12**. Autonomous evolution of MeTDDI for drug-drug interaction prediction. **a.** Comparison of the original MeTDDI and evolved MeTDDI-Evo architectures. MeTDDI-Evo introduces substructure-aware interaction modeling and orthogonal residual fusion to enhance pairwise drug representations. **b.** Regression performance comparison between MeTDDI and MeTDDI-Evo. Scatter plots show the relationship between predicted and experimental drug-drug interaction values, with performance evaluated using RMSE and Pearson correlation (Rp).

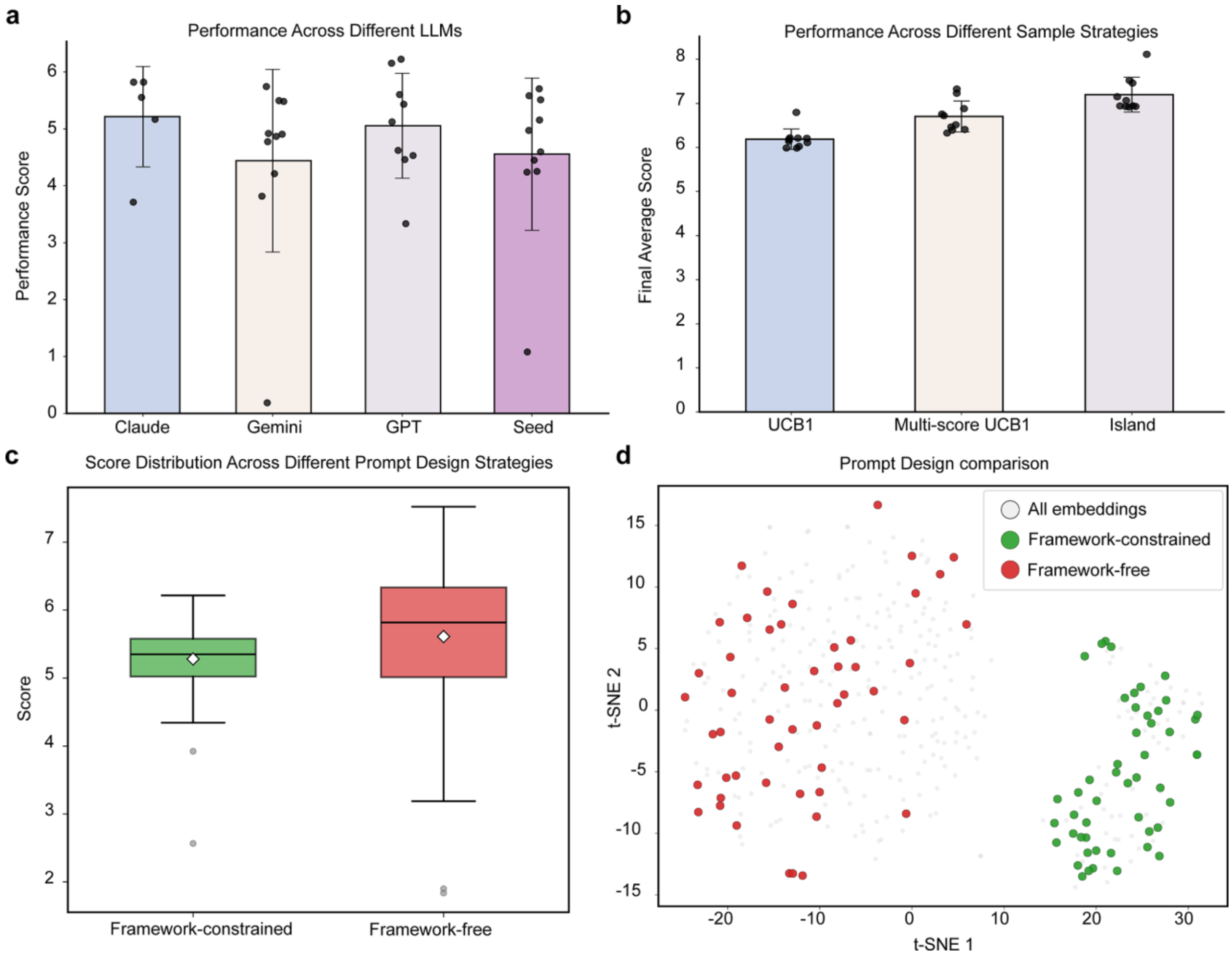

**Figure S13**. Analysis of evolutionary search configurations in DrugEvolve. **a.** Comparison of different foundation models as system-level controllers of DrugEvolve using protein druggable site annotation as a representative task. Score distributions across evolutionary runs are shown for Gemini 3.5 Flash, Claude Sonnet 4.6, GPT-5.5, and Seed 2.0 Pro, with each configuration evaluated over ten independent evolutionary runs. **b.** Comparison of evolutionary sampling strategies. UCB1-based sampling, UCB1 sampling with auxiliary task-specific scores, and island-based sampling are compared using their top-performing evolutionary trajectories. **c.** Comparison of framework-constrained and framework-free prompt designs. Score distributions are shown for the two prompting schemes. Box plots indicate the 25th-75th percentiles, central lines indicate medians, whiskers extend to values within 1.5 × the interquartile range, and white diamonds indicate mean values. **d.** t-SNE visualization of generated algorithm proposals under framework-constrained and framework-free prompt designs. Each colored point represents one design proposal (n = 50 for both settings), allowing qualitative comparison of the distributions produced by the two prompting schemes in the two-dimensional t-SNE projection.
